# Characterization and engineering of highly efficient Cas12j genome editors

**DOI:** 10.1101/2025.09.16.676532

**Authors:** Gundra Sivakrishna Rao, Wenjun Jiang, Mustapha Aouida, Qiaochu Wang, Ahmed M. Kazlak, Ali H. A. Elbehery, Ahmed Saleh, Maazallah Masood, Ahmed Ghouneimy, Magdy Mahfouz

## Abstract

The large size of CRISPR-Cas enzymes limits their delivery for therapeutic applications. Cas12j nucleases offers hypercompact alternative but show moderate editing efficiency. To overcome this limitation, we identified eight novel Cas12j orthologues (Cas12j-11 to Cas12j-18) from viral metagenomes. All showed low editing activity in mammalian cells. We engineered T5 exonuclease-Cas12j fusions (T5Exo-Cas12j), two of which, T5Exo-Cas12j-12, and -18 exhibited up to 42% editing in HEK293T and 9% in K-562 cells, outperforming wild-type Cas12j counterparts and comparable to LbCas12a. Intriguingly, robust *in cellula* editing in both HEK293T and K-562 cells was strictly dependent on the presence of 5′-TAC trinucleotides within the target DNA sequence. Furthermore, we fused the Cas12j orthologues with the TadA8e deaminase and developed base editors, termed Be-(d)Cas12j. Among these, Be-(d)Cas12j-13 demonstrated efficient A-to-G base conversion in mammalian cells. This study expands the CRISPR toolbox by characterizing and engineering novel Cas12j orthologues into compact, high-efficiency genome editors.

## Introduction

The CRISPR (Clustered Regularly Interspaced Short Palindromic Repeats) is a prokaryotic adaptive immune system that has been repurposed for genome editing applications across prokaryotic and eukaryotic organisms [1,2]. Different CRISPR types and subtypes have been characterized and optimized for gene editing applications. However, a significant focus is placed on the class II CRISPR types and subtypes, which require a single effector to target the genome for editing. For example, class II type II with a signature Cas9 protein marked the beginning of CRISPR for gene editing and its adoption for biotechnological applications [3]. Class II type V with Cas12 as a signature protein has been harnessed for genome editing applications and molecular diagnostics. Although Cas9 and Cas12 exhibit robust gene editing activities, they suffer from key application challenges, including the delivery of the CRISPR cargo into target cells and their on-target specificity in the genome compared with other potential off-targets, complicating their usefulness for translational applications [4]. For example, type II SpyCas9 or type V Cas12a are widely used in genome editing for functional analysis [2]. However, their large gene sizes pose a challenge for packaging into viral vectors, which can hinder their delivery and thus limit their utility in diverse applications, including gene therapy. To address this challenge, efforts have been dedicated to discovering miniature Cas effectors to enable efficient delivery into diverse cell types. Class II type V enzymes have been quite promising in addressing the delivery challenge as miniature Cas12 enzymes have been identified and characterized, including enzymes such as Cas12k, Cas12m, Cas12f, and Cas12j [5–7]. To address AAV’s limited packaging capacity, several strategies beyond compact enzyme design have been developed. These include split-Cas systems with intein reconstitution, use of minimal promoters, and non-viral delivery platforms such as lipid nanoparticles [8–11]. Together, these approaches expand the delivery options for genome editing *in vivo*.

Cas12k and Cas12m don’t possess nuclease catalytic activity [7]. They could be harnessed as a reprogrammable DNA binding module and fused with functional domains such as transposon or deaminase for gene editing applications. Due to the small size (400-800 AA), Cas12f remains the most miniature Cas12 family member [6,12]. Certain Cas12f variants can specifically recognize and cut double-stranded DNA (dsDNA) when a protospacer adjacent motif (PAM) is present, making them highly promising for gene editing applications [13]. Cas12f performs targeted cleavage of dsDNA, directed by crRNA and dependent on a 5′-TTTR-3′ PAM sequence, where “R” stands for either adenine (A) or guanine (G) [14]. The PAM sequence of Cas12f constrains its target selection across various genomes [15]. Pausch et al., [16] discovered a novel family of type V CRISPR-Cas proteins termed CasΦ (Cas12j). Compared with other compact Cas enzymes, Cas12j nucleases have relatively small ribonucleoprotein complexes (RNP) due to small protein and the lack of trans-activating CRISPR RNA (tracrRNA). Cas12j enzymes have been shown to mediate genome editing in mammalian cells, expanding the genome editing toolkit. Cas12j recognizes and unwinds double-stranded DNA by PAM sequences 5′-TBN-3′ (where B can be G, T, or C) located adjacent to the crRNA-complementary target DNA strand, which is more flexible than Cas12f. However, only Cas12j-2, Cas12j-8, and Cas12j-SF05 displayed modest editing efficiencies in mammalian cells, thereby limiting their translational applications [17,18].

In this study, we conducted a comprehensive search of viral genome databases [19], identifying eight previously uncharacterized Cas12j orthologues (Cas12j-11 to Cas12j-18). While all eight orthologues exhibited low genome editing activity in mammalian cells, similar to the previously characterized Cas12j-8, the small size of Cas12j proteins presents a unique opportunity for engineering improved variants to enhance efficiency. To address the challenges associated with their limited catalytic activity, we designed, built, and tested chimeric fusions between Cas12j orthologues and T5 exonuclease (T5Exo) and TadA8e deaminase enzymes, independently. Furthermore, we conducted exhaustive analysis and determined a 5′-TAC trinucleotide sequence requirement in the target for robust *in cellula* catalytic activity of Cas12j variants. T5Exo-Cas12j-12, -13 and -18 exhibit robust genome editing, and Be-(d)Cas12j-13 showed robust base conversion efficiency, providing compact alternatives to overcome current delivery limitations, maximize targetability, and expand the genome editing toolkit for diverse biotechnological applications, including gene therapy.

## Results

### Discovery of previously uncharacterized Cas12j orthologues from viral metagenomes

Application of the HMM-based search strategy to the high-confidence subset of IMG/VR v4.1 yielded four initially identified Cas12j protein candidates. These discoveries prompted refinement of the Cas12j alignment and HMM models, which were subsequently used to scan the comprehensive IMG/VR v4.1 repository of 15,722,824 viral genomes. This iterative approach led to the discovery of four additional Cas12j putative orthologues, bringing the total to eight. All candidates exhibited a maximum E-value of 1e-10, were positioned within 1,000 bp of a neighboring CRISPR array, and fell within the characteristic length range (700–800 AA) (Fig. 1A). Clustering analysis at 90% identity confirmed that these eight sequences were non-redundant. Validation by CRISPRCasTyper further revealed the presence of compact cassettes encoding Cas12j along with CRISPR arrays, consistent with the canonical single-effector CRISPR-Cas architecture.

**Figure 1.**
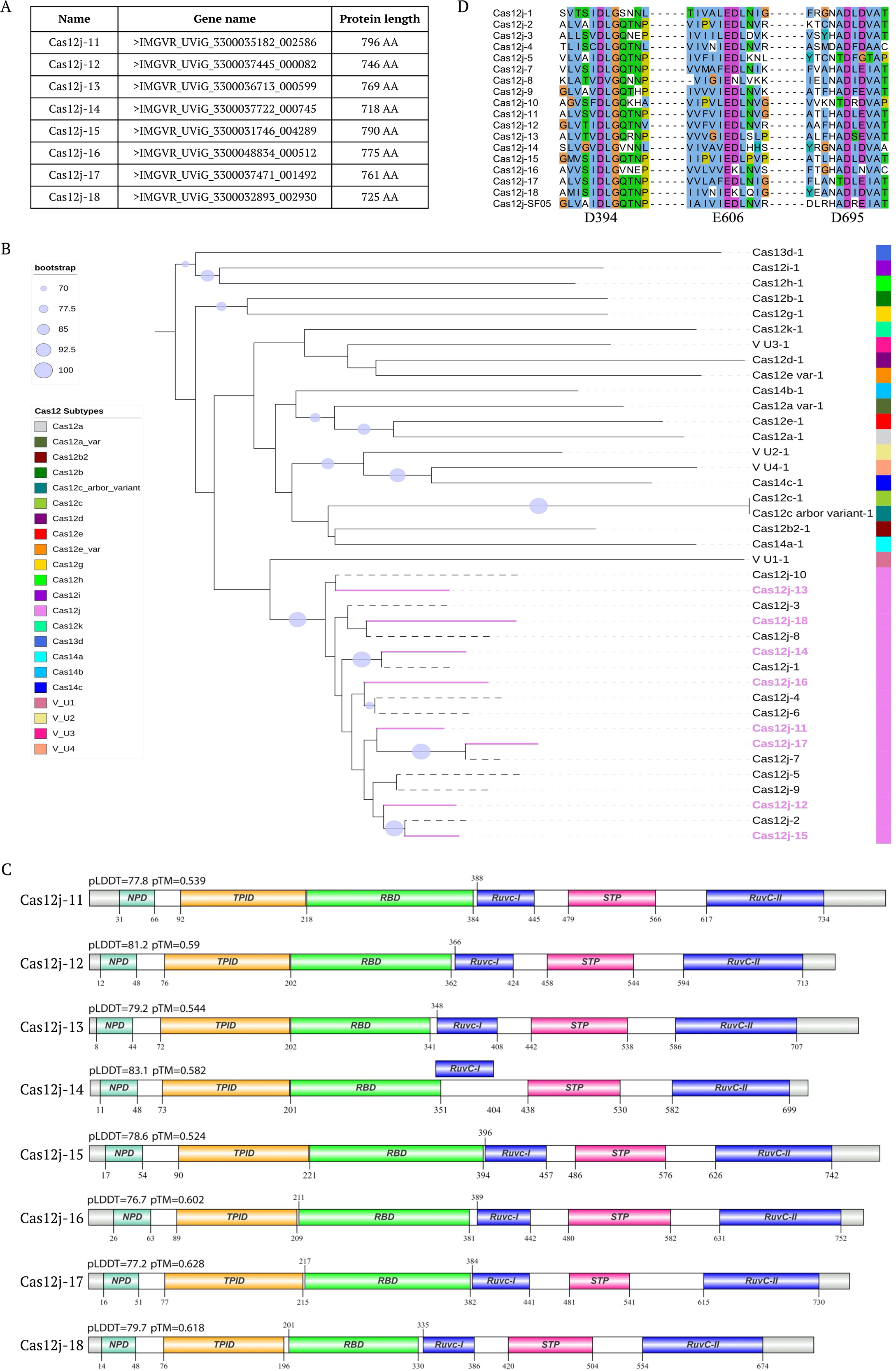
Discovery of novel Cas12j orthologues from viral metagenomes. **(A)** The size and contig source of eight Cas12j candidates identified. **(B)** The phylogenetic tree shows the evolutionary relationships among various Cas protein types, featuring one representative from each major Cas subtype. The newly identified Cas12j variants are highlighted in purple, while previously identified variants of the same family are represented by dashed lines. Circles represent bootstrapping, with only scores above 70 displayed. The tree is rooted at the midpoint. **(C)** Domain architecture of eight newly identified Cas12j candidates mapped onto their full-length protein sequences (gray backbone, relative to the protein length). Domains are color-coded for clarity: NPD (sea foam), TPID (yellow), RBD (green), RuvC-I, RuvC-II (blue), and STP (pink). The predicted local distance difference test (pLDDT) and predicted TM-score (pTM) for each protein structure were generated using AlphaFold2. Cas12j-14 has a RuvC-I domain that overlaps with the RBD domain in 6 AAs highlighted by displacing the domain above the sequence line. **(D)** Multiple sequence alignment of predicted versus reference Cas12j proteins showing conserved active site residues. Only selected regions of the alignment are shown; dashed lines indicate breaks where intervening sequences have been omitted.

Multiple sequence alignment indicated that these novel Cas12j proteins share 40-50% sequence identity with reference Cas12j variants, and phylogenetic analysis shows their interspersing among previously identified Cas12j proteins, supporting their designation as bona fide Cas12j orthologues (Fig. 1B). Protein structural predictions performed using AlphaFold2 identified highly conserved domain architectures, containing T-strand and NT-strand PAM interacting domains (TPID, NPD), the RNA-handle binding domain (RBD), the bridge helices (BH-I and BH-II), the RuvC domain, and the stop (STP) domain (Fig. 1C) [20]. Catalytic residues corresponding to the previously characterized RuvC catalytic triad (D394, E606, D695) were likewise conserved, underscoring functional conservation within this subfamily (Fig. 1D) [21]. These Cas12j loci were found primarily in large bacteriophages with genome sizes of approximately 100 kb, a feature consistent with earlier reports documenting Cas12j in expanded phage genomes [16]. The samples originated from ecologically diverse habitats, encompassing wetlands, lakes, sediments, coastal areas, and intertidal zones, suggesting that Cas12j may be more widespread in aquatic phage populations than previously observed.

### Design, construction, and *in vitro* characterization of Cas12j orthologues

To test the catalytic activities of the newly identified Cas12j orthologues *in vitro*, we designed, synthesized, and constructed the bacterial expression clones for each Cas12j protein (Supplementary files S1A; Supplementary Table S1). The purified Cas12j orthologues were assessed for nuclease activity using *in vitro* dsDNA cleavage assays. Cas12j-8 and LbCas12a were included as control proteins. For these assays, both circular and BglI-linearized dsDNA substrates (Supplementary files S1C) containing target sequences flanked by 5′-TTG-3′ protospacer adjacent motifs (PAMs) (Supplementary Fig. S1A) - an optimal variant of the consensus 5′-TTN-3′ PAM of Cas12j proteins were incubated with the assembled RNP complexes. Cleavage activity was evaluated by monitoring the DNA fragment generation after incubation of each Cas12j orthologues with BglI-linearized dsDNA substrates. The activity of different Cas12j orthologues on circular dsDNA substrates was assessed by incubating the Cas12j proteins with the template for 45 minutes followed by 15 minutes BsaIHFv2 restriction enzyme digestion to facilitate fragment release. All eight Cas12j proteins, along with the control proteins, exhibited specific target cleavage activity on linear and circular plasmids with varying cleavage efficiencies, confirming their catalytic potential *in vitro* (Supplementary Fig. S1B and C). Interestingly, all the Cas12j proteins showed greater cleavage efficiencies on circular targets. These findings underscore the potential of Cas12j variants as promising candidates for *in vivo* gene editing applications.

### Gene editing activity of Cas12j orthologues in mammalian cells

We next assessed the catalytic activities of the Cas12j orthologues *in cellula* in mammalian cells. Specifically, we evaluated the activities of eight Cas12j orthologues (Cas12j-11 to -18) alongside LbCas12a and Cas12j-8 as controls, in cultured HEK293T cells. The assays targeted the *CLTA* target-1 and *EMX* target-1 genomic regions, both of which contain a 5′-TTC-3′ PAM sequence. To facilitate expression in HEK293T cells, we human-codon optimized the eight Cas12j orthologues and constructed these clones in mammalian expression plasmids (Fig. 2A; Supplementary file S1B; Supplementary Table S1). Additionally, crRNAs specific to each Cas12j protein were designed to target the *CLTA* target-1 and *EMX* target-1 regions were independently expressed under the *U6* promoter (Supplementary file S1D; Supplementary Tables S4-S5).

**Figure 2.**
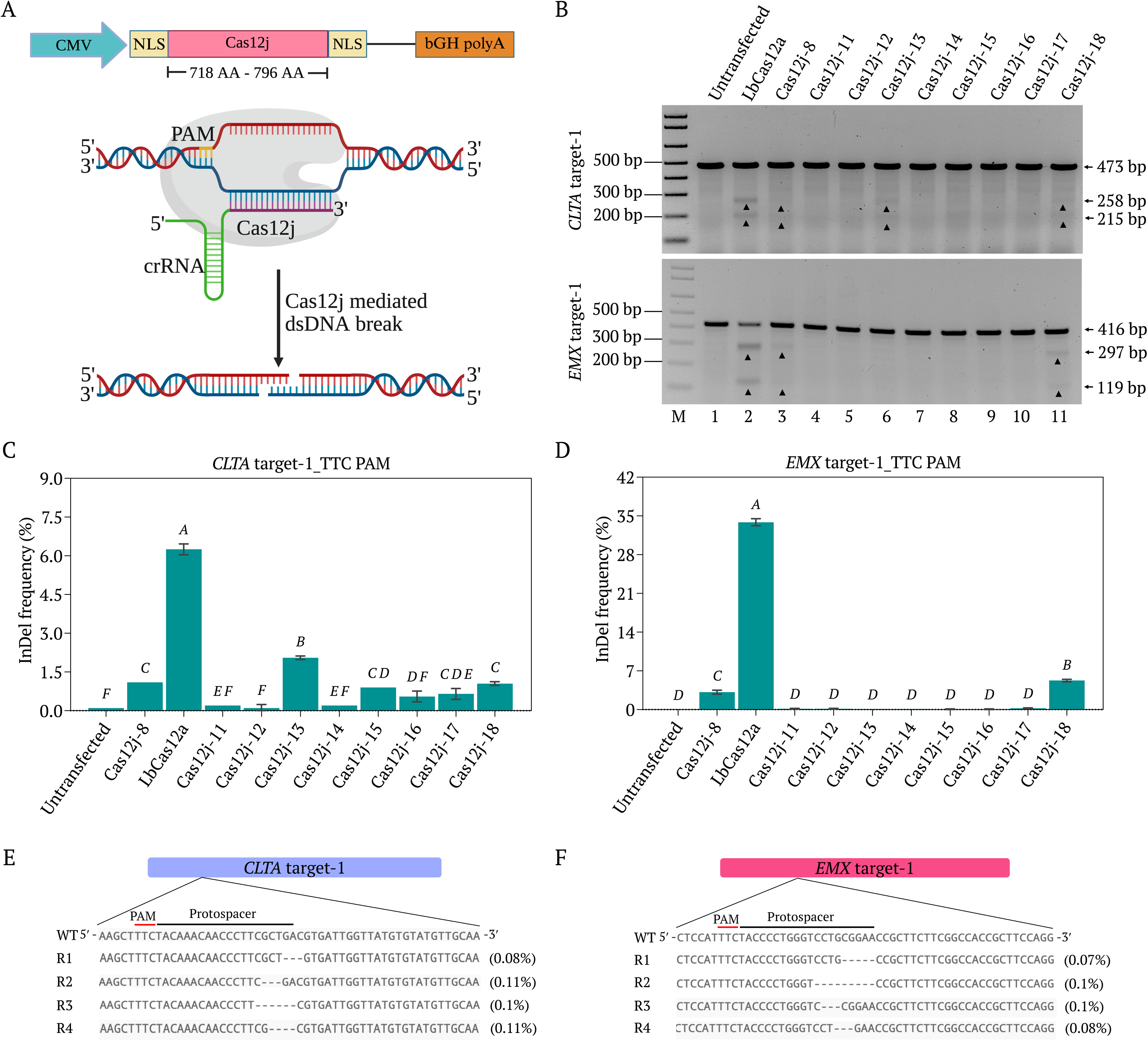
Gene editing activity of Cas12j proteins in mammalian cells. **(A)** Sketch showing the plasmid map for the mammalian cell expression of Cas12j proteins under CMV promoter and Cas12j variants mediated dsDNA break. **(B)** Gel images showing the T7EI assay for the evaluation of genome editing activity of LbCas12a (Lane 2), Cas12j-8 (Lane 3) and Cas12j-11 to -18 (Lanes 4-11) in HEK293T cells at *CLTA* target-1 and *EMX* target-1 regions, independently. Lane 1 is the T7EI of the DNA from untransfected cells used as negative control. Lane M is the 1kb plus marker. The arrow heads in both the gels indicate the T7EI enzyme cleavage products. **(C)** Deep amplicon sequencing data showing the variable indel efficiencies at *CLTA* target-1 using different Cas12j variants. Data represent mean with SD (n = 2 biologically independent replicates). Statistical analysis was performed using one-way ANOVA followed by Tukey’s multiple comparisons test. Bars labeled with different letters are significantly different (*P* < 0.05). Bars sharing the same letter are not significantly different (*P* > 0.05). **(D)** Deep amplicon sequencing data showing the variable indel efficiencies at *EMX* target-1 using different Cas12j variants. Data represent mean with SD (n = 2 biologically independent replicates). *P*-values are calculated as in (C). All statistical analysis was performed using GraphPad Prism 10 software. **(E)** Few of the Cas12j-13 produced deletions of *CLTA* target-1, PAM and target sequences indicated. **(F)** Some of the Cas12j-18 generated deletions of *EMX* target-1, PAM and target sequences are indicated. R1-R4 represents the different predominant deep amplicon sequencing reads. Editing outcomes are mentioned in the closed brackets next to the each read.

Three days post-transfection, T7EI assays were performed to assess the genome editing activity. Among the tested Cas12j orthologues, Cas12j-13 and -18 demonstrated detectable editing activity on *CLTA* target-1. Whereas, *EMX* target-1 was edited only by Cas12j-18. The performance of Cas12j-13 and -18 was comparable to that of the previously characterized Cas12j-8 control (Fig. 2B). To gain deeper insights, we performed deep amplicon sequencing, confirming that Cas12j-13 and Cas12j-18 were the most active orthologues. Specifically, Cas12j-13 generated 2.0–2.1% indels at *CLTA* target-1, while Cas12j-18 generated 1.0–1.1% indels at *CLTA* target-1 and 5.1–5.4% indels at *EMX* target-1 (Fig. 2C and D). Minimal editing activity was also detected for Cas12j-15, Cas12j-16, and Cas12j-17 at *CLTA* target-1 (Fig. 2C). Further analysis of sequencing reads of Cas12j-13 and -18 on *CLTA* target-1 and on *EMX* target-1, respectively, confirmed that the predominant editing outcomes were small deletions (Fig. 2E and F).

Overall, deep amplicon sequencing validated that Cas12j-13 and Cas12j-18 possess measurable genome-editing activity in mammalian cells, making them promising candidates for further optimization and engineering. However, their current editing efficiencies remain suboptimal compared to LbCas12a, highlighting the need for further enhancements to maximize their potential for broader gene-editing applications.

### Development of T5Exo-Cas12j editors for efficient gene editing

Given the modest editing efficiencies observed by the Cas12j orthologues, we explored whether fusing them with catalytic nuclease domains could enhance their performance. Previous studies have shown that degradation of DNA ends at double-strand breaks (DSBs) by exonucleases can promote larger deletions via the non-homologous end joining (NHEJ) or microhomology mediated end-joining (MMEJ) repair pathways. Among these, T5 exonuclease (T5Exo), a 33 kDa enzyme that digests single-stranded and double-stranded DNA in the 5′ to 3′ direction, has been reported to improve the editing efficiency of Cas9, Cas12a, and IscB nucleases [22–25]. To harness this potential, we fused T5Exo to the N termini of each Cas12j orthologues and constructed the mammalian expression plasmids (Fig. 3A; Supplementary Table S2; Supplementary file S1B). The resulting T5Exo-Cas12j constructs had a size range of 1025–1103 AA, smaller than common Cas enzymes like Cas9 or Cas12a.

**Figure 3.**
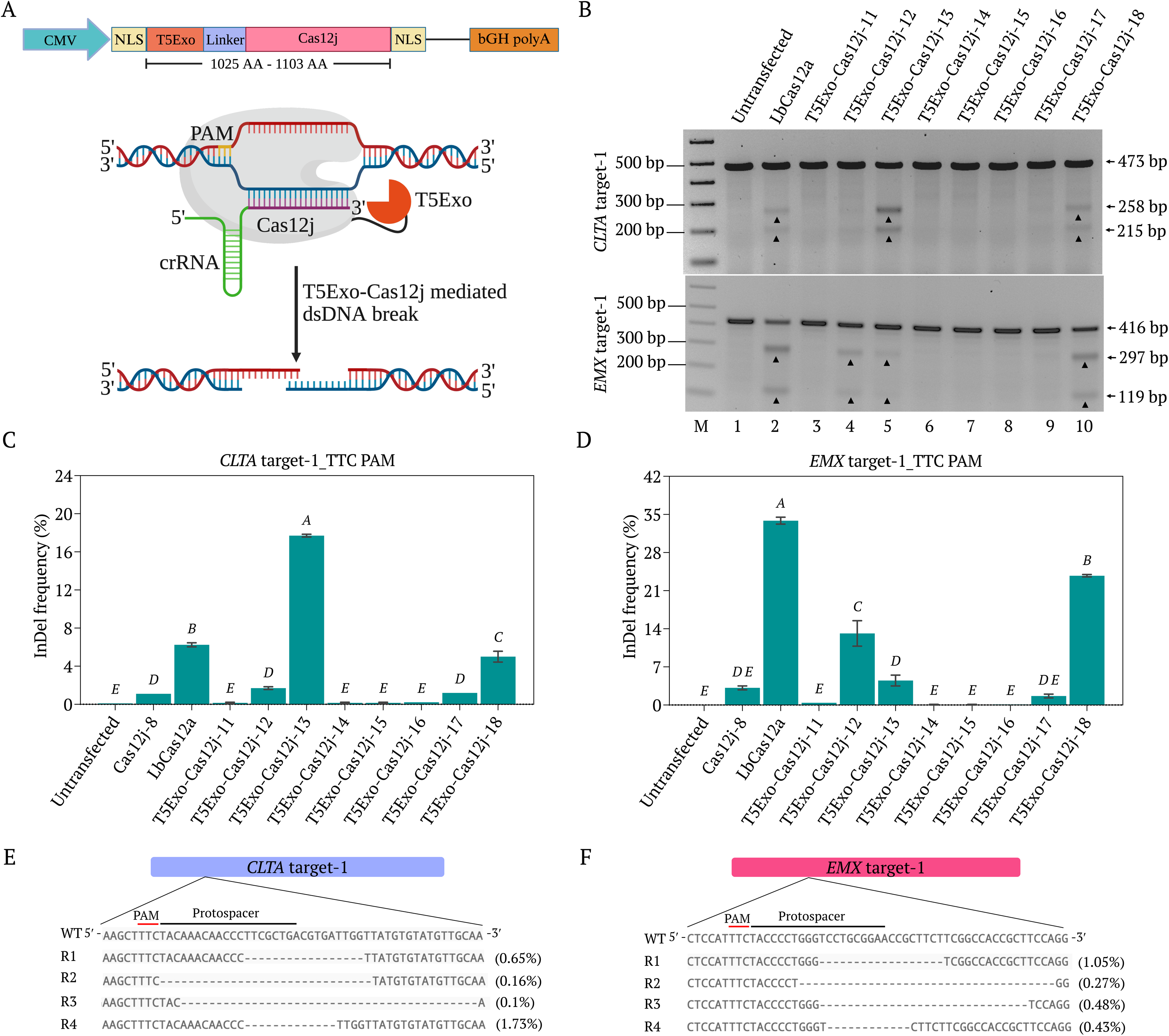
Gene editing activity of T5Exo-Cas12j fusion proteins in mammalian cells. **(A)** Sketch showing the plasmid map expressing the T5Exo-Cas12j proteins under CMV promoter and T5Exo-Cas12j proteins mediated dsDNA break. **(B)** Gel images showing the T7EI assay results of *CLTA* target-1 and *EMX* target-1 gene editing by LbCas12a (Lane 2) and T5Exo-Cas12j-11 to -18 (Lanes 3-10) in HEK293T cells. Lane 1 is the T7EI of the DNA from untransfected cells used as negative control. Lane M is the 1kb plus marker. The arrow heads in both the gels indicate the T7EI enzyme cleavage products. **(C)** Deep amplicon sequencing data showing the variable indel efficiencies at *CLTA* target-1 using different T5Exo-Cas12j variants. Data represent mean with SD (n = 2 biologically independent replicates). Statistical analysis was performed using one-way ANOVA followed by Tukey’s multiple comparisons test. Bars labeled with different letters are significantly different (*P* < 0.05). Bars sharing the same letter are not significantly different (*P* > 0.05). **(D)** Deep amplicon sequencing data showing the variable indel efficiencies at *EMX* target-1 using different T5Exo-Cas12j variants Data represent mean with SD (n = 2 biologically independent replicates). *P*-values are calculated as in (C). All statistical analysis was performed using GraphPad Prism 10 software. **(E)** Few of the T5Exo-Cas12j-13 produced deletions of *CLTA* target-1, PAM and target sequences indicated. **(F)** Some of the T5Exo-Cas12j-18 generated deletions of *EMX* target-1, PAM and target sequences are indicated. R1-R4 represents the different predominant deep amplicon sequencing reads. Editing outcomes are mentioned in the closed brackets next to the each read.

We evaluated the genome-editing performance of all the T5Exo-Cas12j proteins in HEK293T cells targeting *CLTA* target-1 and *EMX* target-1 (5′-TTC-3′ PAM) by T7EI assay. Our experiments revealed that T5Exo-Cas12j-12, -13, and -18 demonstrated better editing efficiencies (Fig. 3B). Additionally, the deep amplicon sequencing results revealed that, T5Exo-Cas12j-13 exhibited 17.6-17.8% indels on *CLTA* target-1 and 3.8-5.2% indels on *EMX* target-1, while T5Exo-Cas12j-18 exhibited 4.6-5.4% indels on *CLTA* target-1 and 23.6-23.9% indels on *EMX* target-1 (Fig. 3C and D). Among all T5Exo fusions, T5Exo-Cas12j-13 achieved significantly higher editing efficiency compared to LbCas12a and Cas12j-8 on *CLTA* target-1. Whereas, T5Exo-Cas12j-12 and -18 proteins showed higher editing efficiency compared to Cas12j-8 on *EMX* target-1 (Fig. 3C and D). The sequencing reads of T5Exo-Cas12j-13 and -18 on *CLTA* target-1 and on *EMX* target-1 respectively, showed that the reads are majorly the longer deletions compared to wild-type Cas12j proteins (Fig. 3E and F). These results demonstrate the importance of the T5Exo fusion for the elevated indel efficiencies.

Next, we compared the editing efficiencies of the T5Exo-Cas12j and Cas12j editors, guided by the mature crRNA (26-nt) and the pre-processed crRNA (36-nt) on *CLTA* target-1 (Supplementary Table S4). All four T5Exo-Cas12j editors (-12, -13, -17, -18) and Cas12j editors (-12, -13, -17, -18) showed lower editing efficiencies when guided by the mature crRNA (Supplementary Fig. S2A-D).

### Cas12j and T5Exo-Cas12j editors: activity on additional genomic targets containing different PAM sequences

To evaluate the editing efficiencies of T5Exo-Cas12j editors (-12, -13, -17, and -18) on alternative genomic targets, we designed and expressed crRNAs targeting *CLTA* target-2, *CLTA* target-3, and *EMX* target-2, which contain TTG, TTA, and TTT PAM sequences, respectively (Supplementary Tables S4-S5). The crRNAs were expressed under *U6* promoters, and genome-editing activity was assessed via T7EI assays. The results revealed no detectable editing activity by any T5Exo-Cas12j editors at these targets (Supplementary Fig. S3A-D).

To further investigate PAM preference, we tested additional genomic targets containing TTC, TTG, TTA, and TTT PAM sequences using all Cas12j and T5Exo-Cas12j editors (Supplementary Fig. S4A). Deep amplicon sequencing confirmed minimal editing activity (<5% indels) only for T5Exo-Cas12j-12 at *HBB* target-3 (TTG PAM) and *HBB* target-4 (TTT PAM), as well as T5Exo-Cas12j-13 at *HBB* target-4 (TTT PAM). All other targets tested with Cas12j and T5Exo-Cas12j editors failed to generate significant indels (Supplementary Fig. S4B).

To better understand this sequence dependency, we performed *in silico* analysis of all the target sequences tested *in cellula*. Interestingly, the targets that exhibited editing activity (*CLTA* target-1 and *EMX* target-1) contained a 5′-TAC trinucleotide within the target sequence, whereas all other tested target sequences lacked 5′-TAC. Based on this observation, we hypothesize that Cas12j-mediated editing is highly dependent on the presence of a 5′-TAC sequence in the target sequence, with other trinucleotide sequences tested in this study displaying little or no detectable editing activity (Supplementary Fig. S5).

### Requirement of 5**′**-TAC trinucleotides in the target sequence for robust Cas12j editing activity

While all Cas12j proteins exhibited robust catalytic activity *in vitro*, their *in cellula* editing efficiency remained relatively modest. To enhance their activity, we engineered chimeric fusions of Cas12j with the T5Exo enzyme, leading to improved genome-editing performance. To systematically evaluate the editing efficiency of these engineered editors, we tested multiple genomic targets and compared their activity to LbCas12a. Strikingly, we observed significantly enhanced editing efficiency when the DNA target sequence started with a 5′-TAC trinucleotide after the PAM sequence (Fig. 4A). To determine whether this trinucleotide sequence is essential for robust Cas12j-mediated editing, we targeted multiple regions within the *CLTA* gene (Fig. 4B; Supplementary Table S5). Notably, target sequences containing 5′-TAC in after TTN PAM sequences significantly improved the editing efficiency of Cas12j editors, reaching levels comparable to LbCas12a.

**Figure 4.**
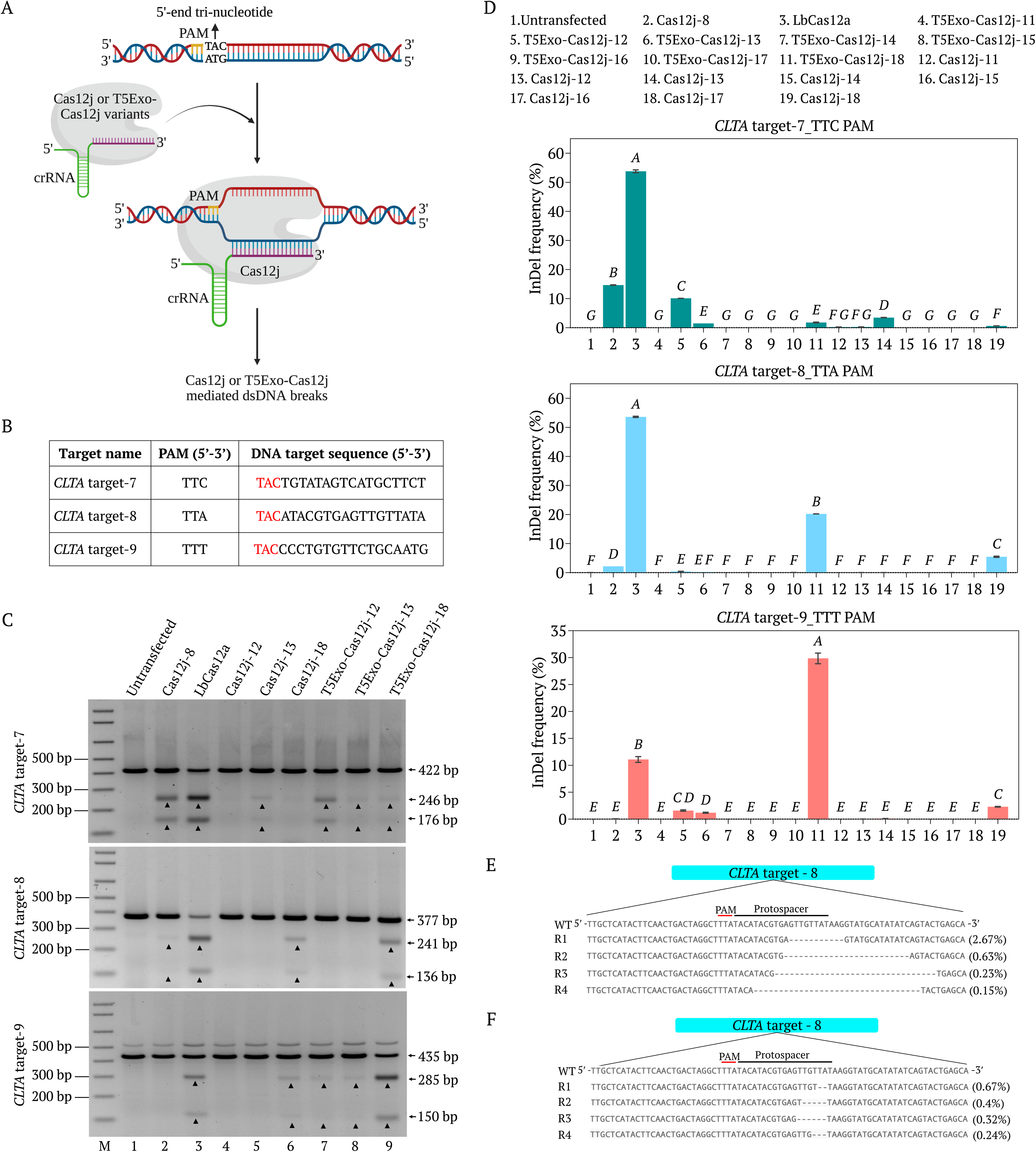
Gene editing activity of Cas12j and T5Exo-Cas12j fusion proteins on different targets in mammalian cells using crRNAs with 5′-end TAC tri-nucleotide sequence. **(A)** Sketch showing the Cas12j and T5Exo-Cas12j variants mediated editing of targets containing 5′-TAC tri-nucleotide sequence. **(B)** Table showing the different DNA target sequences and corresponding PAM sequences. **(C)** Gels showing the T7EI assay results of the genome editing activity using Cas12j-8 (Lane 2), LbCas12a (Lane 3), Cas12j-12, -13, -18 (Lanes 4-6) and T5Exo-Cas12j-12, -13, -18 (Lanes 7-9) in HEK293T cells at *CLTA* target-7, -8, -9 regions, independently. Lane 1 is the T7EI for the DNA from untransfected cells used as negative control. Lane M is the 1kb plus marker. The arrow heads indicate the T7EI enzyme cleavage products. **(D)** Deep amplicon sequencing data showing the variable indel efficiencies at *CLTA* target-7, -8, -9 regions using different T5Exo-Cas12j and Cas12j variants. Data represent mean with SD (n = 2 biologically independent replicates). Statistical analysis was performed using one-way ANOVA followed by Tukey’s multiple comparisons test. Bars labeled with different letters are significantly different (*P* < 0.05). Bars sharing the same letter are not significantly different (*P* > 0.05). All statistical analysis was performed using GraphPad Prism 10 software. **(E)** Some of the T5Exo-Cas12j-18 produced deletions of *CLTA* target-8, PAM and target sequences are indicated. **(F)** Few of the Cas12j-18 generated deletions of *CLTA* target-8, PAM and target sequences indicated. R1-R4 represents the different predominant deep amplicon sequencing reads. Editing outcomes are mentioned in the closed brackets next to the each read.

T7EI and deep amplicon sequencing confirmed that the presence of 5′-TAC at the beginning of the target sequence consistently correlated with higher editing efficiencies (Fig. 4C and D). For example, targeting *CLTA* target-7 containing TAC at the 5′ end and a TTC PAM enabled T5Exo-Cas12j-12 to achieve >10% editing efficiency. Moreover, targeting *CLTA* target-8 containing a TAC sequence after TTA PAM resulted in significant editing by Cas12j-18 and T5Exo-Cas12j-18 when compared to Cas12j-8. Notably, T5Exo-Cas12j-18 outperformed LbCas12a when targeting *CLTA* target-9, demonstrating superior editing efficiency (Fig. 4C and D). In contrast, none of the wild-type Cas12j editors or T5Exo-Cas12j-11, -14, -15, -16, or -17 displayed detectable genome editing at *CLTA* target-7, -8, and -9 sites (Supplementary Fig. S6; Fig. 4D). Further sequencing analysis of T5Exo-Cas12j-18 and Cas12j-18 editing at *CLTA* target-8 revealed distinct mutation patterns, with T5Exo-Cas12j-18 generating wider deletions, while Cas12j-18 produced shorter deletions (Fig. 4E and F).

To investigate the significance of TAC as part of the PAM sequence, we designed crRNAs targeting *CLTA*-target-1a, *CLTA*-target-1b, *CLTA*-target-9a, and *CLTA*-target-9b, all of which contained a TTCTAC PAM sequence (Supplementary Tables S4-S5). First, we evaluated the genome-editing activity of Cas12j-11 to -18 and their T5Exo-Cas12j fusion variants on *CLTA*-target-1a and *CLTA*-target-9a, using 20-nt crRNAs. T7EI assays showed no detectable target editing (Supplementary Fig. S7A). Next, to determine whether truncating the crRNA sequence would affect editing efficiency, we tested Cas12j-13, Cas12j-17, and Cas12j-18, along with their T5Exo-Cas12j variants, on *CLTA*-target-1b and *CLTA*-target-9b, using 17-nt crRNAs. However, T7EI assays again showed no detectable editing activity (Supplementary Fig. S7B). These findings suggest that TTNTAC as PAM does not have any impact on Cas12j-mediated editing, indicating that 5′-TAC as target sequence functions optimally rather than as an extension of the PAM.

### Editing disease relevant genes using Cas12j editors

Next, we attempted to substantiate the requirement of TAC trinucleotide to edit disease relevant genes and assess the usefulness of Cas12j editors compared with LbCas12a. Therefore, we designed two targets for *Caspase*-3 and two for *AIFM* (Apoptosis-Inducing Factor Mitochondrial 1) genes starting with 5′-TAC trinucleotide sequence with TTN PAM sequences (Fig. 5A; Supplementary Table S5). We transfected Cas12j-8 and LbCas12a control effectors to assess the catalytic efficiencies of the T5Exo-Cas12j editors. We transfected these editors with the crRNA encoding DNA into HEK293T cells and assessed the on-target catalytic activities using T7EI assay and deep amplicon sequencing studies. Intriguingly, our data showed that for *AIFM* target-1, T5Exo-Cas12j-12, -13, and -18 exhibited robust editing activities. Moreover, for the *AIFM* target-2, Cas12j-8, T5Exo-Cas12j-11, -12, -13, -17, and -18 exhibited robust editing activities (Fig. 5B and C). Of note, T5Exo-Cas12j-12 and -18 exhibited matched or exceeding catalytic activities compared with LbCas12a. Moreover, the wild-type Cas12j variants such as Cas12j-12, and -18 are also showing elevated editing efficiencies on *AIFM* target-2 (Fig. 5C; Supplementary Fig. S8). For the *Caspase-3* target-1, T5Exo-Cas12j-12, and -18 exhibited robust catalytic activities. Interestingly, T5Exo-Cas12j-12 and -18 exceeded the LbCas12a editing efficiency significantly, indicating the usefulness of these editors in the translational applications (Fig. 5B and C). Moreover, for the *Caspase-3* target-2, T5Exo-Cas12j-12 also outperformed the LbCas12a and the Cas12j-8, confirming the robust activities of these editors (Fig. 5B and C). Whereas the wild-type Cas12j variants did not show any editing of *Caspase-3* target-1 and *Caspase-3* target-2 (Fig. 5C; Supplementary Fig. S8). Intriguingly, T5Exo-Cas12j-12 and -18 exhibited catalytic efficiency of >20%, comparable or exceeding those of LbCas12a across these disease relevant targets pointing to its potential utility in precision editing of these targets (Fig. 5C). These data collectively demonstrate that T5Exo-Cas12j editors provide an efficient and compact genome-editing tool applicable for disease relevant genes, supporting its potential therapeutic relevance.

**Figure 5.**
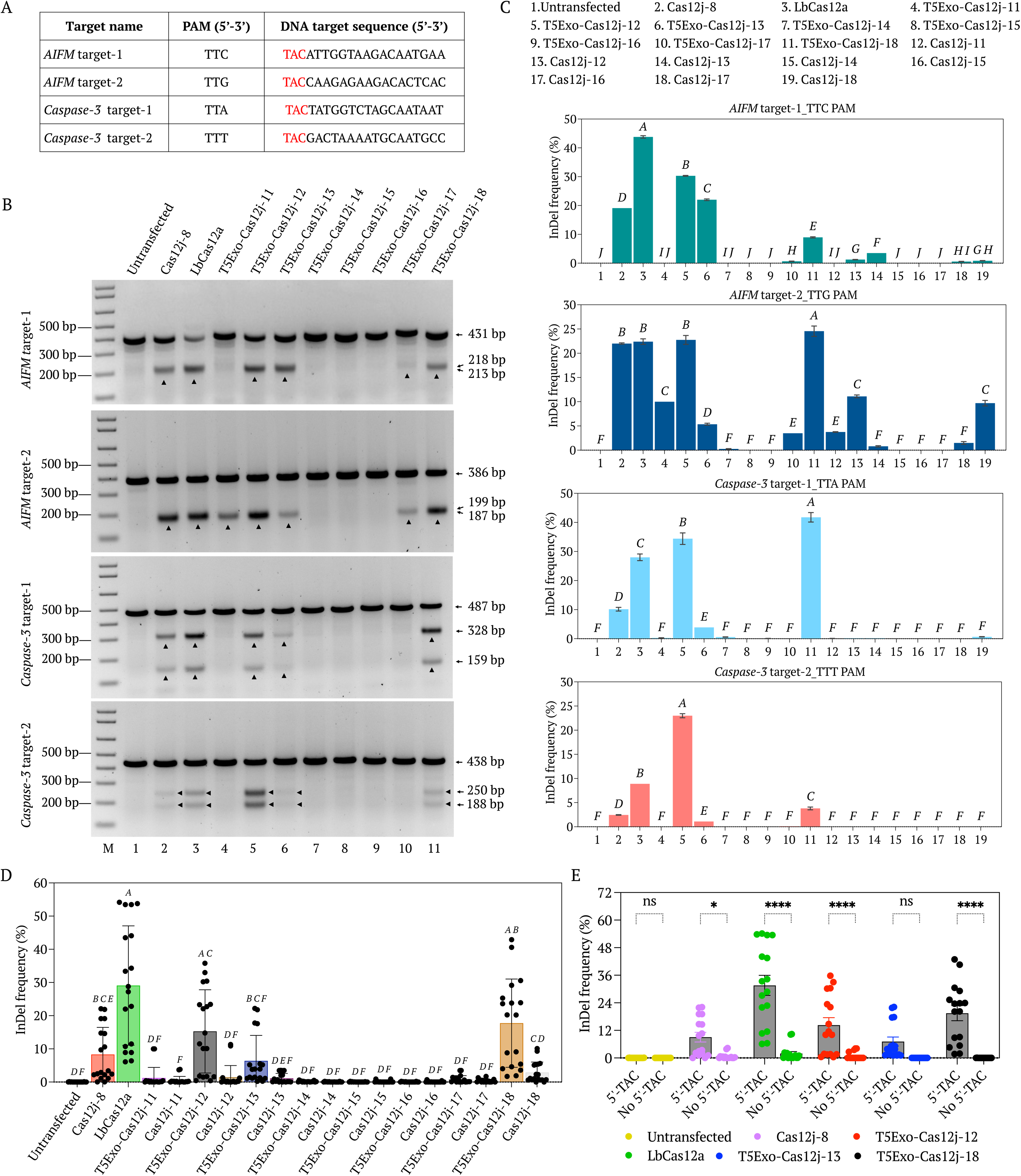
Gene editing activity of Cas12j and T5Exo-Cas12j fusion proteins on disease related genes in mammalian cells using crRNAs with 5′-end TAC tri-nucleotide sequence. **(A)** Table showing the different DNA target sequences and respective PAM sequences. **(B)** Gels showing the T7EI assay results of the genome editing activity using Cas12j-8 (Lane 2), LbCas12a (Lane 3), and T5Exo-Cas12j-11 to -18 (Lanes 4-11) in HEK293T cells at *AIFM* target-1, -2, and *Caspase-3* target-1, -2 regions, independently. Lane 1 is the T7EI for the untransfected cells used as negative control. Lane M is the 1kb plus marker. The arrow heads indicate the T7EI enzyme cleavage products. **(C)** Deep amplicon sequencing data showing the variable indel efficiencies at *AIFM* target-1, -2, and *Caspase-3* target-1, -2 regions using different T5Exo-Cas12j and Cas12j variants. Data represent mean with SD (n = 2 biologically independent replicates). Statistical analysis was performed using one-way ANOVA followed by Tukey’s multiple comparisons test. Bars labeled with different letters are significantly different (*P* < 0.05). Bars sharing the same letter are not significantly different (*P* > 0.05). **(D)** Comparison of different T5Exo-Cas12j editors over their wild-type counter parts along with Cas12j-8 and LbCas12a proteins targeting multiple genes (*CLTA* target-1, -7, -8, -9, *EMX* target-1, *AIFM* target-1, -2, *Caspase-3* target-1 and -2) containing 5′-TAC at DNA target sites. Data represent mean with SD (n = 18; whole independent data points). *P*-values are calculated as mentioned in (C). **(E)** Cleavage of multiple targets containing 5′-TAC (*CLTA* target-1, -7, -8, -9, *EMX* target-1, *AIFM* target-1, -2, and *Caspase-3* target-1) and lacking 5′-TAC (*CLTA* target-4, -5, -6, -12, *HBB* target-1, -2, -3, and -4) next to the PAM in the target DNA sequence using selected T5Exo-Cas12j editors along with Cas12j-8 and LbCas12a proteins. Data represent mean with SD (n = 16; whole independent data points). *P*-values are calculated using a two-way ANOVA, Bonferroni test (*P*-values indicated are: ns is not significant (0.1234), * (0.0332), ** (0.0021), *** (0.0002), **** (<0.0001)). All statistical analysis was performed using GraphPad Prism 10 software.

To evaluate the overall editing efficiencies of various Cas12j editors and LbCas12a across different targets, we performed statistical comparisons. T5Exo-Cas12j-12 and T5Exo-Cas12j-18 demonstrated significantly higher editing efficiencies across multiple DNA targets containing the 5′-TAC motif adjacent to the PAM, compared to their respective wild-type counterparts (Fig. 5D). Furthermore, both T5Exo-Cas12j-12 and -18 outperformed Cas12j-8 in editing efficiency (Fig. 5D).

Analysis of the catalytic activities of T5Exo-Cas12j editors (T5Exo-Cas12j-12, -13, and -18), along with Cas12j-8 and LbCas12a, across multiple genomic targets, with or without the 5′-TAC trinucleotides immediately downstream of the PAM revealed notable trends. T5Exo-Cas12j-12 and -18 exhibited robust editing efficiencies at targets harboring the 5′-TAC motif. In contrast, targets lacking the 5′-TAC sequence showed minimal or no detectable editing activity relative to TAC-containing sites (Fig. 5E). Interestingly, Cas12j-8 and LbCas12a also displayed enhanced editing efficiencies when targeting regions that included the 5′-TAC sequence. Collectively, these results highlight the 5′-TAC trinucleotide sequence as a critical determinant of Cas12j-mediated editing efficiency. This discovery provides a valuable design parameter for optimizing Cas12j-based genome editing systems, enhancing their utility in translational research and therapeutic applications.

Although previous studies have explored Cas12j’s single-locus genome editing in mammalian and microbial systems and showed modest efficiencies. In this study, we attempted to assess the utility of our T5Exo-Cas12j-18 editor for multiplex editing which is of particular importance in simultaneous engineering of multiple genomic loci. We transfected T5Exo-Cas12j-18 with multiple crRNAs targeting *CLTA* target-9, *EMX* target-1, *AIFM* target-2, and *Caspase-3* target-1 in HEK293T cells. Next, we conducted T7EI assays, and our data showed that the T5Exo-Cas12j-18 editor exhibited detectable catalytic activities in these four targets (Supplementary Fig. S9A and B). These results demonstrate that T5Exo-Cas12j enables multiplex genome editing at disease-relevant loci and highlights its potential as a powerful genome editing tool for potential therapeutic applications.

### Characterizing the nature of edits using deep amplicon sequencing

To investigate the nature, position, and efficiency of edits introduced by T5Exo-Cas12j effectors, we analyzed deep amplicon sequencing data from *CLTA* target-8, *CLTA* target-9, *AIFM* target-1, and *AIFM* target-2 using the CRISPResso2 pipeline. Multiple Cas12j effectors, including T5Exo-Cas12j-13, T5Exo-Cas12j-18, Cas12j-13, Cas12j-18, LbCas12a, and Cas12j-8, were tested to assess their respective editing patterns, particularly the nature of insertions and deletions (indels) introduced at the target sites.

Quantification of editing efficiencies revealed that T5Exo-Cas12j-13 and -18 exhibited comparable or superior performance compared to Cas12j-8, while LbCas12a displayed a distinct editing profile, favoring extensive deletions (Supplementary Fig. S6A). Although LbCas12a consistently exhibited high editing efficiency, T5Exo-Cas12j effectors demonstrated strong activity and, in certain cases, outperformed LbCas12a. Notably, T5Exo-Cas12j-18 surpassed LbCas12a when targeting *AIFM* target-2 and *CLTA* target-9, emphasizing its robust catalytic efficiency (Supplementary Fig. S6A). These findings confirm the effectiveness of T5Exo-Cas12j effectors and their potential for precision genome editing.

Analysis of amplicon sequencing data revealed that the editing window was largely restricted to the near PAM region, supporting the precision of these editors (Supplementary Fig. S6B). Further comparative analysis demonstrated that small deletions of up to 25 bp were the predominant mutation type, with all editors favoring deletions over insertions. Interestingly, T5Exo-Cas12j effectors exhibited a higher frequency of deletions compared to other editors, yet these deletions remained confined within a 0–25 bp range (Supplementary Fig. S6C). These insights provide a deeper understanding of Cas12j-based genome editing precision and reinforce its potential for customized genetic modifications in therapeutic applications.

### Assessment of off-target effects highlights the specificity of T5Exo-Cas12j editors

We employed the Cas-OFFinder online tool [26] to predict potential off-target sites for the most efficiently edited endogenous targets by T5Exo-Cas12j/crRNA complexes in HEK293T cells. Notably, all highly active target sites in HEK293T cells lacked predicted off-targets with a single nucleotide mismatch. Previous studies have demonstrated that even off-targets with two or three nucleotides mismatches can be susceptible to cleavage by LbCas12a and Cas9 effectors [27].

Therefore, we extended our search to include off-target sites with two and three nucleotide mismatches, identifying several PAM-proximal and PAM-distal potential off-targets for *CLTA* target-7, -8, *AIFM* target-1, and *Caspase-3* target-1 genomic regions (Fig. 6; Supplementary Table S7). Genomic DNA from HEK293T cells edited at these on-target sites by T5Exo-Cas12j-12, -13, and -18 was independently PCR-amplified using off-target specific primers and followed by deep amplicon sequencing. The sequencing analysis revealed no off-target indels at any of the PAM-proximal or PAM-distal loci examined (Fig. 6). These results demonstrated the high specificity of T5Exo-Cas12j editors and supporting their potential application in precise genome editing tools for therapeutic use.

**Figure 6.**
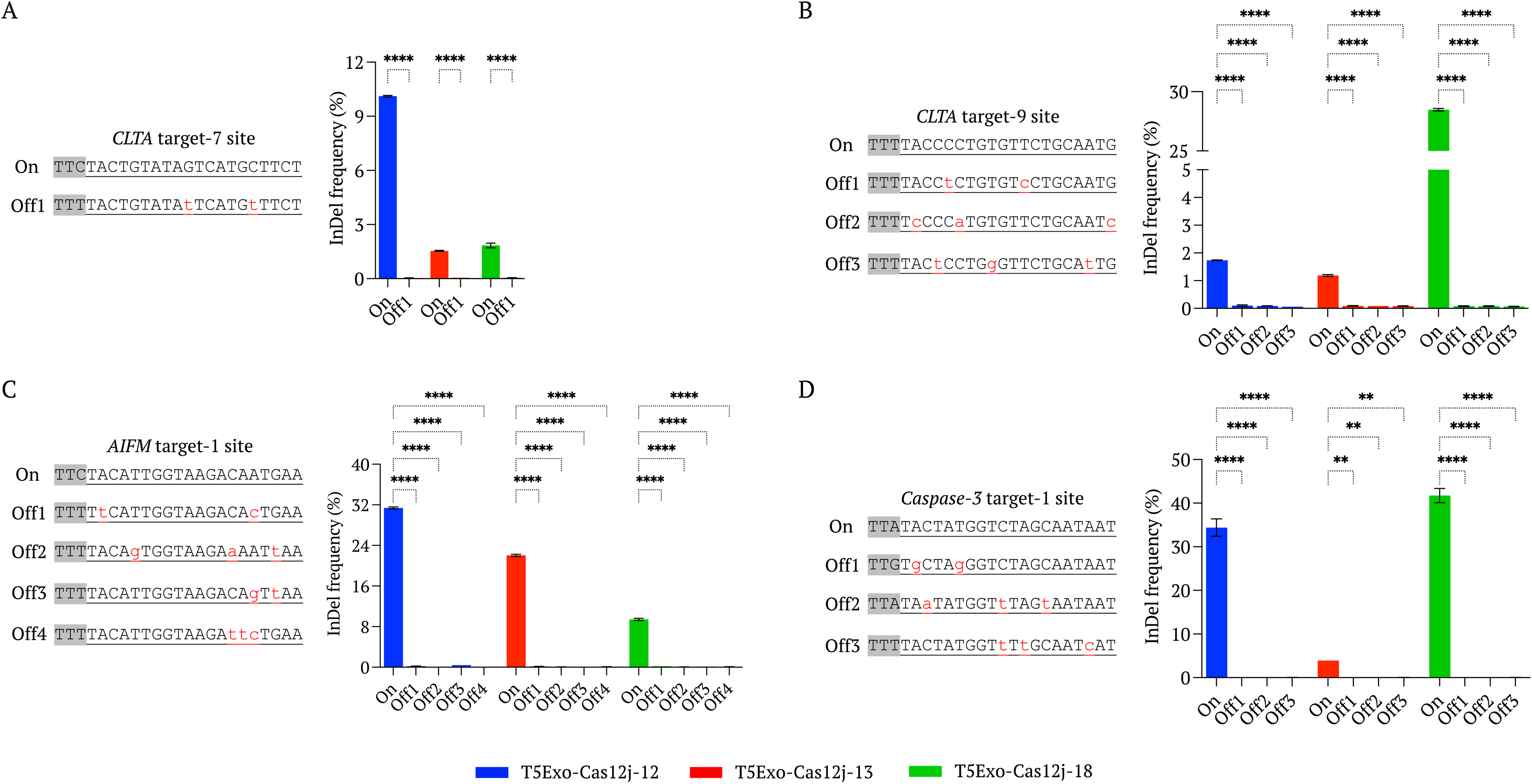
Off-targeting activity of T5Exo-Cas12j fusion proteins in HEK293T cells. **(A-D)** DNA sequences showing are different On-target and their potential off-target sequences for *CLTA* target-7, -8, *AIFM* target-1, and *Caspase-3* target-1genes. PAM sequences are highlighted in grey and target DNA sequences are underlined. Plots showing are the deep amplicon sequencing data of T5Exo-Cas12j editors on *CLTA* target-7, -8, *AIFM* target-1, and *Caspase-3* target-1 on-targets and respective off-targets. Data represent mean with SD (n = 2 biologically independent replicates). *P*-values are calculated using a one-way ANOVA followed by Tukey’s test (*P*-values indicated are: ns is not significant (0.1234), * (0.0332), ** (0.0021), *** (0.0002), **** (<0.0001)). Statistical analysis was performed using GraphPad Prism 10 software.

### T5Exo-Cas12j mediated gene editing in human K562 cell line

To broaden the applicability of T5Exo-Cas12j editors, we evaluated their editing activity in the leukemia-derived lymphoblast cell line K-562, serving as a model for therapeutic applications. To investigate the requirement for a 5′-TAC trinucleotide immediately downstream of the PAM, we targeted genomic regions either containing the 5′-TAC motif (*AIFM* target-1, *CLTA* target-8, and -9) or lacking it (*HBB* target-2, *CLTA* target-5, and -12). Each gene target was independently edited using T5Exo-Cas12j-12, -13, and -18 variants, alongside Cas12j-8 and LbCas12a as controls (Supplementary Fig. S11A).

T7EI assays and deep amplicon sequencing consistently revealed higher indel frequencies at targets containing the 5′-TAC motif (Supplementary Fig. S11B and C). For example, T5Exo-Cas12j-12 achieved over 7.5% editing efficiency at the *AIFM* target-1 site, while T5Exo-Cas12j-18 demonstrated more than 6% editing efficiency at the *CLTA* target-8 site. Notably, T5Exo-Cas12j-18 outperformed LbCas12a at the *CLTA* target-9 site, achieving editing efficiency exceeding 9% (Supplementary Fig. S11C). On their respective targets, T5Exo-Cas12j variants exhibited significantly higher editing efficiencies compared to Cas12j-8 (Supplementary Fig. S11C). By contrast, none of the T5Exo-Cas12j variants generated detectable edits at targets lacking the 5′-TAC motif (*HBB* target-2, *CLTA* target-5, and -12) (Supplementary Fig. S11C). These findings further underscore the strict dependence of T5Exo-Cas12j activity on the presence of a 5′-TAC sequence.

There was no significant difference in T5Exo-Cas12j-mediated editing efficiencies between K-562 and HEK293T cells (Supplementary Fig. S11D). The relative editing activity profiles in K-562 cells closely mirrored those observed in HEK293T cells. Collectively, these results indicate that T5Exo-Cas12j editors are effective across different cell types and hold strong potential for therapeutic genome editing applications.

### Development of Be-(d)Cas12j editor for efficient genome editing

Base editors have been developed by fusing Cas9 nickase (nCas9) or dead Cas9 (dCas9) with a deaminase, enabling targeted single base-pair substitutions without introducing DSBs. However, Cas9-derived base editors are relatively large, complicating *in vivo* delivery, particularly with size-constrained vectors such as AAVs [28,29]. Given the compact size of Cas12j proteins, which makes them well-suited for viral vector-based delivery, we sought to develop a Cas12j-derived base editor by fusing catalytically inactive Cas12j ((d)Cas12j) with the TadA8e deaminase [30] at its N-terminus (Supplementary Table S3). This fusion, termed Be-(d)Cas12j, was designed to mediate adenine-to-guanine (A-to-G) substitutions (Fig. 7A).

**Figure 7.**
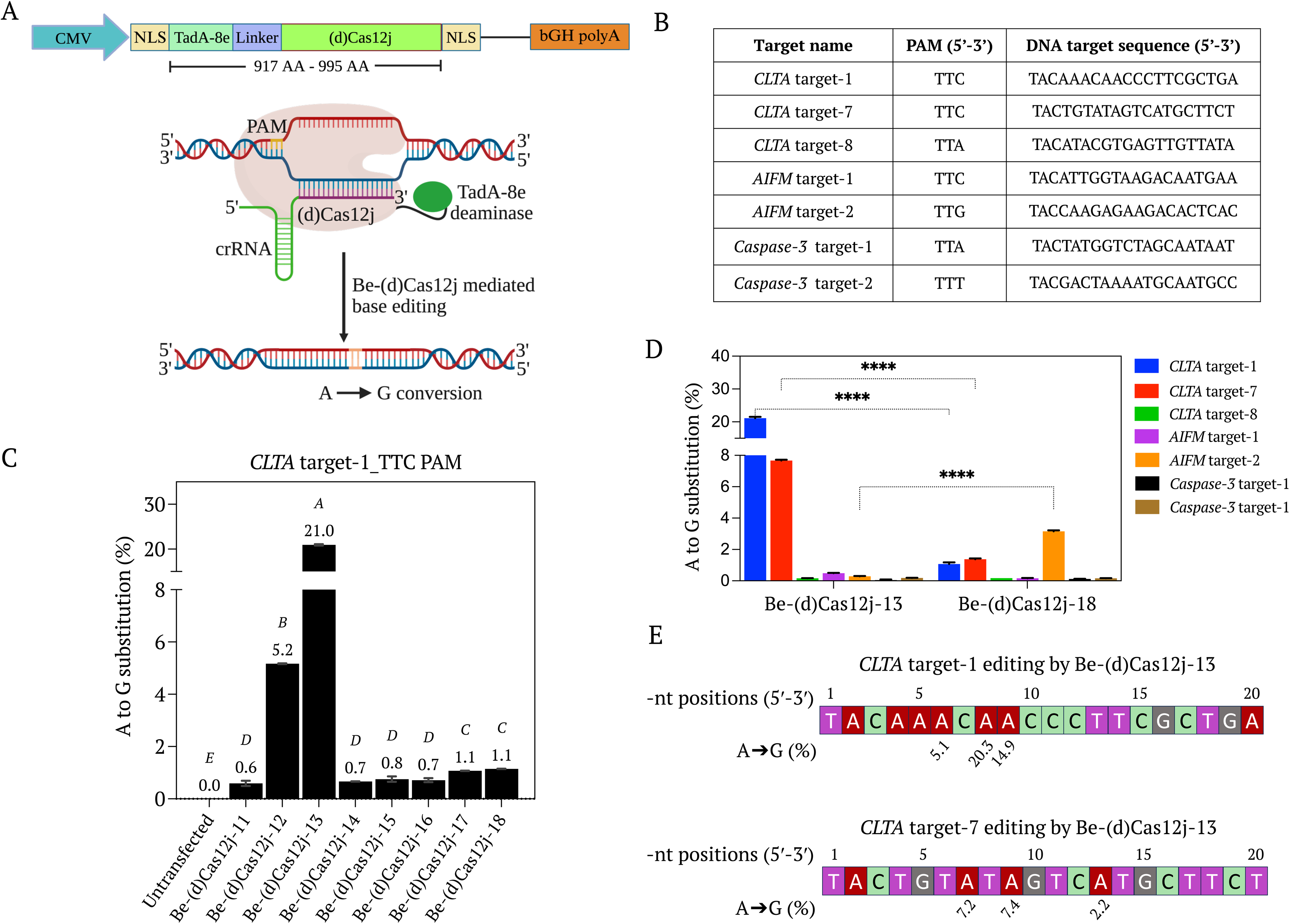
Base editing via newly discovered (d)Cas12j variants fused with TadA8e. **(A)** Sketch showing the plasmid map for the expression of TadA8e-(d)Cas12j (Be-(d)Cas12j) variants under the CMV promoters in mammalian cells. Illustration showing the targeted base substitution (A-to-G) using Be-(d)Cas12j base editor. **(B)** List of target sequences and corresponding PAM sequences employed for base editing in this study. **(C)** Deep amplicon sequencing data showing the A-to-G conversion percentage of different Be-(d)Cas12j-11 to -18 editors at *CLTA* target-1 region in HEK293T cells. Data represent mean with SD (n = 2 biologically independent replicates). Statistical analysis was performed using one-way ANOVA followed by Tukey’s multiple comparisons test. Bars labeled with different letters are significantly different (*P* < 0.05). Bars sharing the same letter are not significantly different (*P* > 0.05). **(D)** Deep amplicon sequencing data showing the A-to-G conversion percentage at different targets *CLTA* target-1, -7, -8, *AIFM* target-1, -2, and *Caspase-3* target-1, -2 regions in HEK293T cells using Be-(d)Cas12j-13 and -18 base editors. Data represent mean with SD (n = 2 biologically independent replicates). Statistical analysis was performed using two-way ANOVA followed by Tukey’s test (*P*-values indicated are: ns is not significant (0.1234), * (0.0332), ** (0.0021), *** (0.0002), **** (<0.0001)). Statistical analysis was performed using GraphPad Prism 10 software. **(E)** A-to-G conversion efficiencies at different nucleotide positions on *CLTA* target-1 and -7 by Be-(d)Cas12j-13.

Screening of eight Be-(d)Cas12j variants (Be-(d)Cas12j-11 to -18) revealed detectable editing efficiencies ranging from 0.52% to 21.06% at *CLTA* target-1 (Fig. 9B-C). Among these, Be-(d)Cas12j-13 demonstrated the highest A-to-G substitution efficiency, identifying it as the most promising candidate (Fig. 7C). To further evaluate its potential, we tested Be-(d)Cas12j-13 and Be-(d)Cas12j-18 across multiple genomic targets (Fig. 7B).

Deep amplicon sequencing revealed variable A-to-G substitution efficiencies across different loci. Notably, Be-(d)Cas12j-13 exhibited the highest A-to-G editing rates at *CLTA* target-1 and *CLTA* target-7, while Be-(d)Cas12j-18 displayed the strongest activity at *AIFM* target-2 (Fig. 7D). Additionally, Be-(d)Cas12j-13 demonstrated a highly specific editing window, with A-to-G conversions predominantly occurring between positions +5 to +10 nt within the target sequence (Fig. 7E). This narrow editing window enhances its precision for gene correction applications while reducing the risk of bystander modifications. These findings establish Be-(d)Cas12j as a promising, compact base-editing tool with potential applications in precision genome engineering and therapeutic gene correction.

## Discussion

CRISPR/Cas systems have revolutionized genome editing, enabling precise modifications for applications in functional genomics, biotechnology, and gene therapy [1,31,32]. However, several challenges limit their broad utility, particularly for *in vivo* applications [33,34]. Chief among these challenges are delivery constraints [35,36], PAM sequence flexibility, and on-target specificity. The need for compact, efficient, and precise genome editors has driven research toward alternative CRISPR systems, with Cas12 family members emerging as promising candidates due to their smaller size when compared to conventional Cas9 nucleases and T-rich PAM sequences [6].

Among the compact Cas12 systems, Cas12j has attracted increasing attention due to its small size and ability to function with short crRNAs [15]. Unlike Cas12f, which requires host RNase-dependent pre-crRNA processing, Cas12j possesses intrinsic crRNA processing activity, simplifying its application [37]. Furthermore, its reliance on 5′-TTN-3′ PAM provides greater targeting flexibility compared to Cas12f, which is constrained to 5′-TTTN-3′ PAMs [6,12]. Despite these advantages, the editing efficiency of Cas12j orthologues in eukaryotic cells remains suboptimal, limiting their translational potential [16,18]. To date, only four of the eleven previously known Cas12j orthologues have shown measurable editing activity in mammalian cells, highlighting the need for further exploration and engineering [17,18].

To address these limitations, we expanded the Cas12j family by conducting HMM-based searches against viral metagenomes, leading to the identification of eight previously uncharacterized Cas12j orthologues (Cas12j-11 to -18). These orthologues, ranging from 718 to 796 AAs, were encoded within CRISPR loci containing 36-bp repeats and 35–38-bp spacers. Structural analysis confirmed that all eight orthologues harbor conserved catalytic residues, validating their classification within the Cas12j family. *In vitro* assays revealed that all eight orthologues exhibit robust catalytic activity, confirming their potential as functional nucleases. To assess their activity in mammalian cells, we codon-optimized these orthologues, expressed them in HEK293T and K-562 cells, and evaluated their ability to generate genome edits at endogenous loci. Among the orthologues tested, Cas12j-12 and Cas12j-18 exhibited the highest editing efficiencies, demonstrating their potential for *in vivo* genome editing applications. While these orthologues showed higher efficiency compared to previously characterized Cas12j variants, their performance remained lower than that of LbCas12a, necessitating further engineering to enhance their efficiency.

To overcome the inherent limitations of Cas12j-mediated editing, we engineered T5 exonuclease-Cas12j (T5Exo-Cas12j) chimeric fusions. T5Exo is a 5′ to 3′ exonuclease [38] known to enhance DNA cleavage and increase the efficiency of Cas effectors. Previous studies have demonstrated that exonuclease fusions with Cas9 and Cas12a increase indel frequency and modulate the repair outcomes, making them attractive candidates for precision genome editing [22,23,25,39]. Our study demonstrates that T5Exo-Cas12j-12 and T5Exo-Cas12j-18 exhibited comparable editing efficiencies to LbCas12a. Importantly, this enhancement was achieved without compromising the compact size of Cas12j, as the T5Exo-Cas12j fusions (1,025-1,103 AAs) remain significantly smaller than LbCas12a (1,228 AAs) and SpCas9 (1,368 AAs) [40]. These findings highlight T5Exo fusion as an effective strategy to improve Cas12j-based genome editing, offering an efficient and size-constrained alternative to existing CRISPR enzymes. Interestingly, target-dependent variations in editing efficiency were observed, consistent with previous reports on Cas-mediated editing in mammalian systems. Notably, Cas12j-18 exhibited consistently high catalytic activity *in vitro* and *in cellula*, particularly when fused to T5Exo, reinforcing its potential as a robust genome editor.

A key discovery in this study was that Cas12j editing activity is significantly enhanced when target DNA begin with a 5′-TAC trinucleotide. While previous studies have reported sequence preferences in CRISPR effectors, our findings reveal that Cas12j editors exhibit a strict requirement for this motif, a feature not observed in Cas9 or Cas12a. To systematically evaluate this effect, we designed multiple crRNAs for targets containing 5′-TAC sequences and compared their editing efficiencies across different genomic targets. The data consistently confirmed that Cas12j editing is most efficient when target DNA starts with TAC, suggesting that this sequence plays a critical role in target recognition and cleavage. The molecular basis of this preference remains poorly understood and warrants further investigation. Future studies should focus on structural analyses of Cas12j-crRNA-DNA interactions, which could uncover the mechanistic basis of TAC dependence and inform rational engineering strategies to further optimize Cas12j activity. Genome-wide analysis revealed that 32.34%–70.7% of open reading frames (ORFs) across different genomes contain TTNTAC motifs, with 44.83% of ORFs in *Homo sapiens* harboring these motifs. This widespread distribution underscores the potential applicability of Cas12j editors for targeted genome engineering.

Beyond conventional DSB-based genome editing, we sought to expand the functional scope of Cas12j by developing Cas12j-based base editors. By fusing TadA8e deaminase to catalytically inactive (d)Cas12j, we created Be-(d)Cas12j, a compact base-editing platform that facilitates A-to-G conversions without inducing DSBs. Among the variants tested, Be-(d)Cas12j-13 (968 AAs) exhibited the highest base-editing efficiency, providing an efficient alternative to Cas9-derived base editors, which are significantly larger. Deep sequencing revealed that Be-(d)Cas12j-13 operates within a narrow editing window (+5 to +10 nt), enhancing its specificity and minimizing off-target modifications. Given its compact size and high specificity, Be-(d)Cas12j represents a promising candidate for therapeutic genome editing, particularly for AAV-mediated *in vivo* applications where vector capacity is a major limitation.

The compact size, enhanced editing efficiency, and programmability of Cas12j editors make them attractive alternatives to conventional CRISPR enzymes. T5Exo-Cas12j fusions provide a highly efficient genome-editing tool, while Be-(d)Cas12j enables precise base editing with minimal collateral damage, expanding the CRISPR toolkit for both gene-disruption and gene-correction applications. Given the demonstrated multiplex-editing capability of T5Exo-Cas12j, these editors could be leveraged for synthetic biology, metabolic engineering, and crop trait enhancement. Additionally, the high target specificity of Be-(d)Cas12j suggests its potential for therapeutic applications requiring precision editing of disease-related genes.

Future efforts should focus on structural studies to elucidate the molecular basis of Cas12j specificity, particularly its TAC sequence dependence, which may inform further optimization strategies. Additionally, engineering approaches aimed at improving crRNA compatibility and broadening PAM recognition will be essential for expanding the utility of Cas12j editors across diverse genomic targets. Finally, optimization of delivery strategies is critical to enhancing *in vivo* efficacy, particularly in animal models and plants, where efficient and precise genome editing remains a major challenge.

### Concluding remarks

Our study expands the CRISPR genome-editing toolkit by identifying novel Cas12j orthologues and demonstrates that their genome editing activity in mammalian cells can be substantially improved through fusion with T5 exonuclease. The resulting T5Exo-Cas12j fusions retain a compact size while reaching editing efficiencies on par with LbCas12a. Additionally, the development of Be-(d)Cas12j base editors enables precise A-to-G conversions, broadening the functional scope of Cas12j systems. A consistent preference of 5′-TAC target sequences emerged as a key determinant of editing activity, offering new avenue for understanding and optimizing Cas12j targeting. Collectively, these innovations establish Cas12j-based editors as compact, efficient, and versatile tools for therapeutic and biotechnological applications.

## STAR METHODS

KEY RESOURCES TABLE.

KEY RESOURCES TABLE
| Reagent or resource | Source | Identifier |
| --- | --- | --- |
| **Experimental models** |  |  |
| HEK293T cell line | Abcam | ab255593 |
| K-562 cell line | RIKEN BRC | RCB1897 |
| *E. coli* BL21(DE3) cells | Thermo Scientific™ | EC0114 |
| One Shot TOP10 Chemically Competent *E. coli* | Invitrogen™ | C404010 |
| **Recombinant DNA** |  |  |
| pMBP-LbCas12a | Addgene | 113431 |
| Mammalian expression vector | Addgene | 194965 |
| crRNA expression plasmid | Addgene | 204639 |
| Additional plasmids and more information | This study | Supplementary Files 1A- 1D |
| **Oligo nucleotides and sequence-based reagents** |  |  |
| crRNA, guide RNA, and primer sequences | This study | Tables S4-S7 |
| **Chemicals, enzymes and other reagents** |  |  |
| HiScribe<sup>®</sup> T7 quick high yield RNA synthesis kit | New England Biolabs (NEB) | E2050 |
| Direct-zol<sup>™</sup> RNA miniprep kit | Zymo Research | R2050 |
| BbsI | NEB | R0539L |
| Terrific broth | IBI scientific | IB49140 |
| IPTG (isopropyl β-D-1-thiogalactopyranoside) | GoldBio | I2481C100 |
| 1X rCutsmart buffer | NEB | B6004S |
| BglI | NEB | R0143L |
| BsaI-HFv2 | NEB | R3733S |
| Proteinase K | Thermo Fisher Scientific | EO0491 |
| 6x purple loading dye | NEB | B7024S |
| Dulbecco's Modified Eagle Medium | Thermo Fisher Scientific | 31966 |
| RPMI 1640 Medium | Gibco<sup>™</sup> | 21875034 |
| Fetal bovine serum | Thermo Fisher Scientific | 10082147 |
| Penicillin-Streptomycin | Thermo Fisher Scientific | 15140122 |
| Gelatin | Sigma-Aldrich | 48723 |
| FuGENE<sup>®</sup> HD Transfection Reagent | Promega | E2312 |
| SF Cell Line 4D-Nucleofector™ X Kit S | Lonza | V4XC-2032 |
| Monarch® genomic DNA extraction kit | NEB | T3010L |
| T7 endonuclease I | NEB | M0302L |
| Phusion™ High-Fidelity DNA Polymerase | NEB | MO530L |
| **Software** |  |  |
| ImageJ | https://imagej.nih.gov/ij/index.html |  |
| GraphPad Prism 10 | https://www.graphpad.com |  |
| BioRender | https://www.biorender.com |  |
| CRISPResso2 (v2.3.2) | https://crispresso2.pinellolab.org |  |
| MAFFT v7.525 | https://mafft.cbrc.jp/alignment/software/source.html |  |
| HMMER v3.4 | http://hmmer.org/ |  |
| CRISPRone | https://omics.informatics.indiana.edu/CRISPRone/ |  |
| CRISPRCasTyper v1.8 | https://crisprcastyper.crispr.dk/ |  |
| AlphaFold2 v2.3.2 | https://github.com/google-deepmind/alphafold |  |
| Foldseek | https://github.com/steineggerlab/foldseek |  |
| CD-HIT v4.8.1 | https://sites.google.com/view/cd-hit |  |
| TrimAI v1.4 | https://github.com/inab/trimal |  |
| IQ-TREE v2.4 | https://github.com/iqtree/iqtree2 |  |

## EXPERIMENTAL MODEL AND STUDY PARTICIPANT DETAILS

Genome editing experiments were conducted using the human wild-type HEK293T cell line (Abcam - ab255593) and K-562 cell line (RIKEN BRC - RCB1897). HEK293T cells were cultured in Dulbecco’s Modified Eagle Medium (Thermo Fisher Scientific - 31966) and K-562 cells in RPMI 1640 Medium (Gibco^TM^ - 21875034), each supplemented with 10% fetal bovine serum (Thermo Fisher Scientific - 10082147), and 1% Penicillin-Streptomycin (Thermo Fisher Scientific - 15140122). All cell cultures were maintained at 37°C in a humidified incubator with 5% CO_2_.

## METHOD DETAILS

### Identification of uncharacterized Cas12j orthologues from metagenome

Putative Cas12j orthologues were identified by constructing a suite of hidden Markov model (HMM) profiles derived from class 2 type V Cas proteins [17]. Each Cas12 subtype (Cas12a, Cas12b, etc.) was separately aligned using MAFFT v7.47 [41], and the resulting multiple sequence alignments were used to build corresponding HMMs with HMMER tools v3.4 [42]. These independents HMM profiles were subsequently combined into a single consolidated database.

Two tiers of the Joint Genome Institute’s (JGI) IMG/VR database v4.1 [19] were then searched for novel Cas12j proteins. First, High-Confidence 5,621,398 viral genomes representing 112,567,455 putative open reading frames (ORFs) were queried using an E-value cutoff of 1e-10. Hits for which the Cas12j HMM model ranked as the top-scoring match were retained. In the second tier, a broader dataset of 15,722,824 viral genomes encompassing 220,799,163 ORFs was queried using the same threshold. Only those ORFs located within 1,000 bp of a predicted CRISPR array, identified by CRISPRone [43], and falling within a typical Cas12j protein length range (700-800 amino acids) were considered for subsequent analysis. CRISPRCasTyper [44] was used to verify the presence of canonical Cas12j genetic cassettes.

### *In silico* domain annotation of identified orthologues

To confirm the structural features of candidate proteins, three-dimensional structure prediction was performed using AlphaFold2 [45]. The crystal structure of Cas12j-3 (PDB: 7ODF) was retrieved, and domain boundaries were identified [20]. These domains were then clipped from the reference structure using a custom Python script and subsequently indexed as a structural database with Foldseek [46]. Candidate proteins were queried against this database for structure-based domain annotation.

### Phylogenetic analysis

We incorporated newly discovered Cas12j variants alongside representative type V Cas proteins obtained from 21 profiles described by Makarova et al., [47]. Sequences within each profile were clustered to 70% identity with CD-HIT v4.8.1 [48], and a single representative from each cluster was selected. Newly identified Cas12j proteins were merged with existing Cas12j references from Pausch et al., work [16]. A multiple sequence alignment was generated using MAFFT v7.525 [41] with the L-INS-I algorithm, iterated 1,000 times, and trimmed with TrimAl v1.4 [49] under default settings to remove poorly aligned regions. Maximum-likelihood phylogeny was constructed using IQ-TREE [50], employing 100 parsimony-based and BIONJ starting trees. ModelFinder [51] determined Q.pfamFR4 as the best-fit protein substitution model according to the Bayesian Information Criterion, and ultrafast bootstrapping with 1,000 replicates was conducted to assess branch support.

### Construction of Cas12j expression plasmids

For *in vitro* studies, genes encoding the eight new Cas12j orthologues (Cas12j-11 to -18) (Supplementary Table S1) were *Escherichia coli* (*E. coli*) codon-optimized, synthesized as gblocks from Integrated DNA Technologies, Inc. (IDT) and cloned into pET28a expression vector under the T7 promoter, independently (Supplementary files S1A). Cas12j-8 protein sequence [18] was *E. coli* codon-optimized, synthesized and cloned into pET28a at BamHI and HindIII restriction sites (Supplementary files S1A) by Twist bioscience. pMBP-LbCas12a (Addgene – 113431) was purified as described in Chen et al., [52]. For mammalian cell culturing experiments, eight Cas12j orthologues (Cas12j-11 to -18), eight T5Exo-Cas12j variants (T5Exo-Cas12j-11 to -18) (Supplementary Table S2) and eight Be-(d)Cas12j base editors (Be-(d)Cas12j-11 to -18) (Supplementary Table S3) were human codon-optimized, synthesized and cloned into the mammalian expression vector under the CMV promoter (Addgene – 194965) (Supplementary files S1B) by GenScript Co., Ltd.

### Construction of crRNA expression plasmids

For the Cas12j mediated *in vitro* assays, the T7 promoter-crRNAs for all the eight Cas12j orthologues were ordered as top and bottom DNA oligos from IDT. The top and bottom oligos were re-annealed and *in vitro* transcribed independently using the HiScribe^®^ T7 quick high yield RNA synthesis kit (New England Biolabs (NEB) – E2050). The synthesized crRNAs were purified using the Direct-zol^TM^ RNA miniprep kit (Zymo Research – R2050). The purified crRNAs were further employed for Cas12j mediated *in vitro* cleavage studies (Supplementary Table S4-S5). All the *in cellula* tested crRNAs were also ordered as top and bottom DNA oligos from IDT. (Supplementary Table S4-S5). For *in cellula* crRNA expression, a plasmid from Addgene-204639 was modified and employed for further cloning. The modified plasmid contains AgeI and BbsI restriction sites after *U6* promoter and self-cleaving HDV ribozyme [53] after the BbsI restriction site (Supplementary files S1D). After the re-annealing of the top and bottom oligos, the annealed dsDNA (encoding crRNA) containing 4-nt overhangs compatible with AgeI at 5′-end and BbsI at 3′-end in the expression plasmid under the *U6* promoter (Supplementary files S1D). After the cloning, the *U6* promoter-crRNA-HDV ribozyme-terminator sequences were PCR amplified using the crRNA forward and crRNA reverse primers (Supplementary Table S6). These PCR amplicons were purified and further co-delivered with Cas12j expression plasmids for gene editing studies.

### Purification of newly identified Cas12j proteins

Protein purification was carried out using a modified version of the protocol described by Wang et al., [54]. Bacterial expression vectors of Cas12j-8 and eight Cas12j variants were independently expressed in *E. coli* BL21(DE3) cells. Each overnight grown *E. coli* BL21(DE3) Cas12j protein expression cultures were inoculated independently into 1-Liter terrific broth medium supplemented with 50 µg/mL kanamycin and grown at 37°C until reaching an OD-600 to 0.4. Followed by IPTG (isopropyl β-D-1-thiogalactopyranoside) induction at a final concentration of 0.1 mM. Cells were harvested after 16 hours post-induction at 18°C by centrifugation at 6,000 × g for 15 minutes at 4°C. The harvested cells were resuspended in a lysis buffer containing a protease inhibitor cocktail and lysed further by sonication. The lysate was clarified by centrifugation at 16,000 × g for 30 minutes at 4°C to remove cell debris and insoluble material. The soluble fractions containing the recombinant Cas12j proteins were collected and applied to an affinity chromatography. For affinity purification, the recombinant Cas12j orthologues were tagged with His-tag for purification. The bound Cas12j orthologues were eluted from the affinity resin using an elution buffer. Later, the proteins were collected and applied onto a cation exchange column, washed with low-salt buffer, and eluted. The eluted proteins were further purified by size-exclusion chromatography on an S200 column (GE Healthcare). After examination by SDS-PAGE electrophoresis, the fractions containing the protein were pooled, concentrated, snap-frozen, and stored at -80°C.

### *In vitro* cleavage assay

For *in vitro* cleavage assay, 5 µM of Cas12j RNP for each orthologue was prepared independently, by mixing an equimolar amount of Cas12j protein (Supplementary Table S1) and respective crRNA (Supplementary Tables S4-S5) in 1X rCutsmart buffer (NEB - B6004S) in a 20 µL volume. The mixture was incubated at room temperature for 20 minutes. Later, 400 nM of the RNP was added to 100 ng of BglI (NEB - R0143L) linearized pJET1.2 and circular pJET1.2 plasmids independently (Supplementary files 1C) and incubated at 37°C for 45 minutes in 1X rCutsmart buffer. For cleavage reactions using circular DNA targets, 0.5 µL of BsaI-HFv2 (NEB - R3733S) was added, and the incubation continued at 37°C for and additional 15 minutes to facilitate the release of the desired DNA fragment. The same protocol was followed for both LbCas12a and Cas12j-8 proteins. Following cleavage, 1 µL of Proteinase K (20 mg/mL; Thermo Fisher Scientific - EO0491) was added to all the cleavage reactions and incubated at 37°C for 20 minutes to inactivate the proteins. The reaction products were then mixed with 6x purple loading dye (NEB - B7024S) and resolved on a 1 % agarose gel.

### Mammalian cell culture and transfection

HEK293T cells were sub-cultured upon reaching approximately 80% confluency. K-562 cells maintained in suspension and passaged upon reaching a density of 1 x 10^6^ cells/mL. For genome editing experiments in HEK293T cells, 24-well plates were pre-coated with 500 µL of 0.1% gelatin solution and incubated at 37°C for 10 minutes. Following the removal of the gelatin, 8 x 10^4^ HEK293T cells were seeded in each well and grown at 37°C for 16-18 hours until approximately 70% confluency was achieved. Cells were then transfected with 250 ng of a Cas12 protein expression plasmid and 250 ng crRNA-encoding amplicon using the FuGENE^®^ HD Transfection Reagent (Promega - E2312), following the manufacturer’s protocol. For genome editing experiments in K-562 cells, 2 x 10^5^ cells were electroporated with 750 ng of Cas12 protein expression plasmid and 350 ng crRNA encoding amplicon using SF Cell Line 4D-Nucleofector^TM^ X Kit S (Lonza - V4XC-2032) with the FF-120 pulse code, according to the manufacturer’s instructions. HEK293T and K-562 cells were harvested after 60-72 hours post-transfection for downstream analysis.

### Genotyping by T7EI assays

To evaluate the genome editing activity of Cas12j-11 to -18, T5Exo-Cas12j-11 to -18, and Be-(d)Cas12j-11 to -18 editors, the extracted genomic DNA from transfected mammalian cells using Monarch^®^ genomic DNA extraction kit (NEB - T3010L). The extracted genomic DNA samples were PCR amplified using respective primer sets mentioned in the Supplementary Table S6. Later the PCR amplicons were subjected to the T7 endonuclease I (T7EI) assay. T7EI mutation detection reactions were conducted using 200 ng of PCR amplicons following the previously described method [55]. The T7EI treated products were resolved on 1.5% agarose gels.

### Multiple genes targeting and genotyping

For the multiple target cleavage, we co-delivered 250 ng of Cas12 protein expression plasmid and four different crRNA encoding amplicons (each 65 ng) using the Fugene^®^ HD transfection kit. Cells were harvested after 60-72 hours of post-transfection. For each target cleavage analysis, the genomic DNA extracted from the co-transfected HEK293T cells. Later, the four targeted regions in the genomic DNA were independently amplified using target specific primers (Supplementary Table S6). Further, T7EI assay conducted for each amplicon independently.

### Deep amplicon sequencing

To further characterize the genome editing efficiencies of different Cas12j variants, we prepared sequencing libraries as following, (i) the targeted loci were amplified using the target specific primers (Supplementary Table S6). [51] 33-nt adapters were attached to the 5′ and 3′ ends of the PCR amplicons from the first step through PCR amplification using target specific primers linked to the 33-nt adapters (Supplementary Table S6). (iii) the PCR products from the second step were barcoded using Illumina index primer pairs. The amplicons were further quantified using a Qubit 2.0 fluorometer then pooled the amplicons in equimolar ratio and prepared the libraries. The amplicon libraries were then subjected to paired-end sequencing on the MiSeq Illumina platform. After Miseq run, the raw reads were subjected to quality control using *fastp* and remove the adapter sequences and low-quality reads. High-fidelity reads were processed with CRISPResso2 (v2.3.2) and quantify genome editing outcomes [56]. To characterize mutation distributions, editing events were categorized based on insertion, deletion, or substitution size, and read counts and frequencies were summed across replicates. This approach enabled a comprehensive assessment of mutation patterns while mitigating technical variability and ensuring consistency in data interpretation.

## QUANTIFICATION AND STATISTICAL ANALYSIS

DNA bands on Agarose gels were quantified using ImageJ software. All statistical analysis was performed using GraphPad Prism 10 software. Data represent mean with standard deviation [4]. As indicated in the figure legends, various statistical tests were employed to calculate *P*-values, . *P*-values indicated are: ns is not significant (0.1234), * (0.0332), ** (0.0021), *** (0.0002), **** (<0.0001).

## RESOURCE AVAILABILITY

### Lead contact

Requests for further information and resources should be directed to, and will be fulfilled by, the lead contact, Magdy Mahfouz.

### Data availability

NGS raw data is available in the NCBI database with BioProject ID PRJNA1236335. Nucleotide sequences of *E. coli* codon optimized Cas12j-11 to -18, human codon optimized Cas12j-11 to - 18, T5Exo-Cas12j-1 to -18 and Be-(d)Cas12j-11 to -18 employed in this study are available in the NCBI database with nucleotide accessions PV362975 to PV363006. All other data are provided in the main text and supporting information.

## Author contributions

G.S.R., W.J., M.A. and M.M. designed the research; A.K., A.E., G.S.R., and W.J. conducted the bioinformatics studies; G.S.R., W.J., Q.W., A.S., Maazallah M. and A.G. performed the experiments; G.S.R., W.J., M.A. and M.M. analyzed the data. M.M, W.J. and G.S.R wrote the manuscript with inputs from all authors.

## Acknowledgements

We would like to thank all the members of the genome engineering and synthetic biology laboratory for their useful discussions and consistent help. Schematic figures were created with BioRender (biorender.com).

## Technology readiness

The compact Cas12j editors presented here are moving from proof-of-concept toward practical readiness. By creating T5 exonuclease-Cas12j fusions, we substantially improved their performance, achieving efficiencies on par with the LbCas12a in mammalian cells. A remarkable finding is the strict requirement for a 5′-TAC tri-nucleotide downstream of the PAM, which transforms these editors from modest performers into robust tools. This sequence preference, while initially a limitation, can be harnessed as a powerful design principle to ensure predictable activity across disease relevant and synthetic targets. The development of Be-(d)Cas12j base editors further broadens the scope, enabling precise A-to-G substitutions within a compact format well suited for AAV and non-viral delivery. Currently, the technology stands at an advanced laboratory validation stage, functional in multiple human cell lines and across clinically relevant loci, but not yet tested *in vivo*. With continued optimization of delivery and broader PAM and sequence compatibility, these miniature editors are poised to impact therapeutic genome editing, synthetic biology, and plant engineering.

## References

1. Liu, G. et al. (2022) The CRISPR-Cas toolbox and gene editing technologies. Molecular Cell 82, 333–347. 10.1016/j.molcel.2021.12.002

2. Wang, J.Y. and Doudna, J.A. (2023) CRISPR technology: A decade of genome editing is only the beginning. Science 379, eadd8643. doi:10.1126/science.add8643

3. Zhang, F. and Huang, Z. (2022) Mechanistic insights into the versatile class II CRISPR toolbox. Trends in Biochemical Sciences 47, 433–450. 10.1016/j.tibs.2021.11.007

4. Demirci, S. et al. (2022) Advances in CRISPR Delivery Methods: Perspectives and Challenges. The CRISPR Journal 5, 660–676. 10.1089/crispr.2022.0051

5. Aquino-Jarquin, G. (2023) Genome and transcriptome engineering by compact and versatile CRISPR-Cas systems. Drug Discovery Today 28, 103793. 10.1016/j.drudis.2023.103793

6. Tang, N. and Ji, Q. (2024) Miniature CRISPR-Cas12 Systems: Mechanisms, Engineering, and Genome Editing Applications. ACS Chemical Biology 19, 1399–1408. 10.1021/acschembio.4c00247

7. Wu, H. et al. (2024) Advances in miniature CRISPR-Cas proteins and their applications in gene editing. Archives of Microbiology 206, 231. 10.1007/s00203-024-03962-0

8. Truong, D.-J.J. et al. (2015) Development of an intein-mediated split–Cas9 system for gene therapy. Nucleic Acids Research 43, 6450–6458. 10.1093/nar/gkv601

9. Chew, W.L. et al. (2016) A multifunctional AAV–CRISPR–Cas9 and its host response. Nature Methods 13, 868–874. 10.1038/nmeth.3993

10. Yin, H. et al. (2016) Therapeutic genome editing by combined viral and non-viral delivery of CRISPR system components in vivo. Nature Biotechnology 34, 328–333. 10.1038/nbt.3471

11. Diaz-Riascos, Z.V. et al. (2019) Expression and Role of MicroRNAs from the miR-200 Family in the Tumor Formation and Metastatic Propensity of Pancreatic Cancer. Molecular Therapy - Nucleic Acids 17, 491–503. 10.1016/j.omtn.2019.06.015

12. Wu, Z. et al. (2021) Programmed genome editing by a miniature CRISPR-Cas12f nuclease. Nature Chemical Biology 17, 1132–1138. 10.1038/s41589-021-00868-6

13. Sharrar, A. et al. (2023) Discovery and Characterization of Novel Type V Cas12f Nucleases with Diverse Protospacer Adjacent Motif Preferences. The CRISPR Journal 6, 350–358. 10.1089/crispr.2023.0006

14. Kong, X. et al. (2023) Engineered CRISPR-OsCas12f1 and RhCas12f1 with robust activities and expanded target range for genome editing. Nature Communications 14, 2046. 10.1038/s41467-023-37829-7

15. Nguyen, G.T. et al. (2022) Miniature CRISPR-Cas12 endonucleases – Programmed DNA targeting in a smaller package. Current Opinion in Structural Biology 77, 102466. 10.1016/j.sbi.2022.102466

16. Pausch, P. et al. (2020) CRISPR-CasΦ from huge phages is a hypercompact genome editor. Science 369, 333–337. doi:10.1126/science.abb1400

17. Duan, Z. et al. (2023) Molecular basis for DNA cleavage by the hypercompact Cas12j-SF05. Cell Discovery 9, 117. 10.1038/s41421-023-00612-5

18. Wang, Y. et al. (2023) A highly specific CRISPR-Cas12j nuclease enables allele-specific genome editing. Science Advances 9, eabo6405. doi:10.1126/sciadv.abo6405

19. Camargo, A.P. et al. (2023) IMG/VR v4: an expanded database of uncultivated virus genomes within a framework of extensive functional, taxonomic, and ecological metadata. Nucleic Acids Research 51, D733–D743. 10.1093/nar/gkac1037

20. Carabias, A. et al. (2021) Structure of the mini-RNA-guided endonuclease CRISPR-Cas12j3. Nature Communications 12, 4476. 10.1038/s41467-021-24707-3

21. Pausch, P. et al. (2021) DNA interference states of the hypercompact CRISPR–CasΦ effector. Nature Structural & Molecular Biology 28, 652–661. 10.1038/s41594-021-00632-3

22. Wu, Y. et al. (2020) Improving FnCas12a Genome Editing by Exonuclease Fusion. The CRISPR Journal 3, 503–511. 10.1089/crispr.2020.0073

23. Zhang, Q. et al. (2020) Fusing T5 exonuclease with Cas9 and Cas12a increases the frequency and size of deletion at target sites. Science China Life Sciences 63, 1918–1927. 10.1007/s11427-020-1671-6

24. Han, D., et al. (2023) Development of miniature base editors using engineered IscB nickase. Nature Methods 20, 1029–1036. 10.1038/s41592-023-01898-9

25. Liang, Z. et al. (2023) Addition of the T5 exonuclease increases the prime editing efficiency in plants. Journal of Genetics and Genomics 50, 582–588. 10.1016/j.jgg.2023.03.008

26. Bae, S. et al. (2014) Cas-OFFinder: a fast and versatile algorithm that searches for potential off-target sites of Cas9 RNA-guided endonucleases. Bioinformatics 30, 1473–1475. 10.1093/bioinformatics/btu048

27. Kang, S.-H. et al. (2020) Prediction-based highly sensitive CRISPR off-target validation using target-specific DNA enrichment. Nature Communications 11, 3596. 10.1038/s41467-020-17418-8

28. Rees, H.A. and Liu, D.R. (2018) Base editing: precision chemistry on the genome and transcriptome of living cells. Nature Reviews Genetics 19, 770–788. 10.1038/s41576-018-0059-1

29. Anzalone, A.V., et al. (2020) Genome editing with CRISPR–Cas nucleases, base editors, transposases and prime editors. Nature Biotechnology 38, 824–844. 10.1038/s41587-020-0561-9

30. Xiao, Y.-L., et al. (2024) An adenine base editor variant expands context compatibility. Nature Biotechnology 42, 1442–1453. 10.1038/s41587-023-01994-3

31. Katti, A. et al. (2022) CRISPR in cancer biology and therapy. Nature Reviews Cancer 22, 259–279. 10.1038/s41568-022-00441-w

32. Wang, S.-W. et al. (2022) Current applications and future perspective of CRISPR/Cas9 gene editing in cancer. Molecular Cancer 21, 57. 10.1186/s12943-022-01518-8

33. Chen, M. et al. (2019) CRISPR-Cas9 for cancer therapy: Opportunities and challenges. Cancer Letters 447, 48–55. 10.1016/j.canlet.2019.01.017

34. van Haasteren, J., et al. (2020) The delivery challenge: fulfilling the promise of therapeutic genome editing. Nature Biotechnology 38, 845–855. 10.1038/s41587-020-0565-5

35. Wang, D. et al. (2020) CRISPR-Based Therapeutic Genome Editing: Strategies and In Vivo Delivery by AAV Vectors. Cell 181, 136–150. 10.1016/j.cell.2020.03.023

36. Taha, E.A. et al. (2022) Delivery of CRISPR-Cas tools for in vivo genome editing therapy: Trends and challenges. Journal of Controlled Release 342, 345–361. 10.1016/j.jconrel.2022.01.013

37. Harrington, L.B. et al. (2018) Programmed DNA destruction by miniature CRISPR-Cas14 enzymes. Science 362, 839–842. doi:10.1126/science.aav4294

38. Garforth, S.J. and Sayers, J.R. (1997) Structure-Specific DNA Binding by Bacteriophage T5 5′→3′ Exonuclease. Nucleic Acids Research 25, 3801–3807. 10.1093/nar/25.19.3801

39. Han, B. et al. (2022) ErCas12a and T5exo-ErCas12a Mediate Simple and Efficient Genome Editing in Zebrafish. Biology 11, 411

40. Swarts, D.C. and Jinek, M. (2018) Cas9 versus Cas12a/Cpf1: Structure–function comparisons and implications for genome editing. WIREs RNA 9, e1481. 10.1002/wrna.1481

41. Katoh, K. and Standley, D.M. (2013) MAFFT Multiple Sequence Alignment Software Version 7: Improvements in Performance and Usability. Molecular Biology and Evolution 30, 772–780. 10.1093/molbev/mst010

42. Johnson, L.S. et al. (2010) Hidden Markov model speed heuristic and iterative HMM search procedure. BMC Bioinformatics 11, 431. 10.1186/1471-2105-11-431

43. Zhang, Q. and Ye, Y. (2017) Not all predicted CRISPR–Cas systems are equal: isolated cas genes and classes of CRISPR like elements. BMC Bioinformatics 18, 92. 10.1186/s12859-017-1512-4

44. Russel, J. et al. (2020) CRISPRCasTyper: Automated Identification, Annotation, and Classification of CRISPR-Cas Loci. The CRISPR Journal 3, 462–469. 10.1089/crispr.2020.0059

45. Yang, Z. et al. (2023) AlphaFold2 and its applications in the fields of biology and medicine. Signal Transduction and Targeted Therapy 8, 115. 10.1038/s41392-023-01381-z

46. van Kempen, M., et al. (2024) Fast and accurate protein structure search with Foldseek. Nature Biotechnology 42, 243–246. 10.1038/s41587-023-01773-0

47. Makarova, K.S. et al. (2020) Evolutionary classification of CRISPR–Cas systems: a burst of class 2 and derived variants. Nature Reviews Microbiology 18, 67–83. 10.1038/s41579-019-0299-x

48. Li, W. and Godzik, A. (2006) Cd-hit: a fast program for clustering and comparing large sets of protein or nucleotide sequences. Bioinformatics 22, 1658–1659. 10.1093/bioinformatics/btl158

49. Capella-Gutiérrez, S. et al. (2009) trimAl: a tool for automated alignment trimming in large-scale phylogenetic analyses. Bioinformatics 25, 1972–1973. 10.1093/bioinformatics/btp348

50. Minh, B.Q. et al. (2020) IQ-TREE 2: New Models and Efficient Methods for Phylogenetic Inference in the Genomic Era. Molecular Biology and Evolution 37, 1530–1534. 10.1093/molbev/msaa015

51. Kalyaanamoorthy, S. et al. (2017) ModelFinder: fast model selection for accurate phylogenetic estimates. Nature Methods 14, 587–589. 10.1038/nmeth.4285

52. Chen, J.S. et al. (2018) CRISPR-Cas12a target binding unleashes indiscriminate single-stranded DNase activity. Science 360, 436–439. 10.1126/science.aar6245

53. Ferré-D’Amaré, A.R. and Scott, W.G. (2010) Small self-cleaving ribozymes. Cold Spring Harb Perspect Biol 2, a003574. 10.1101/cshperspect.a003574

54. Wang, Q. et al. (2024) Fusion of FokI and catalytically inactive prokaryotic Argonautes enables site-specific programmable DNA cleavage. J Biol Chem 300, 107720. 10.1016/j.jbc.2024.107720

55. Reyon, D. et al. (2012) FLASH assembly of TALENs for high-throughput genome editing. Nature Biotechnology 30, 460–465. 10.1038/nbt.2170

56. Clement, K. et al. (2019) CRISPResso2 provides accurate and rapid genome editing sequence analysis. Nature Biotechnology 37, 224–226. 10.1038/s41587-019-0032-3

