## Supplementary material for "Characterization and engineering of highly efficient Cas12j genome editors": Supporting information.docx

**Supplementary Figures**

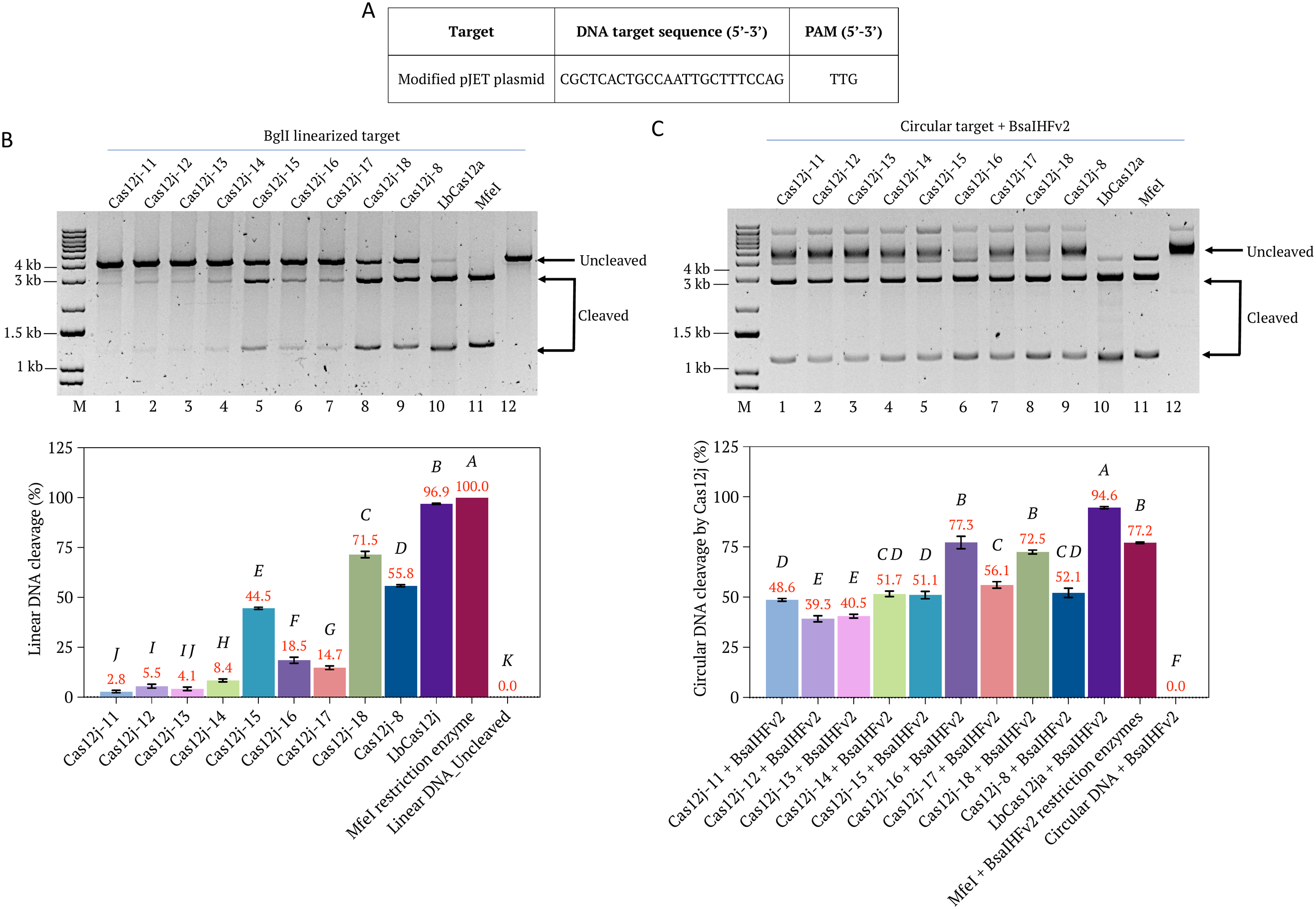

**Supplementary Figure S1.** Comparison of different Cas12j variants mediated *in vitro* cleaved dsDNA. **(A)** Protospacer sequence and PAM sequences of *in vitro* tested target DNA. **(B)** Top panel showing the gel image of targeted cleavage of the BglI-linearized DNA by different Cas12j orthologues (Lanes 1-8), Cas12j-8 (Lane 9), and LbCas12a (Lane 10). Lane 11 is the MfeI treated DNA target and Lane 12 is untreated DNA as a negative control. Lower panel showing the cleavage efficiencies of different Cas12j proteins and LbCas12a. Gels are quantified using ImageJ software. Percentages were calculated by dividing the ImageJ-quantified intensity of the uncleaved band for each sample by the sum of the uncleaved and cleaved band intensities of the same sample, then multiplying the result by 100. Data represent mean with SD (n = 3 independent experiments). Statistical analysis was performed using one-way ANOVA followed by Tukey’s multiple comparisons test. Bars labeled with different letters are significantly different (*P* < 0.05). Bars sharing the same letter are not significantly different (*P* > 0.05). **(C)** Top panel showing the gel image of targeted cleavage of the circular pJET plasmid by different Cas12j orthologues (Lanes 1-8), Cas12j-8 (Lane 9), and LbCas12a (Lane 10). Lane 11 is the MfeI + BsaIHFv2 treated circular DNA target and Lane 12 is circular DNA treated with BsaIHFV2 as a negative control. Circular DNA was initially treated with Cas12j proteins independently. Later BsaIHFv2 restriction enzyme added to the total reaction volume to facilitate the fragment release. Lower panel showing the cleavage efficiencies of different Cas12j proteins and LbCas12a. Gels are quantified using ImageJ software. Percentages are calculated as mentioned in the (B). Data represent mean with SD (n = 3 independent experiments). *P*-values are calculated as mentioned in the (B).

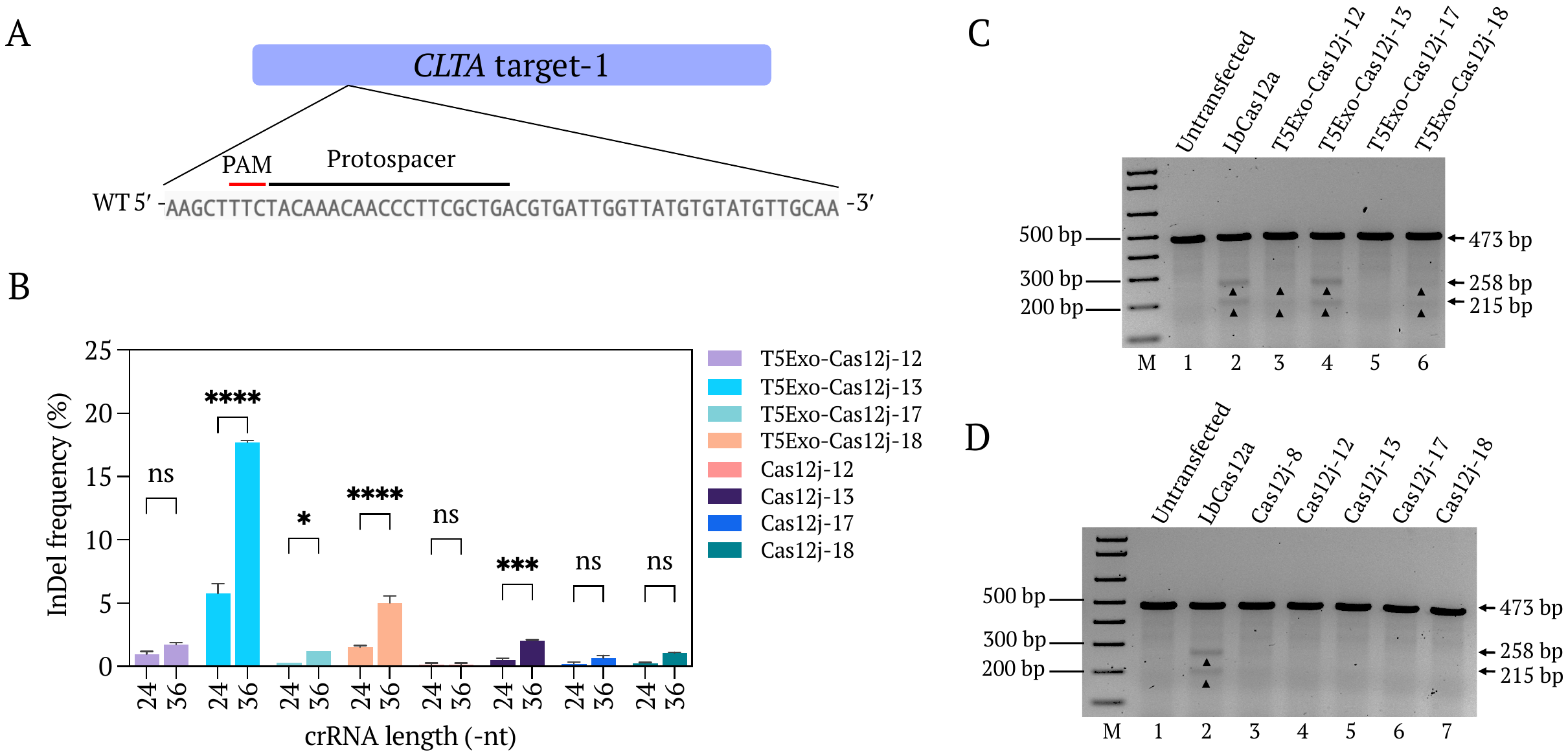

**Supplementary Figure S2.** Activity of different Cas12j and T5Exo-Cas12j variants using mature crRNA (24-nt) and pre-processed crRNA (36-nt) on *CLTA* target-1 genomic region. **(A)** Gene map showing the *CLTA* target-1 seed sequence. PAM is in the red color, bold is the seed sequence. **(B)** The indel frequency of Cas12j and T5Exo-Cas12j variants guided by mature crRNA (24-nt) in HEK293T cells at *CLTA* target-1. The InDel frequencies are compared with pre-processed crRNA (36-nt) mediated genome editing at same locus and using same proteins. Data are shown in mean with standard deviation of two independent biological experiments. *P*-values are calculated using a two-way ANOVA, Tukey test (*P*-values indicated are: ns is not significant (0.1234), * (0.0332), ** (0.0021), *** (0.0002), **** (<0.0001)). **(C)** Gel image showing the T7EI assay for the evaluation of genome editing activity using mature crRNA with LbCas12a (Lane 2), and T5Exo-Cas12j-12, -13, -17, -18 (Lanes 3-6) in HEK293T cells at *CLTA* target-1 region, independently. Lane 1 is the T7EI of the DNA from untransfected cells used as negative control. **(D)** Gel image showing the T7EI assay for the evaluation of genome editing activity using mature crRNA with LbCas12a (Lane 2), Cas12j-8 (Lane 3), and Cas12j-12, -13, -17, -18 (Lanes 3-6) in HEK293T cells at *CLTA* target-1 region, independently. Lane 1 is the T7EI of the DNA from untransfected cells used as negative control. Lane M is the 1kb plus marker.

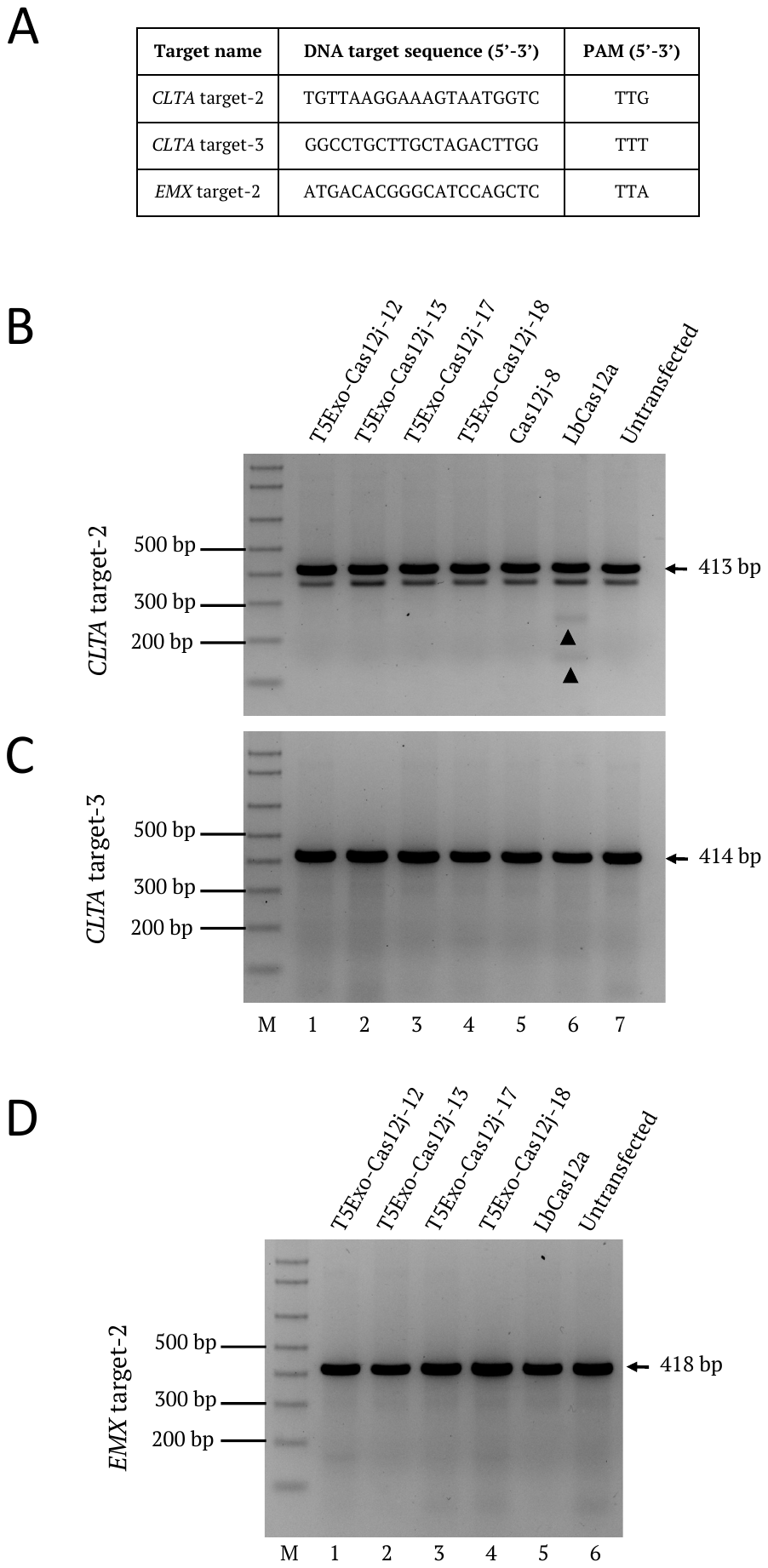

**Supplementary Figure S3.** Testing additional targets with different PAM sequences using T5Exo-Cas12j fusion proteins *in vivo*. **(A)** Table showing the target sequences and the PAM sequences. **(B)** Gel image showing the T7EI assay for the evaluation of genome editing activity using T5Exo-Cas12j-12, -13, -17 -18 (Lanes 1-4), Cas12j-8 (Lane 5) and LbCas12a (Lane 6), in HEK293T cells at *CLTA* target-2 region, independently. Lane 7 is the T7EI of the DNA from untransfected cells used as negative control. Lane M is the 1kb plus marker. **(C)** Gel image showing the T7EI assay for the evaluation of genome editing activity using T5Exo-Cas12j-12, -13, -17 -18 (Lanes 1-4), Cas12j-8 (Lane 5) and LbCas12a (Lane 6), in HEK293T cells at *CLTA* target-3 region, independently. Lane 7 is the T7EI of the DNA from untransfected cells used as negative control. **(D)** Gel image showing the T7EI assay for the evaluation of genome editing activity using T5Exo-Cas12j-12, -13, -17 -18 (Lanes 1-4), and LbCas12a (Lane 5) in HEK293T cells at *EMX* target-2 region, independently. Lane 6 is the T7EI of the DNA from untransfected cells used as negative control. Lane M is the 1kb plus marker.

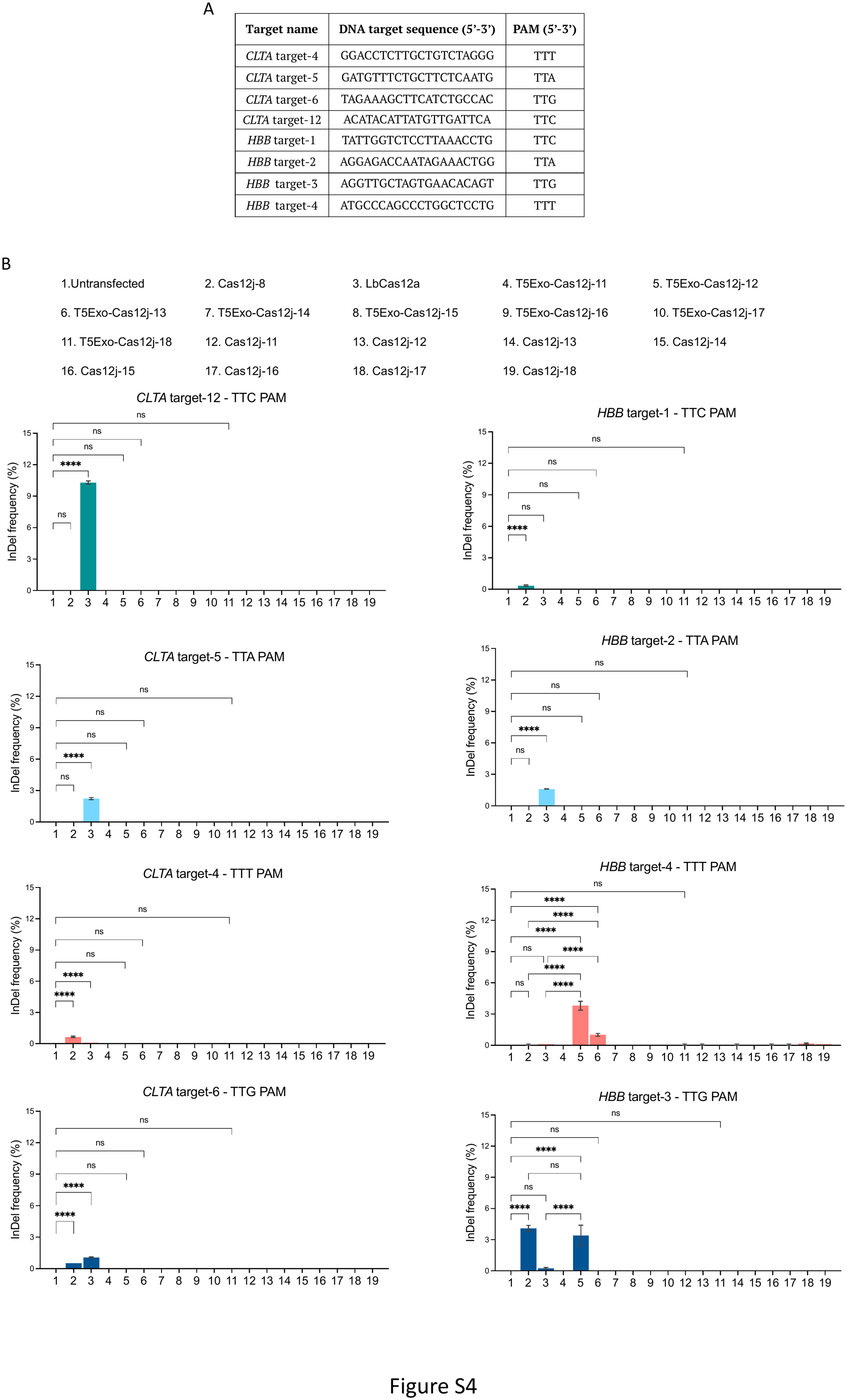

**Supplementary Figure S4.** Gene editing activity of Cas12j and T5Exo-Cas12j fusion proteins on different targets in mammalian cell lines. **(A)** Table showing the target sequences and the different PAM sequences. **(B)** Deep amplicon sequencing data showing the variable indel efficiencies at *CLTA* target-4, -5, -6, -12, *HBB* target-1, -2, -3, -4 regions using different T5Exo-Cas12j and Cas12j variants. Data are shown in mean with standard deviation of two independent biological replicates. *P*-values are calculated using a one-way ANOVA, Bonferroni test (*P*-values indicated are: ns is not significant (0.1234), * (0.0332), ** (0.0021), *** (0.0002), **** (<0.0001)).

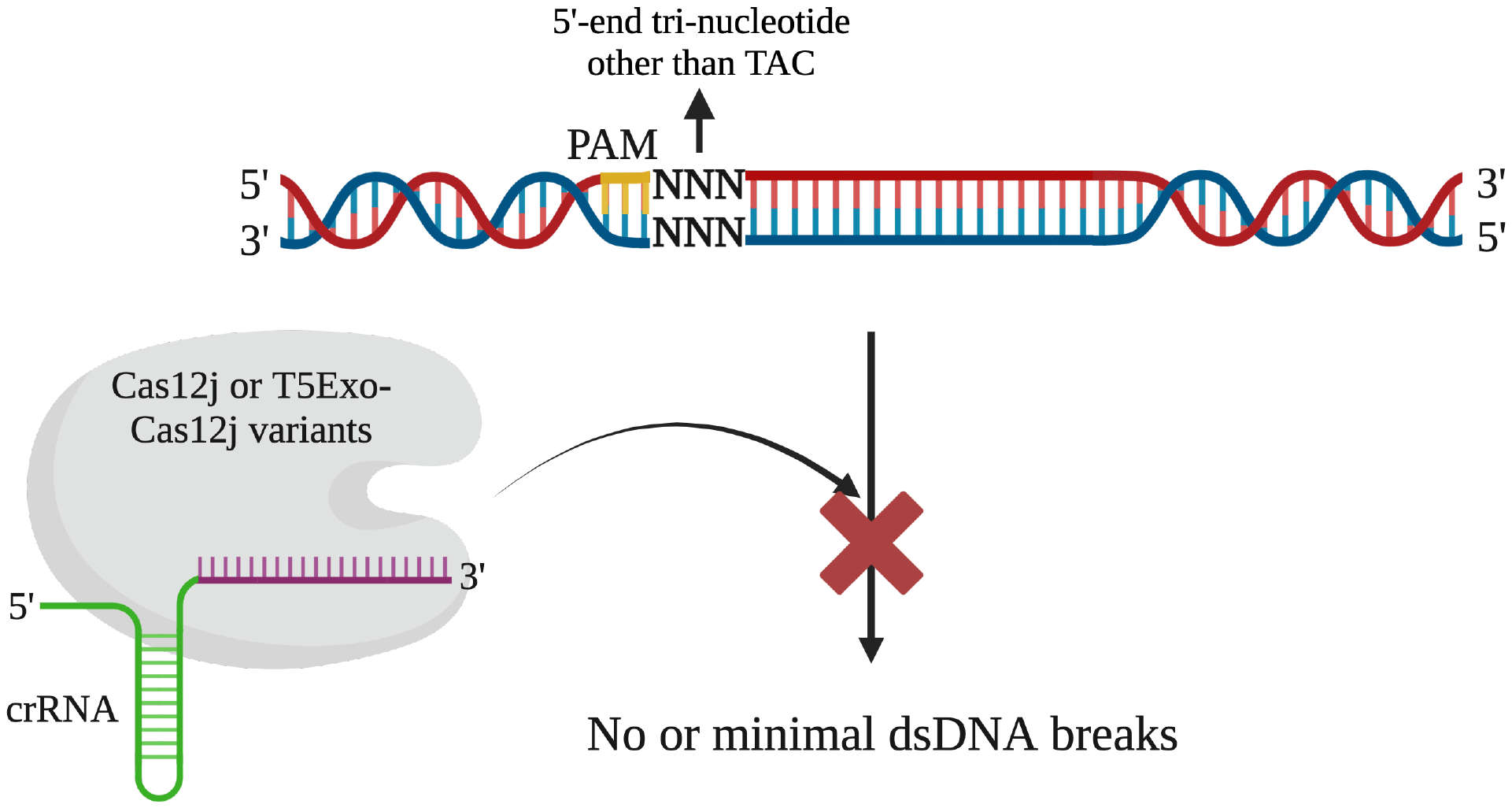

**Supplementary Figure S5.** Representation of Cas12j and T5Exo-Cas12j variants mediated editing of targets containing random 5′-end tri-nucleotide sequences.

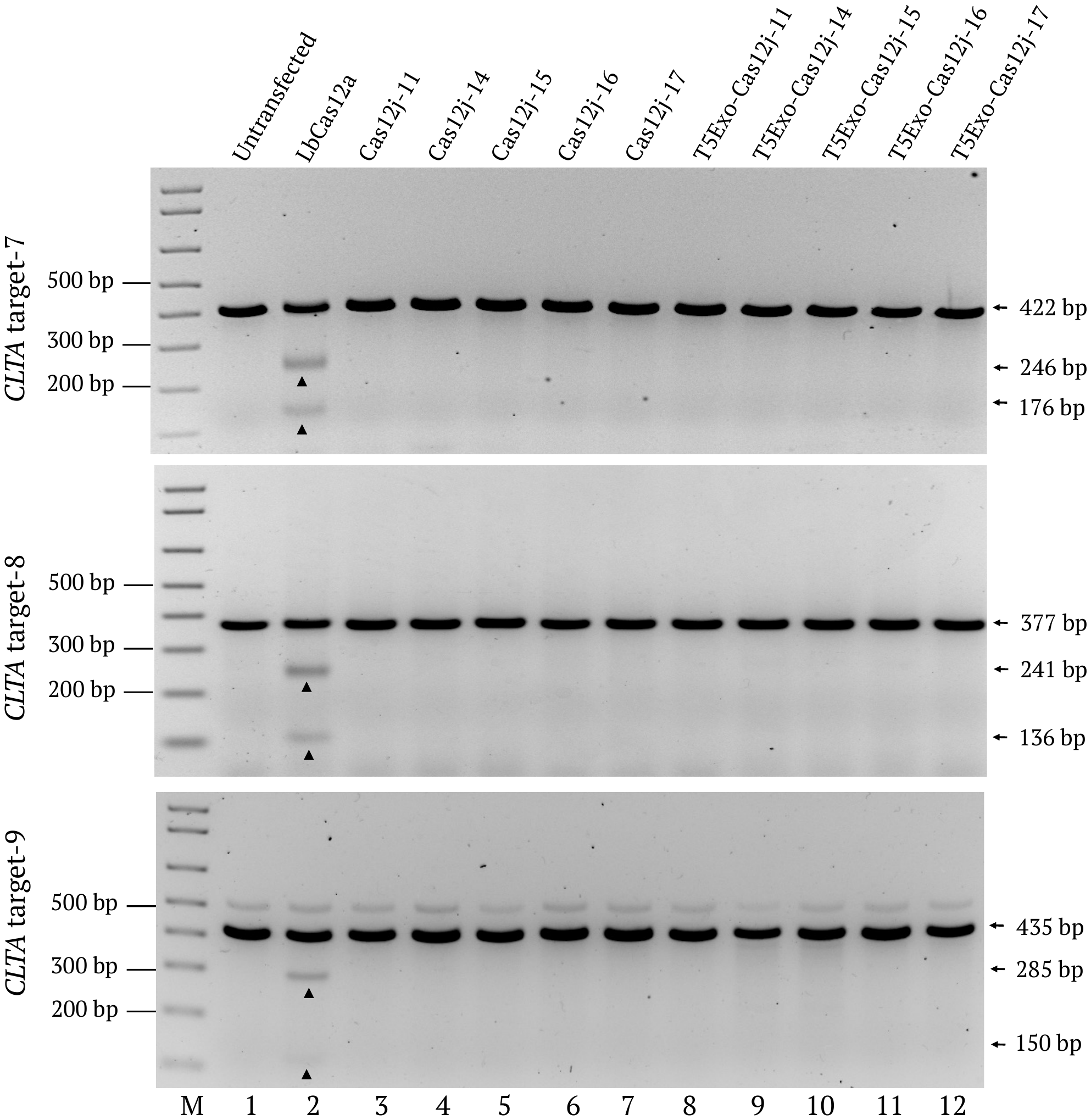

**Supplementary Figure S6.** Gene editing activity of Cas12j and T5Exo-Cas12j fusion proteins on different targets in mammalian cells containing 5′-end TAC tri-nucleotide sequence. Gels showing the T7EI assay results of the genome editing activity using LbCas12a (Lane 2), Cas12j-11, -14, -15, -16, -17 (Lanes 3-7) and T5Exo-Cas12j-11, -14, -15, -16, -17 (Lanes 8-12) in HEK293T cells at *CLTA* target-7, -8, -9 regions, independently. Lane 1 is the T7EI for the DNA from untransfected cells used as negative control. Lane M is the 1kb plus marker.

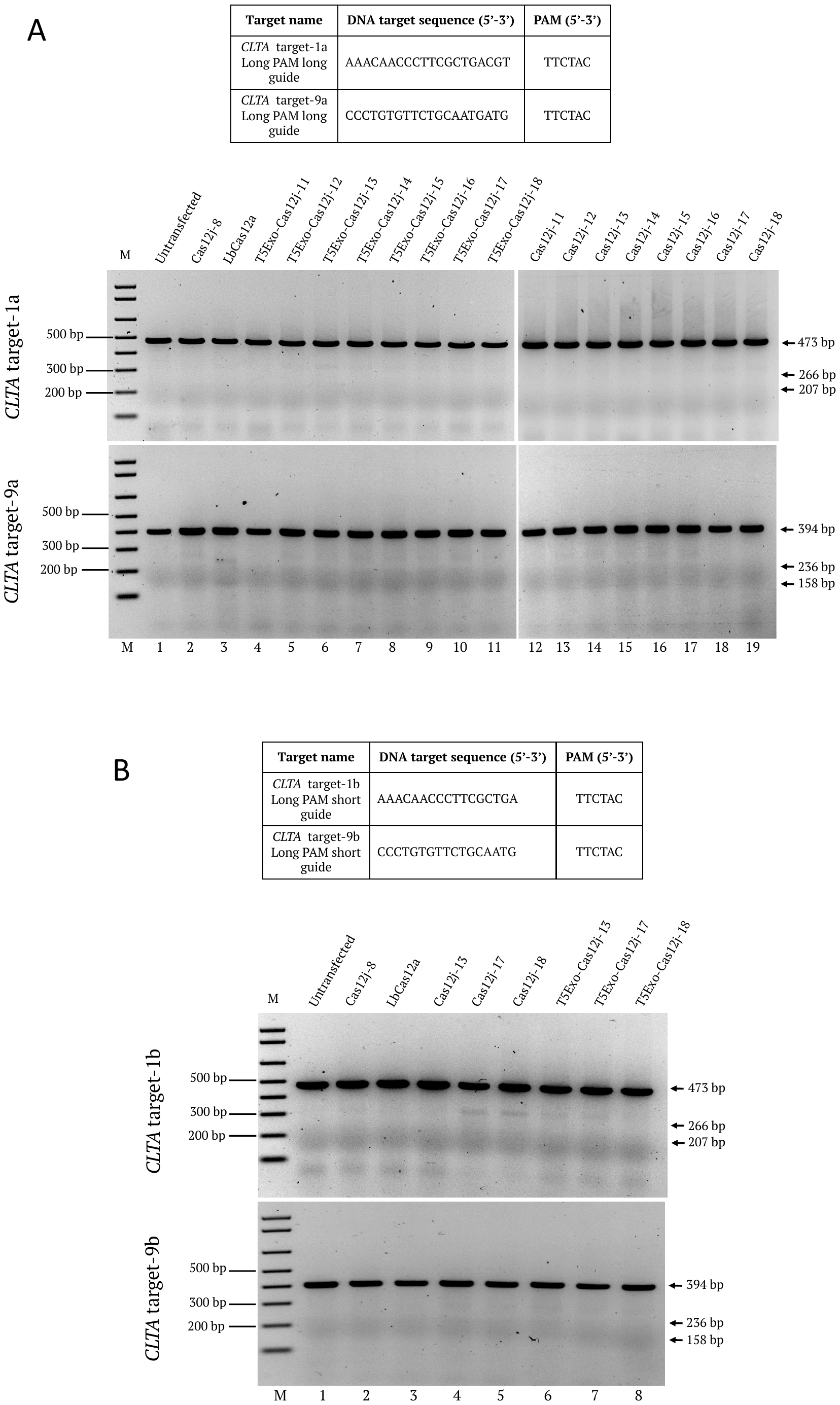

**Supplementary Figure S7.** Genome editing activity of different Cas12j and T5Exo-Cas12j variants on targets containing TTCTAC PAM sequence using different lengths of crRNAs. **(A)** Table showing the target sequences and the PAM sequences. Gel images showing the T7EI assay results of the genome editing activity using Cas12j-8 (Lane 2), LbCas12a (Lane 3), T5Exo-Cas12j-11 to -18 (Lanes 4-11) and Cas12j-11 to -18 (Lanes 12-19) in HEK293T cells at *CLTA* target-1a and *CLTA* target-9a regions, independently. Lane 1 is the T7EI of the DNA from untransfected cells used as negative control. Lane M is the 1kb plus marker. **(B)** Table showing the truncated (17-nt) target sequences and the PAM sequences. Gel images showing the T7EI assay results of the genome editing activity using Cas12j-8 (Lane 2), LbCas12a (Lane 3), Cas12j-13, -17, -18 (Lanes 4-6) and T5Exo-Cas12j-13, -17, -18 (Lanes 7-9) in HEK293T cells at *CLTA* target-1b and *CLTA* target-9b regions, independently. Lane 1 is the T7EI of the DNA from untransfected cells used as negative control. Lane M is the 1kb plus marker.

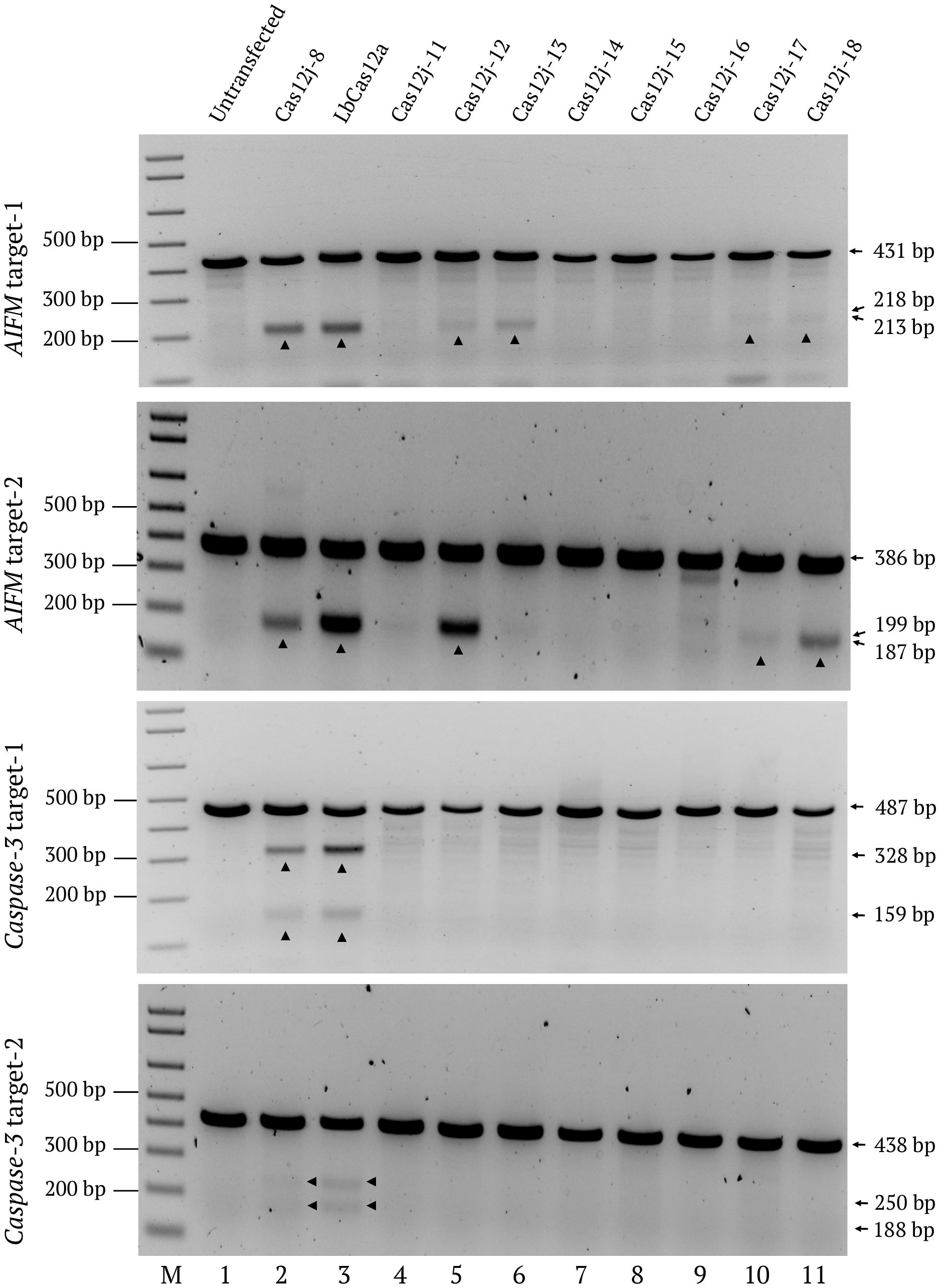

**Supplementary Figure S8.** Gene editing activity of Cas12j and T5Exo-Cas12j fusion proteins on disease related genes in mammalian cells using crRNAs with 5′-end TAC tri-nucleotide sequence. Gels showing the T7EI assay results of the genome editing activity using Cas12j-8 (Lane 2), LbCas12a (Lane 3), and Cas12j-11 to -18 (Lanes 4-11) in HEK293T cells at *AIFM* target-1, -2, and *Caspase3* target-1, -2 regions, independently. Lane 1 is the T7EI of the DNA from untransfected cells used as negative control. Lane M is the 1kb plus marker.

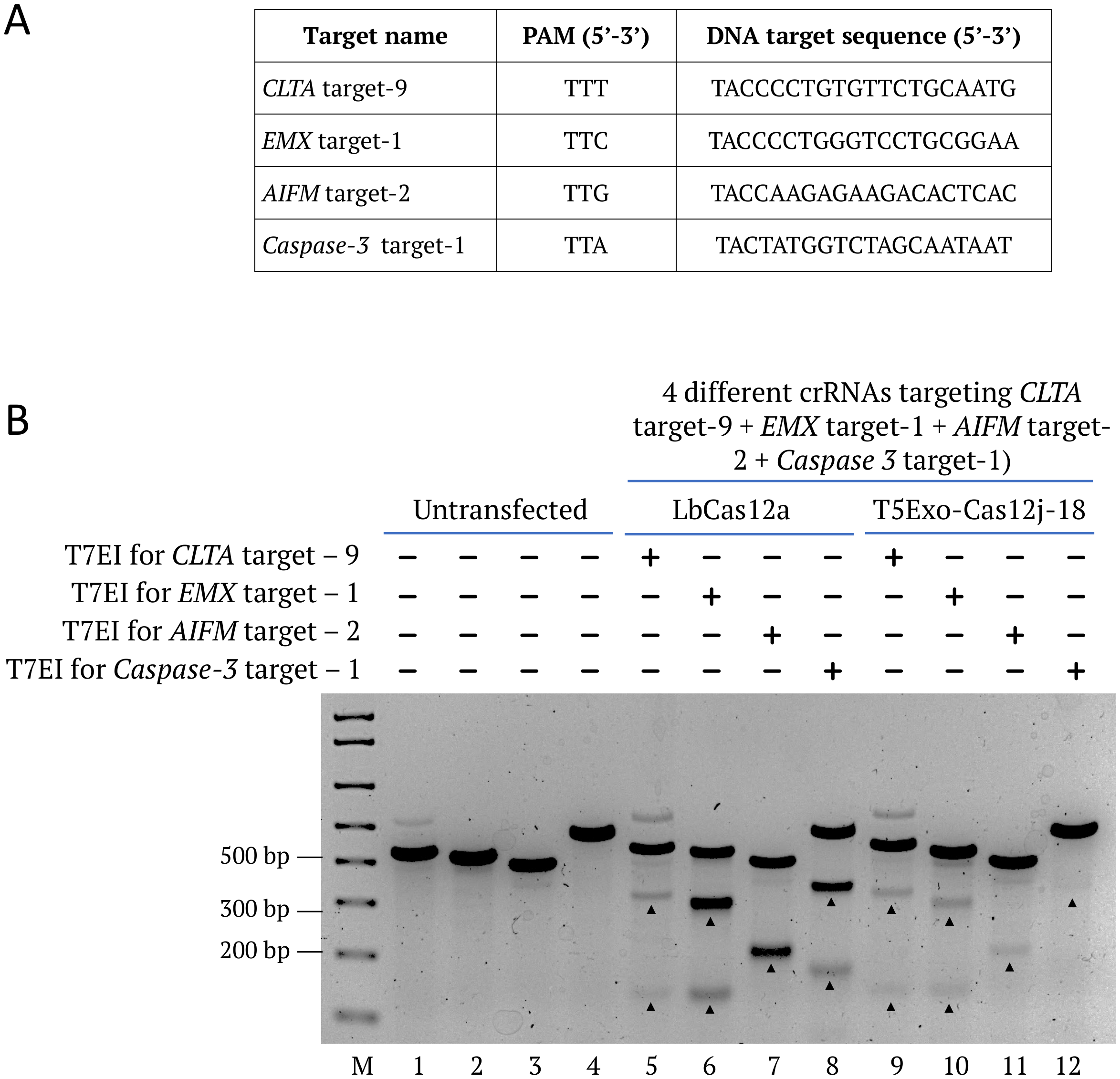

**Supplementary Figure S9.** Co-delivery of multiple crRNAs and gene editors for the multiple target cleavage. **(A)** Table showing the target and PAM sequences of four different genes. **(B)** Gel image showing the T7EI results of four different targets *CLTA* target-9, *EMX* target-1, *AIFM* target-2 and *Caspase-3* target-1 edited simultaneously by LbCas12a (Lanes 5-8) and T5Exo-Cas12j-18 (Lanes 9-12). Lanes 1-4 are the T7EI of the DNA from untransfected cells used as negative control. Lane M is the 1kb plus marker.

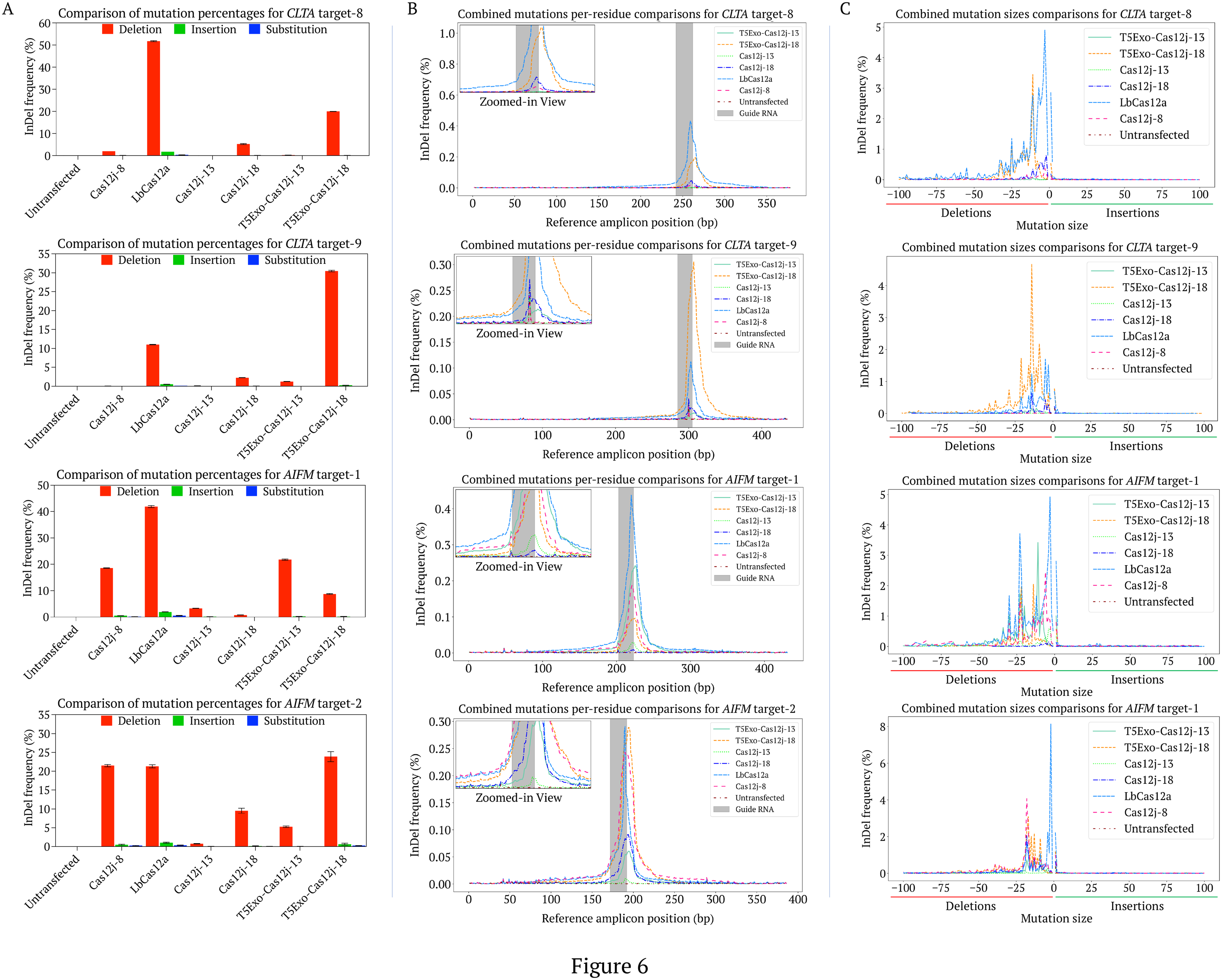

**Supplementary Figure S10.** Types of edits and distribution of edits generated by different Cas12j and T5Exo-Cas12j fusion proteins on different genes in HEK293T cells. **(A)** Comparison of different mutation types generated by Cas12j-8, LbCas12a, Cas12j-13, -18 and T5Exo-Cas12j-13, -18 on *CLTA* target-8, -9, *AIFM* target-1, -2. Data represent mean with SD (n = 2 biologically independent replicates). **(B)** The distribution of indels around the guide RNA of *CLTA* target-8, -9, *AIFM* target-1, -2 generated by Cas12j-8, LbCas12a, Cas12j-13, -18 and T5Exo-Cas12j-13, -18. **(C)** The mutation sizes generated by Cas12j-8, LbCas12a, Cas12j-13, -18 and T5Exo-Cas12j-13, -18 on *CLTA* target-8, -9, and *AIFM* target-1, -2. On X-axis 0 to -100 indicates the different deletion sizes and 0 to 100 indicates the insertion sizes. Untransfected indicates the NGS sequencing results of DNA from untransfected cells used as negative control. All data were normalized by two replicates.

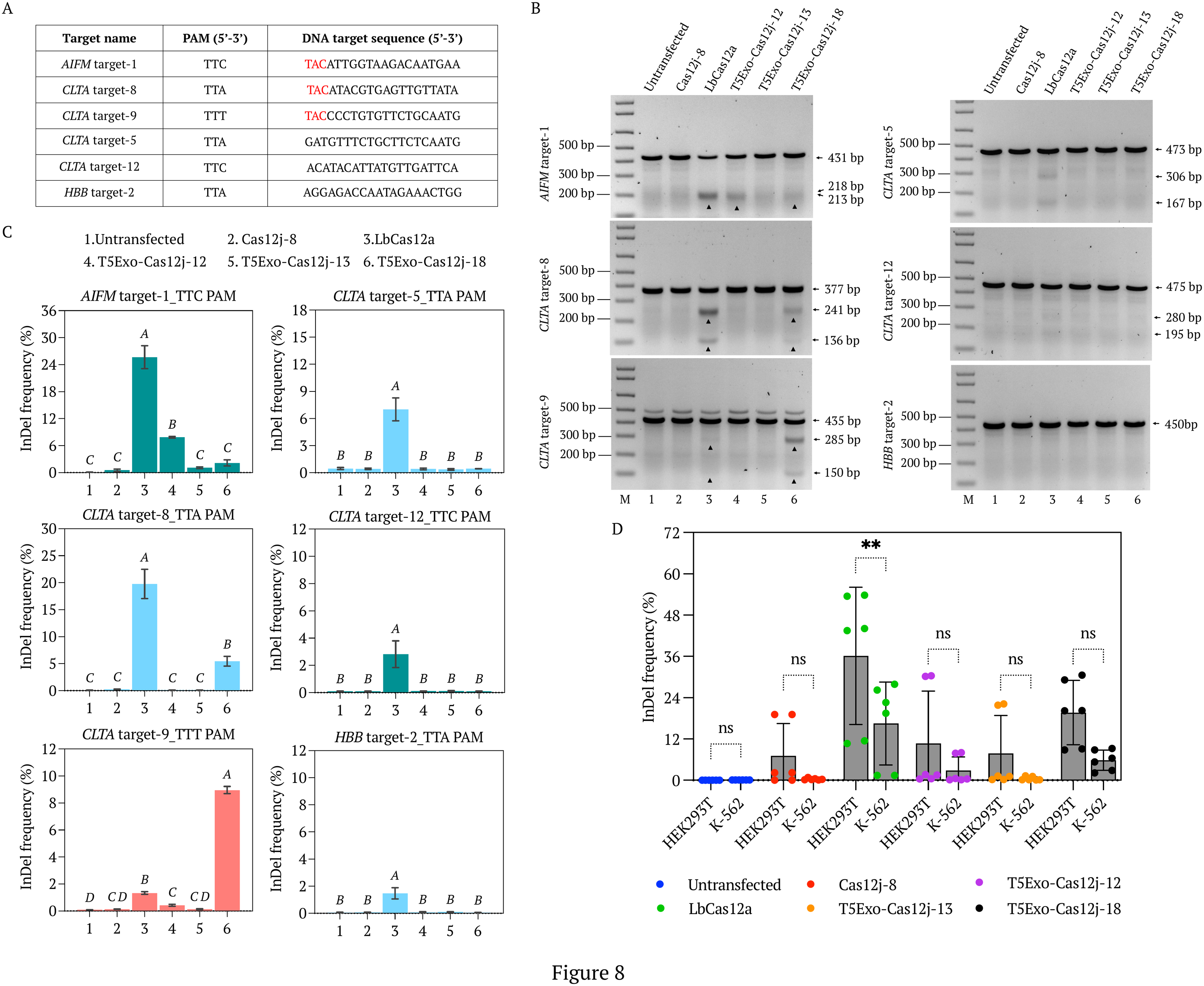

**Supplementary Figure S11.** Gene editing activity of T5Exo-Cas12j fusion proteins in K-562 mammalian cells using crRNAs with and without 5′-end UAC tri-nucleotide sequence. **(A)** Table showing the different DNA target sequences and respective PAM sequences. **(B)** Gels showing the T7EI assay results of the genome editing activity using Cas12j-8 (Lane 2), LbCas12a (Lane 3), and T5Exo-Cas12j-12, -13, and -18 (Lanes 4-6) in K-562 cells at *AIFM* target-1, *HBB* target-2, *CLTA* target-5, -8, -9 and -12 regions, independently. Lane 1 is the T7EI for the untransfected cells used as negative control. Lane M is the 1kb plus marker. The arrow heads indicate the T7EI enzyme cleavage products. **(C)** Deep amplicon sequencing data showing the variable indel efficiencies at *AIFM* target-1, *HBB* target-2, *CLTA* target-5, -8, -9 and -12 regions using different T5Exo-Cas12j variants, Cas12j-8 and LbCas12a. Data represent mean with SD (n = 2 biologically independent replicates). Statistical analysis was performed using one-way ANOVA followed by Tukey’s multiple comparisons test. Bars labeled with different letters are significantly different (*P* < 0.05). Bars sharing the same letter are not significantly different (*P* > 0.05). **(D)** Comparison of T5Exo-Cas12j variants, Cas12j-8 and LbCas12a editing activity on multiple targets (*AIFM* target-1, *CLTA* target-8, and -9) in HEK293T cells and K-562 cells. Data represent mean with SD (n = 6; whole independent data points). *P*-values are calculated using a two-way ANOVA, Bonferroni test (*P*-values indicated are: ns is not significant (0.1234), * (0.0332), ** (0.0021), *** (0.0002), **** (<0.0001)).

**Supplementary files**

**1.** Plasmids used in this study

**(A)** pET28a plasmid backbone used for cloning the *E. coli* expression plasmids*

agcgcctgatgcggtattttctccttacgcatctgtgcggtatttcacaccgCAATGGTGCACTCTCAGTACAATCTGCTCTGATGCCGCATAGTTAAGCCAGTATACACTCCGCTATCGCTACGTGACTGGGTCATGGCTGCGCCCCGACACCCGCCAACACCCGCTGACGCGCCCTGACGGGCTTGTCTGCTCCCGGCATCCGCTTACAGACAAGCTGTGACCGTCTCCGGGAGCTGCATGTGTCAGAGGTTTTCACCGTCATCACCGAAACGCGCGAGGCAGCTGCGGTAAAGCTCATCAGCGTGGTCGTGAAGCGATTCACAGATGTCTGCCTGTTCATCCGCGTCCAGCTCGTTGAGTTTCTCCAGAAGCGTTAATGTCTGGCTTCTGATAAAGCGGGCCATGTTAAGGGCGGTTTTTTCCTGTTTGGTCACTGATGCCTCCGTGTAAGGGGGATTTCTGTTCATGGGGGTAATGATACCGATGAAACGAGAGAGGATGCTCACGATACGGGTTACTGATGATGAACATGCCCGGTTACTGGAACGTTGTGAGGGTAAACAACTGGCGGTATGGATGCGGCGGGACCAGAGAAAAATCACTCAGGGTCAATGCCAGCGCTTCGTTAATACAGATGTAGGTGTTCCACAGGGTAGCCAGCAGCATCCTGCGATGCAGATCCGGAACATAATGGTGCAGGGCGCTGACTTCCGCGTTTCCAGACTTTACGAAACACGGAAACCGAAGACCATTCATGTTGTTGCTCAGGTCGCAGACGTTTTGCAGCAGCAGTCGCTTCACGTTCGCTCGCGTATCGGTGATTCATTCTGCTAACCAGTAAGGCAACCCCGCCAGCCTAGCCGGGTCCTCAACGACAGGAGCACGATCATGCGCACCCGTGGGGCCGCCATGCCGGCGATAATGGCCTGCTTCTCGCCGAAACGTTTGGTGGCGGGACCAGTGACGAAGGCTTGAGCGAGGGCGTGCAAGATTCCGAATACCGCAAGCGACAGGCCGATCATCGTCGCGCTCCAGCGAAAGCGGTCCTCGCCGAAAATGACCCAGAGCGCTGCCGGCACCTGTCCTACGAGTTGCATGATAAAGAAGACAGTCATAAGTGCGGCGACGATAGTCATGCCCCGCGCCCACCGGAAGGAGCTGACTGGGTTGAAGGCTCTCAAGGGCATCGGTCGAGATCCCGGTGCCTAATGAGTGAGCTAACTTACATTAATTGCGTTGCGCTCACTGCCCGCTTTCCAGTCGGGAAACCTGTCGTGCCAGCTGCATTAATGAATCGGCCAACGCGCGGGGAGAGGCGGTTTGCGTATTGGGCGCCAGGGTGGTTTTTCTTTTCACCAGTGAGACGGGCAACAGCTGATTGCCCTTCACCGCCTGGCCCTGAGAGAGTTGCAGCAAGCGGTCCACGCTGGTTTGCCCCAGCAGGCGAAAATCCTGTTTGATGGTGGTTAACGGCGGGATATAACATGAGCTGTCTTCGGTATCGTCGTATCCCACTACCGAGATATCCGCACCAACGCGCAGCCCGGACTCGGTAATGGCGCGCATTGCGCCCAGCGCCATCTGATCGTTGGCAACCAGCATCGCAGTGGGAACGATGCCCTCATTCAGCATTTGCATGGTTTGTTGAAAACCGGACATGGCACTCCAGTCGCCTTCCCGTTCCGCTATCGGCTGAATTTGATTGCGAGTGAGATATTTATGCCAGCCAGCCAGACGCAGACGCGCCGAGACAGAACTTAATGGGCCCTAATACGACTCACTATAGGCCTCTAGAAATAATTTTGTTTAACTTTAAGAAGGAGATATACCATGGGCAGCAGCCATCATCATCATCATCACAGCAGCGGCCTGGTGCCGCGCGGCAGCCATATGGCTAGCATGACTGGTGGACAGCAAATGGGTCGCGGATCCGAATTCGAGCTCCGTCGACAAGCTTGCGGCCGCNNNGATCCGGCTGCTAACAAAGCCCGAAAGGAAGCTGAGTTGGCTGCTGCCACCGCTGAGCAATAACTAGCATAACCCCTTGGGGCCTCTAAACGGGTCTTGAGGGGTTTTTTGCTGAAAGGAGGAACTATATCCGGATTGGCGAATGGGACGCGCCCTGTAGCGGCGCATTAAGCGCGGCGGGTGTGGTGGTTACGCGCAGCGTGACCGCTACACTTGCCAGCGCCCTAGCGCCCGCTCCTTTCGCTTTCTTCCCTTCCTTTCTCGCCACGTTCGCCGGCTTTCCCCGTCAAGCTCTAAATCGGGGGCTCCCTTTAGGGTTCCGATTTAGTGCTTTACGGCACCTCGACCCCAAAAAACTTGATTAGGGTGATGGTTCACGTAGTGGGCCATCGCCCTGATAGACGGTTTTTCGCCCTTTGACGTTGGAGTCCACGTTCTTTAATAGTGGACTCTTGTTCCAAACTGGAACAACACTCAACCCTATCTCGGTCTATTCTTTTGATTTATAAGGGATTTTGCCGATTTCGGCCTATTGGTTAAAAAATGAGCTGATTTAACAAAAATTTAACGCGAATTTTAACAAAATATTAACGCTTACAATTTAGGTGGCACTTTTCGGGGAAATGTGCGCGGAACCCCTATTTGTTTATTTTTCTAAATACATTCAAATATGTATCCGCTCATGAATTAATTCTTAGAAAAACTCATCGAGCATCAAATGAAACTGCAATTTATTCATATCAGGATTATCAATACCATATTTTTGAAAAAGCCGTTTCTGTAATGAAGGAGAAAACTCACCGAGGCAGTTCCATAGGATGGCAAGATCCTGGTATCGGTCTGCGATTCCGACTCGTCCAACATCAATACAACCTATTAATTTCCCCTCGTCAAAAATAAGGTTATCAAGTGAGAAATCACCATGAGTGACGACTGAATCCGGTGAGAATGGCAAAAGTTTATGCATTTCTTTCCAGACTTGTTCAACAGGCCAGCCATTACGCTCGTCATCAAAATCACTCGCATCAACCAAACCGTTATTCATTCGTGATTGCGCCTGAGCGAGACGAAATACGCGATCGCTGTTAAAAGGACAATTACAAACAGGAATCGAATGCAACCGGCGCAGGAACACTGCCAGCGCATCAACAATATTTTCACCTGAATCAGGATATTCTTCTAATACCTGGAATGCTGTTTTCCCGGGGATCGCAGTGGTGAGTAACCATGCATCATCAGGAGTACGGATAAAATGCTTGATGGTCGGAAGAGGCATAAATTCCGTCAGCCAGTTTAGTCTGACCATCTCATCTGTAACATCATTGGCAACGCTACCTTTGCCATGTTTCAGAAACAACTCTGGCGCATCGGGCTTCCCATACAATCGATAGATTGTCGCACCTGATTGCCCGACATTATCGCGAGCCCATTTATACCCATATAAATCAGCATCCATGTTGGAATTTAATCGCGGCCTAGAGCAAGACGTTTCCCGTTGAATATGGCTCATAACACCCCTTGTATTACTGTTTATGTAAGCAGACAGTTTTATTGTTCATGACCAAAATCCCTTAACGTGAGTTTTCGTTCCACTGAGCGTCAGACCCCGTAGAAAAGATCAAAGGATCTTCTTGAGATCCTTTTTTTCTGCGCGTAATCTGCTGCTTGCAAACAAAAAAACCACCGCTACCAGCGGTGGTTTGTTTGCCGGATCAAGAGCCACCAACTCTTTTTCCGAAGGTAACTGGCTTCAGCAGAGCGCAGATACCAAATACTGTCCTTCTAGTGTAGCCGTAGTTAGGCCACCACTTCAAGAACTCTGTAGCACCGCCTACATACCTCGCTCTGCTAATCCTGTTACCAGTGGCTGCTGCCAGTGGCGATAAGTCGTGTCTTACCGGGTTGGACTCAAGACGATAGTTACCGGATAAGGCGCAGCGGTCGGGCTGAACGGGGGGTTCGTGCACACAGCCCAGCTTGGAGCGAACGACCTACACCGAACTGAGATACCTACAGCGTGAGCTATGAGAAAGCGCCACGCTTCCCGAAGGGAGAAAGGCGGACAGGTATCCGGTAAGCGGCAGGGTCGGAACAGGAGAGCGCACGAGGGAGCTTCCAGGGGGAAACGCCTGGTATCTTTATAGTCCTGTCGGGTTTCGCCACCTCTGACTTGAGCGTCGATTTTTGTGATGCTCGTCAGGGGGGCGGAGCCTATGGAAAAACGCCAGCAACGCGGCCTTTTTACGGTTCCTGGCCTTTTGCTGGCCTTTTGCTCACATGTTCTTTCCTGCGTTATCCCCTGATTCTGTGGATAACCGTATTACCGCCTTTGAGTGAGCTGATACCGCTCGCCGCAGCCGAACGACCGAGCGCAGCGAGTCAGTGAGCGAGGAAGCGGAAG

* *E. coli* optimized codon sequences of Cas12j variants are replaced with NNN sequence in the plasmid and cloned the bacterial expression plasmids

* *E. coli* optimized Cas12j-8 sequence (Wang et al., (2023)) synthesized and cloned between BamHI and HindIII sites (Pink and green highlights are BamHI and HindIII respectively)

**(B)** Plasmid backbone used for cloning different protein coding sequences (Cas12j orthologues, T5Exo-Cas12j variants and Be-(d)Cas12j base editors) for expression in mammalian cells (Addgene - 194965)*

cctgcaggcagctgcgcgctcgctcgctcactgaggccgcccgggcgtcgggcgacctttggtcgcccggcctcagtgagcgagcgagcgcgcagagagggagtggccaactccatcactaggggttcctgcggcctctagactcgaggcgttgacattgattattgactagttattaatagtaatcaattacggggtcattagttcatagcccatatatggagttccgcgttacataacttacggtaaatggcccgcctggctgaccgcccaacgacccccgcccattgacgtcaataatgacgtatgttcccatagtaacgccaatagggactttccattgacgtcaatgggtggagtatttacggtaaactgcccacttggcagtacatcaagtgtatcatatgccaagtacgccccctattgacgtcaatgacggtaaatggcccgcctggcattatgcccagtacatgaccttatgggactttcctacttggcagtacatctacgtattagtcatcgctattaccatggtgatgcggttttggcagtacatcaatgggcgtggatagcggtttgactcacggggatttccaagtctccaccccattgacgtcaatgggagtttgttttggcaccaaaatcaacgggactttccaaaatgtcgtaacaactccgccccattgacgcaaatgggcggtaggcgtgtacggtgggaggtctatataagcagagctctctggctaactaccggtgccaccatggccccaaagaagaagcggaaggtcNNNaaaaggccggcggccacgaaaaaggccggccaggcaaaaaagaaaaagggatccgctagcggcagcggcgccaccaacttcagcctgctgaagcaggccggcgacgtggaggagaaccccggccccatgaccgagtacaagcccacggtgcgcctcgccacccgcgacgacgtccccagggccgtacgcaccctcgccgccgcgttcgccgactaccccgccacgcgccacaccgtcgatccggaccgccacatcgagcgggtcaccgagctgcaagaactcttcctcacgcgcgtcgggctcgacatcggcaaggtgtgggtcgcggacgacggcgccgcggtggcggtctggaccacgccggagagcgtcgaagcgggggcggtgttcgccgagatcggcccgcgcatggccgagttgagcggttcccggctggccgcgcagcaacagatggaaggcctcctggcgccgcaccggcccaaggagcccgcgtggttcctggccaccgtcggagtctcgcccgaccaccagggcaagggtctgggcagcgccgtcgtgctccccggagtggaggcggccgagcgcgccggggtgcccgccttcctggaaacctccgcgccccgcaacctccccttctacgagcggctcggcttcaccgtcaccgccgacgtcgaggtgcccgaaggaccgcgcacctggtgcatgacccgcaagcccggtgcctacccatacgatgttccagattacgcttacccatacgatgttccagattacgcttacccatacgatgttccagattacgcttaagaattcctagagctcgctgatcagcctcgactgtgccttctagttgccagccatctgttgtttgcccctcccccgtgccttccttgaccctggaaggtgccactcccactgtcctttcctaataaaatgaggaaattgcatcgcattgtctgagtaggtgtcattctattctggggggtggggtggggcaggacagcaagggggaggattgggaagagaatagcaggcatgctggggaggtaccgagggcctatttcccatgattccttcatatttgcatatacgatacaaggctgttagagagataattggaattaatttgactgtaaacacaaagatattagtacaaaatacgtgacgtagaaagtaataatttcttgggtagtttgcagttttaaaattatgttttaaaatggactatcatatgcttaccgtaacttgaaagtatttcgatttcttggctttatatatcttgtggaaaggacgtcttctcagaagacgttttttttgcggccgcaggaacccctagtgatggagttggccactccctctctgcgcgctcgctcgctcactgaggccgggcgaccaaaggtcgcccgacgcccgggctttgcccgggcggcctcagtgagcgagcgagcgcgcagctgcctgcaggggcgcctgatgcggtattttctccttacgcatctgtgcggtatttcacaccgcatacgtcaaagcaaccatagtacgcgccctgtagcggcgcattaagcgcggcgggtgtggtggttacgcgcagcgtgaccgctacacttgccagcgccttagcgcccgctcctttcgctttcttcccttcctttctcgccacgttcgccggctttccccgtcaagctctaaatcgggggctccctttagggttccgatttagtgctttacggcacctcgaccccaaaaaacttgatttgggtgatggttcacgtagtgggccatcgccctgatagacggtttttcgccctttgacgttggagtccacgttctttaatagtggactcttgttccaaactggaacaacactcaactctatctcgggctattcttttgatttataagggattttgccgatttcggtctattggttaaaaaatgagctgatttaacaaaaatttaacgcgaattttaacaaaatattaacgtttacaattttatggtgcactctcagtacaatctgctctgatgccgcatagttaagccagccccgacacccgccaacacccgctgacgcgccctgacgggcttgtctgctcccggcatccgcttacagacaagctgtgaccgtctccgggagctgcatgtgtcagaggttttcaccgtcatcaccgaaacgcgcgagacgaaagggcctcgtgatacgcctatttttataggttaatgtcatgataataatggtttcttagacgtcaggtggcacttttcggggaaatgtgcgcggaacccctatttgtttatttttctaaatacattcaaatatgtatccgctcatgagacaataaccctgataaatgcttcaataatattgaaaaaggaagagtatgagtattcaacatttccgtgtcgcccttattcccttttttgcggcattttgccttcctgtttttgctcacccagaaacgctggtgaaagtaaaagatgctgaagatcagttgggtgcacgagtgggttacatcgaactggatctcaacagcggtaagatccttgagagttttcgccccgaagaacgttttccaatgatgagcacttttaaagttctgctatgtggcgcggtattatcccgtattgacgccgggcaagagcaactcggtcgccgcatacactattctcagaatgacttggttgagtactcaccagtcacagaaaagcatcttacggatggcatgacagtaagagaattatgcagtgctgccataaccatgagtgataacactgcggccaacttacttctgacaacgatcggaggaccgaaggagctaaccgcttttttgcacaacatgggggatcatgtaactcgccttgatcgttgggaaccggagctgaatgaagccataccaaacgacgagcgtgacaccacgatgcctgtagcaatggcaacaacgttgcgcaaactattaactggcgaactacttactctagcttcccggcaacaattaatagactggatggaggcggataaagttgcaggaccacttctgcgctcggcccttccggctggctggtttattgctgataaatctggagccggtgagcgtggaagccgcggtatcattgcagcactggggccagatggtaagccctcccgtatcgtagttatctacacgacggggagtcaggcaactatggatgaacgaaatagacagatcgctgagataggtgcctcactgattaagcattggtaactgtcagaccaagtttactcatatatactttagattgatttaaaacttcatttttaatttaaaaggatctaggtgaagatcctttttgataatctcatgaccaaaatcccttaacgtgagttttcgttccactgagcgtcagaccccgtagaaaagatcaaaggatcttcttgagatcctttttttctgcgcgtaatctgctgcttgcaaacaaaaaaaccaccgctaccagcggtggtttgtttgccggatcaagagctaccaactctttttccgaaggtaactggcttcagcagagcgcagataccaaatactgttcttctagtgtagccgtagttaggccaccacttcaagaactctgtagcaccgcctacatacctcgctctgctaatcctgttaccagtggctgctgccagtggcgataagtcgtgtcttaccgggttggactcaagacgatagttaccggataaggcgcagcggtcgggctgaacggggggttcgtgcacacagcccagcttggagcgaacgacctacaccgaactgagatacctacagcgtgagctatgagaaagcgccacgcttcccgaagggagaaaggcggacaggtatccggtaagcggcagggtcggaacaggagagcgcacgagggagcttccagggggaaacgcctggtatctttatagtcctgtcgggtttcgccacctctgacttgagcgtcgatttttgtgatgctcgtcaggggggcggagcctatggaaaaacgccagcaacgcggcctttttacggttcctggccttttgctggccttttgctcacatgt

* Human optimized codon sequences of Cas12j variants, T5Exo-Cas12j variants, Be-(d)Cas12j variants are cloned at NNN sequence

**(C)** Modified pJET 1.2 plasmid sequence used as *in vitro* target (4041bp)

GCCCCTGCAGCCGAATTATATTATTTTTGCCAAATAATTTTTAACAAAAGCTCTGAAGTCTTCTTCATTTAAATTCTTAGATGATACTTCATCTGGAAAATTGTCCCAATTAGTAGCATCACGCTGTGAGTAAGTTCTAAACCATTTTTTTATTGTTGTATTATCTCTAATCTTACTACTCGATGAGTTTTCGGTATTATCTCTATTTTTAACTTGGAGCAGGTTCCATTCATTGTTTTTTTCATCATAGTGAATAAAATCAACTGCTTTAACACTTGTGCCTGAACACCATATCCATCCGGCGTAATACGACTCACTATAGGGAGAGCGGCCGCCAGATCTTCCGGATGGCTCGAGTTTTTCAGCAACACATGGGTGTGTGCAAACCGTTTTGGGTTACACATTTACAAGCAACTTATATAATAATACTAAACTACAATAATTCATGTATAAAACTAAGGGCGTAACCGAAATCGGTTGAACCGAAACCGGTTAGTATAAAAGCAGACATTTTATGCACCAAAAGAGAACTGCAATGTTTCAGGACCCACAGGAGCGACCCAGAAAGTTACCACAGTTATGCACAGAGCTGCAAACAACTATACATGATATAATATTAGAATGTGTGTACTGCAAGCAACAGTTACTGCGACGTGAGGTATATGACTTTGCTTTTCGGGATTTATGCATAGTATATAGAGATGGGAATCCATATGCTGTATGTGATAAATGTTTAAAGTTTTATTCTAAAATTAGTGAGTATAGACATTATTGTTATAGTTTGTATGGAACAACATTAGAACAGCAATACAACAAACCGTTGTGTGATTTGTTAATTAGGTGTATTAACTGTCAAAAGCCACTGTGTCCTGAAGAAAAGCAAAGACATCTGGACAAAAAGCAAAGATTCCATAATATAAGGGGTCGGTGGACCGGTCGATGTATGTCTTGTTGCAGATCATCAAGAACACGTAGAGAAACCCAGCTGTAATCATGCATGGAGATACACCTACATTGCATGAATATATGTTAGATTTGCAACCAGAGACAACTGATCTCTACTGTTATGAGCAATTAAATGACAGCTCAGAGGAGGAGGATGAAATAGATGGTCCAGCTGGACAAGCAGAACCGGACAGAGCCCATTACAATATTGTAACCTTTTGTTGCAAGTGTGACTCTACGCTTCGGTTGTGCGTACAAAGCACACACGTAGACATTCGTACTTTGGAAGACCTGTTAATGGGCACACTAGGAATTGTGTGCCCCATCTGTTCTCAGAAACCATAATCTACCATGGCTGATCCTGCAGGTACCAATGGGGAAGAGGGTACGGGATGTAATGGATGGTTTTATGTAGAGGCTGTAGTGGAAAAAAAAACAGGGGATGCTATATCAGATGACGAGAACGAAAATGACAGTGATACAGGTGAAGGATATCTTTCTAGAAGATCTCCTACAATATTCTCAGCTGCCATGGAAAATCGATGTTCTTCTTTTATTCTCTCAAGATTTTCAGGCTGTATATTAAAACTTATATTAAGAACTATGCTAACCACCTCATCAGGAACCGTTGTAGGTGGCGTGGGTTTTCTTGGCAATCGACTCTCATGAAAACTACGAGCTAAATATTCAATATGTTCCTCTTGACCAACTTTATTCTGCATTTTTTTTGAACGAGGTTTAGAGCAAGCTTCAGGAAACTGAGACAGGAATTTTATTAAAAATTTAAATTTTGAAGAAAGTTCAGGGTTAATAGCATCCATTTTTTGCTTTGCAAGTTCCTCAGCATTCTTAACAAAAGACGTCTCTTTTGACATGTTTAAAGTTTAAACCTCCTGTGTGAAATTATTATCCGCTCATAATTCCACACATTATACGAGCCGGAAGCATAAAGTGTAAAGCCTGGGGTGCCTAATGAGTGAGCTAACTCACATTAATTGCGTTGCGCTCACTGCCAATTGCTTTCCAGTCGGGAAACCTGTCGTGCCAGCTGCATTAATGAATCGGCCAACGCGCGGGGAGAGGCGGTTTGCGTATTGGGCGCTCTTCCGCTTCCTCGCTCACTGACTCGCTGCGCTCGGTCGTTCGGCTGCGGCGAGCGGTATCAGCTCACTCAAAGGCGGTAATACGGTTATCCACAGAATCAGGGGATAACGCAGGAAAGAACATGTGAGCAAAAGGCCAGCAAAAGGCCAGGAACCGTAAAAAGGCCGCGTTGCTGGCGTTTTTCCATAGGCTCCGCCCCCCTGACGAGCATCACAAAAATCGACGCTCAAGTCAGAGGTGGCGAAACCCGACAGGACTATAAAGATACCAGGCGTTTCCCCCTGGAAGCTCCCTCGTGCGCTCTCCTGTTCCGACCCTGCCGCTTACCGGATACCTGTCCGCCTTTCTCCCTTCGGGAAGCGTGGCGCTTTCTCATAGCTCACGCTGTAGGTATCTCAGTTCGGTGTAGGTCGTTCGCTCCAAGCTGGGCTGTGTGCACGAACCCCCCGTTCAGCCCGACCGCTGCGCCTTATCCGGTAACTATCGTCTTGAGTCCAACCCGGTAAGACACGACTTATCGCCACTGGCAGCAGCCACTGGTAACAGGATTAGCAGAGCGAGGTATGTAGGCGGTGCTACAGAGTTCTTGAAGTGGTGGCCTAACTACGGCTACACTAGAAGGACAGTATTTGGTATCTGCGCTCTGCTGAAGCCAGTTACCTTCGGAAAAAGAGTTGGTAGCTCTTGATCCGGCAAACAAACCACCGCTGGTAGCGGTGGTTTTTTTGTTTGCAAGCAGCAGATTACGCGCAGAAAAAAAGGATCTCAAGAAGATCCTTTGATCTTTTCTACGGGGTCTGACGCTCAGTGGAACGAAAACTCACGTTAAGGGATTTTGGTCATGAGATTATCAAAAAGGATCTTCACCTAGATCCTTTTAAATTAAAAATGAAGTTTTAAATCAATCTAAAGTATATATGAGTAAACTTGGTCTGACAGTTACCAATGCTTAATCAGTGAGGCACCTATCTCAGCGATCTGTCTATTTCGTTCATCCATAGTTGCCTGACTCCCCGTCGTGTAGATAACTACGATACGGGAGGGCTTACCATCTGGCCCCAGTGCTGCAATGATACCGCGAGACCCACGCTCACCGGCTCCAGATTTATCAGCAATAAACCAGCCAGCCGGAAGGGCCGAGCGCAGAAGTGGTCCTGCAACTTTATCCGCCTCCATCCAGTCTATTAATTGTTGCCGGGAAGCTAGAGTAAGTAGTTCGCCAGTTAATAGTTTGCGCAACGTTGTTGCCATTGCTACAGGCATCGTGGTGTCACGCTCGTCGTTTGGTATGGCTTCATTCAGCTCCGGTTCCCAACGATCAAGGCGAGTTACATGATCCCCCATGTTGTGCAAAAAAGCGGTTAGCTCCTTCGGTCCTCCGATCGTTGTCAGAAGTAAGTTGGCCGCAGTGTTATCACTCATGGTTATGGCAGCACTGCATAATTCTCTTACTGTCATGCCATCCGTAAGATGCTTTTCTGTGACTGGTGAGTACTCAACCAAGTCATTCTGAGAATAGTGTATGCGGCGACCGAGTTGCTCTTGCCCGGCGTCAATACGGGATAATACCGCGCCACATAGCAGAACTTTAAAAGTGCTCATCATTGGAAAACGTTCTTCGGGGCGAAAACTCTCAAGGATCTTACCGCTGTTGAGATCCAGTTCGATGTAACCCACTCGTGCACCCAACTGATCTTCAGCATCTTTTACTTTCACCAGCGTTTCTGGGTGAGCAAAAACAGGAAGGCAAAATGCCGCAAAAAAGGGAATAAGGGCGACACGGAAATGTTGAATACTCATACTCTTCCTTTTTCAATATTATTGAAGCATTTATCAGGGTTATTGTCTCATGAGCGGATACATATTTGAATGTATTTAGAAAAATAAACAAATAGGGGTTCCGCGCACATTTCCCCGAAAAGTGCCACCTGACGTCTAAGAAACCATTATTATCATGACATTAACCTATAAAAATAGGCGTATCACGAGGCC

Green highlighted: Spacer sequence for *in vitro* targeting

**(D)** Addgene plasmid (Addgene - 204639) backbone modified and used for crRNA cloning and expression*

attgttgccgggaagctagagtaagtagttcgccagttaatagtttgcgcaacgttgttgccattgctacaggcatcgtggtgtcacgctcgtcgtttggtatggcttcattcagctccggttcccaacgatcaaggcgagttacatgatcccccatgttgtgcaaaaaagcggttagctccttcggtcctccgatcgttgtcagaagtaagttggccgcagtgttatcactcatggttatggcagcactgcataattctcttactgtcatgccatccgtaagatgcttttctgtgactggtgagtactcaaccaagtcattctgagaatagtgtatgcggcgaccgagttgctcttgcccggcgtcaatacgggataataccgcgccacatagcagaactttaaaagtgctcatcattggaaaacgttcttcggggcgaaaactctcaaggatcttaccgctgttgagatccagttcgatgtaacccactcgtgcacccaactgatcttcagcatcttttactttcaccagcgtttctgggtgagcaaaaacaggaaggcaaaatgccgcaaaaaagggaataagggcgacacggaaatgttgaatactcatactcttcctttttcaatattattgaagcatttatcagggttattgtctcatgagcggatacatatttgaatgtatttagaaaaataaacaaataggggttccgcgcacatttccccgaaaagtgccacctgacgtcgctagctgtacaaaaaagcaggctttaaaggaaccaattcagtcgactggatccggtaccaaggtcgggcaggaagagggcctatttcccatgattccttcatatttgcatatacgatacaaggctgttagagagataattggaattaatttgactgtaaacacaaagatattagtacaaaatacgtgacgtagaaagtaataatttcttgggtagtttgcagttttaaaattatgttttaaaatggactatcatatgcttaccgtaacttgaaagtatttcgatttcttggctttatatatcttgtggaaaggacgaaacaccggtgtcaacgccagcgcggaggcgtcaaatccgcGACtggtcttcatctacggagaagacatggccggcatggtcccagcctcctcgctggcgccggctgggcaacatgcttcggcatggcgaatgggacttttttttaagcttgggccgctcgaggtacctctctacatatgacatgtgagcaaaaggccagcaaaaggccaggaaccgtaaaaaggccgcgttgctggcgtttttccataggctccgcccccctgacgagcatcacaaaaatcgacgctcaagtcagaggtggcgaaacccgacaggactataaagataccaggcgtttccccctggaagctccctcgtgcgctctcctgttccgaccctgccgcttaccggatacctgtccgcctttctcccttcgggaagcgtggcgctttctcatagctcacgctgtaggtatctcagttcggtgtaggtcgttcgctccaagctgggctgtgtgcacgaaccccccgttcagcccgaccgctgcgccttatccggtaactatcgtcttgagtccaacccggtaagacacgacttatcgccactggcagcagccactggtaacaggattagcagagcgaggtatgtaggcggtgctacagagttcttgaagtggtggcctaactacggctacactagaagaacagtatttggtatctgcgctctgctgaagccagttaccttcggaaaaagagttggtagctcttgatccggcaaacaaaccaccgctggtagcggtggtttttttgtttgcaagcagcagattacgcgcagaaaaaaaggatctcaagaagatcctttgatcttttctacggggtctgacgctcagtggaacgaaaactcacgttaagggattttggtcatgagattatcaaaaaggatcttcacctagatccttttaaattaaaaatgaagttttaaatcaatctaaagtatatatgagtaaacttggtctgacagttaccaatgcttaatcagtgaggcacctatctcagcgatctgtctatttcgttcatccatagttgcctgactccccgtcgtgtagataactacgatacgggagggcttaccatctggccccagtgctgcaatgataccgcgagacccacgctcaccggctccagatttatcagcaataaaccagccagccggaagggccgagcgcagaagtggtcctgcaactttatccgcctccatccagtctatta

Red highlighted: AgeI

Pink highlighted: BbsI

* All the crRNA encoding DNA sequences were cloned at AgeI and BbsI restriction sites and for human cell expression

Green highlighted: U6 promoter

Yellow highlighted: HDV ribozyme

Grey highlighted: Forward and reverse primers used for the amplification of U6-crRNA sequences

**Supplementary Tables**

**Supplementary Table S1.** Protein sequences of different Cas12j orthologues

| **Protein** | **Sequence** | **Contig source** |
| --- | --- | --- |
| Cas12j-11 | MPGPTLSDVAPTNCECGAAAPSATLKLSDVIKTHFPAGRFRKDHQKTAGKKLKHEGEEACVEYLRNKVSDYPPNFKPPAKGTIVAQSRPFSEWPIVRASEAIQKYVYGLTVAELDVFSPGTSKPSHAEWFAKTGVENYGYRQVQGLNTIFQNTVNRFKGVLKKVENRNKKSLKRQEGANRRRVEEGLPEVPVTVESATDDEGRLLQPPGVNPSIYGYQGVAPRVCTDLQGFSGMSVDFAGYRRDPDAVLVESLPEGRLSIPKGERGYVPEWQRDPERNKFPLREGSRRQRKWYSNACHKPKPGRTSKYDPEALKKASAKDALLVSISIGEDWAIIDVRGLLRDARRRGFTPEEGLSLNSLLGLFTEYPVFDVQRGLITFTYKLGQVDVHSRKTVPTFRSRALLESLVAKEEIALVSVDLGQTNPASMKVSRVRAQEGALVAEPVHRMFLSDVLLGELSSYRKRMDAFEDAIRAQAFETMTPEQQAEITRVCDVSVEVARRRVCEKYSISPQDVPWGEMTGHSTFIVDAVLRKGGDESLVYFKNKEGETLKFRDLRISRMEGVRPRLTKDTRDALNKAVLDLKRAHPTFAKLAKQKLELARRCVNFIEREAKRYTQCERVVFVIEDLNVGFFHGKGKRDRGWDAFFTAKKENRWVIQALHKAFSDLGLHRGSYVIEVTPQRTSMTCPRCGHCDKGNRNGEKFVCLQCGATLHADLEVATDNIERVALTGKAMPKPPVRERSGDVQKAGTARKARKPLKPKQKTEPSVQEGSSDDGVDKSPGDASRNPVYNPSDTLSI | >IMGVR_UViG_3300035182_002586 |
| Cas12j-12 | MEKAGPTSPLSVLIHKNFEGCRFQIDHLKIAGRKLAREGEAAAIEYLLDKKCEGLPPNFQPPAKGNVIAQSRPFTEWAPYRASVAIQKYIYSLSVDERKVCDPGSSSDSHEKWFKQTGVQNYGYTHVQGLNLIFKHALARYDGVLKKVDNRNEKNRKKAERVNSFRREEGLPEEVFEEEKATDETGHLLQPPGVNHSIYCYQSVRPKPFNPRKPGGISLPEAYSGYSLKPQDELPIGSLDRLSIPPGQPGYVPEWQRSQLTTQKHRRKRSWYSAQKWKPRTGRTSTFDPDRLNCARAQGAILAVVRIHEDWVVFDVRGLLRNALWRELAGKGLTVRDLLDFFTGDPVVDTKRGVVTFTYKLGKVDVHSLRTVRGKRSKKVLEDLTLSSDVGLVTIDLGQTNVLAADYSKVTRSENGELLAVPLSKSFLPKHLLHEVTAYRTSYDQMEEGFRRKALLTLTEDQQVEVTLVRDFSVESSKTKLLQLGVDVTSLPWEKMSSNTTYISDQLLQQGADPASLFFDGERDGKPCRHKKKDRTWAYLVRPKVSPETRKALNEALWALKNTSPEFESLSKRKIQFSRRCMNYLLNEAKRISGCGQVVFVIEDLNVRVHHGRGKRAIGWDNFFKPKRENRWFMQALHKAASELAIHRGMHIIEACPARSSITCPKCGHCDPENRCSSDREKFLCVKCGAAFHADLEVATFNLRKVALTGTALPKSIDHSRDGLIPKGARNRKLKEPQANDEKACA | >IMGVR_UViG_3300037445_000082 |
| Cas12j-13 | MKKPNNIRRIREEHFEGLCFGKDVLTKAGKIYEKDGEEAAIDFLMGKDEEDPPNFKPPAKTTIVAQSRPFDQWPIYQVSQAVQERVFAYTEEEFNASKEALFSGDISSKSRDFWFKTNNISDQGIGAQGLNTILSHAFSRYSGVIKKVENRNKKRLKKLSKKNQLKIEEGLEILEFKPDSAFNENGLLAQPPGINPNIYGYQAVTPFVFDPDNPGDVILPKQYEGYSRKPDDIIEKGPSRLDIPKGQPGYVPEHQRKNLKKKGRVRLYRRTPPKTKALASILAVLQIGKDWVLFDMRGLLRSVYMREAATPGQISAKDLLDTFTGCPVLNTRTGEFTFCYKLRSEGALHARKIYTKGETRTLLTSLTSENNTIALVTVDLGQRNPAAIMISRLSRKEELSEKDIQPVSRRLLPDRYLNELKRYRDAYDAFRQEVRDEAFTSLCPEHQEQVQQYEALTPEKAKNLVLKHFFGTHDPDLPWDDMTSNTHYIANLYLERGGDPSKVFFTRPLKKDSKSKKPRKPTKRTDASISRLPEIRPKMPEDARKAFEKAKWEIYTGHEKFPKLAKRVNQLCREIANWIEKEAKRLTLCDTVVVGIEDLSLPPKRGKGKFQETWQGFFRQKFENRWVIDTLKKAIQNRAHDKGKYVLGLAPYWTSQRCPACGFIHKSNRNGDHFKCLKCEALFHADSEVATWNLALVAVLGKGITNPDSKKPSGQKKTGTTRKKQIKGKNKGKETVNVPPTTQEVEDIIAFFEKDDETVRNPVYKPTGT | >IMGVR_UViG_3300036713_000599 |
| Cas12j-14 | MPDKKETPLVALCKKSFPGLRFKKHDSRQAGRILKSKGEGAAVAFLEGKGGTTQPNFKPPVKCNIVAMSRPLEEWPIYKASVVIQKYVYAQSYEEFKATDPGKSEAGLRAWLKATRVDTDGYFNVQGLNLIFQNARATYEGVLKKVENRNSKKVAKIEQRNEHRAERGLPLLTLDEPETALDETGHLRHRPGINCSVFGYQHMKLKPYVPGSIPGVTGYSRDPSTPIAACGVDRLEIPEGQPGYVPPWDRENLSVKKHRRKRASWARSRGGAIDDNMLLAVVRVADDWALLDLRGLLRNTQYRKLLDRSVPVTIESLLNLVTNDPTLSVVKKPGKPVRYTATLIYKQGVVPVVKAKVVKGSYVSKMLDDTTETFSLVGVDLGVNNLIAANALRIRPGKCVERLQAFTLPEQTVEDFFRFRKAYDKHQENLRLAAVRSLTAEQQAEVLALDTFGPEQAKMQVCGHLGLSVDEVPWDKVNSRSSILSDLAKERGVDDTLYMFPFFKGKGKKRKTEIRKRWDVNWAQHFRPQLTSETRKALNEAKWEAERNSSKYHQLSIRKKELSRHCVNYVIRTAEKRAQCGKVIVAVEDLHHSFRRGGKGSRKSGWGGFFAAKQEGRWLMDALFGAFCDLAVHRGYRVIKVDPYNTSRTCPECGHCDKANRDRVNREAFICVCCGYRGNADIDVAAYNIAMVAITGVSLRKAARASVASTPLESLAAE | >IMGVR_UViG_3300037722_000745 |
| Cas12j-15 | MTPSPQIARLVETPLAAALKAHHPGKKFRSDYLKKAGKILKDQGVEAAMAHLDGKDQAEPPNFKPPAKCRIVARSREFSEWPIVKASVEIQKYIYGLTLEERKACDPGKSSASHKAWFAKTGVNTFGYSSVQGFNLIFGHTLGRYDGVLVKTENLNKKRAEKNERFRAKALAEGRAEPVCPPLVTATNDTGQDVTLEDGRVVRPGQLLQPPGINPNIYAYQQVSPKAYVPGIIELPEEFQGYSRDPNAVILPLVPRDRLSIPKGQPGYVPEPHREGLTGRKDRRMRRYYETERGTKLKRPPLTAKGRADKANEALLVVVRIDSDWVVMDVRGLLRNARWRRLVSKEGITLNGLLDLFTGDPVLNPKDCSVSRDTGDPVNDPRHGVVTFCYKLGVVDVCSKDRPIKGFRTKEVLERLTSSGTVGMVSIDLGQTNPVAAAVSRVTKGLQAETLETFTLPDDLLGKVRAYRAKTDRMEEGFRRNALRKLTAEQQAEITRYNDATEQQAKALVCSTYGIGPEEVPWERMTSNTTYISDHILDHGGDPDTVFFMATKRGQNKPTLHKRKDKAWGQKFRPAISVETRLARQAAEWELRRASLEFQKLSVWKTELCRQAVNYVMERTKKRTQCDVIIPVIEDLPVPLFHGSGKRDPGWANFFVHKRENRWFIDGLHKAFSELGKHRGIYVFEVCPQRTSITCPKCGHCDPDNRDGEKFVCLSCQATLHADLDVATTNLVRVALTGKVMPRSERSGDAQTPGPARKARTGKIKGSKPTSAPQGATQTDAKAHLSQTGV | >IMGVR_UViG_3300031746_004289 |
| Cas12j-16 | MGCYNAGAMKKTTNLKIENGVSPLAQMTRKHFPGKRFPASVLKPAGRKLKDHGEQAAIEFLQANIDVPYGNFKAPAKCNVVATSRPYSEWPLYKFSSELQKAVFALSKDRLMEIEPSKQSDAENEKFLSAIGVSADPSINVTFVSACISKAVHTYLGMEKKAENKYQKKLSRCRSESELSSVTPENVYNEDGTLSDGWRPGFNANLYGNSNSKLSLFGSVRSHNKVELPVWLSEYREWAKNRSKDSKINEYSASVDRLSIPEGQPGHVPLWQRDSSKRTKPGGGEIKEGVRRHRWYSNRNNANRRNKVDQATRLAASAMEMVLAIAFFGEDWVLFDIRGLLRNARYRKLVNKNTTYGDLMELFTADPVLDTKRGIITASYKDTTLKIVQQTIVGEKKSKSKILEEVQKNGPVAVVSIDLGVNEPVSYRVSRVDVAGGNAIVAELAAEGFMSNELKKEISSYREKSDELNGDLREKAVLSLSDEMQAEIRRVDATNASDSKNRICEMLSLDPESVDWSKMTTQTRFIFNKHVENGGDPNVLLFTPTEDKKNKGKKSKNKKGEYGDRVPHSDSGIARNIAREKLSMETCEALNKAKRELQQEDPRYGKLSKRKQEFARRVVNGVVVRAQEVTGCDNVVLVVEKLNVSNKMFSGSGKRAPGWDNFFVHKKENRWFIQALHKAFTDKAAHKGIPVIEIKPSYTSQTCPACEHCDKDNRDGVHFCCTRCGFTGHADLNVACFNIEKVALTGEAMSGPGSATAHKKTRKPKKAMVESDKAA | >IMGVR_UViG_3300048834_000512 |
| Cas12j-17 | MSKEKTPPSAYAILKAKHFPDLDFEKKHKMMAGRMFKNGASEQEVVQYLQGKGSESLMDVKPPAKSPILAQSRPFDEWEMVRTSRLIQETIFGIPKRGSIPKRDGLSETQFNELVASLEVGGKPMLNKQTRAIFYGLLGIKPPTFHAMAQNILIDLAINIRKGVLKKVDNLNEKNRKKVKRIRDAGEQDVMVPAEVTAHDDRGYLNHPPGVNPTIPGYQGVVIPFPEGFEGLPSGMTPVDWSHVLVDYLPHDRLSIPKGSPGYIPEWQRPLLNRHKGRRHRSWYANSLNKPRKSRTEEAKDRQNAGKRTALIEAERLKGVLPVLMRFKEDWLIIDARGLLRNARYRGVLPEGSTLGNLIDLFSDSPRVDTRRGICTFLYRKGRAYSTKPVKRKESKETLLKLTEKSTIALVSIDLGQTNPLTAKLSKVRQVDGCLVAEPVLRKLIDNASEDGKEIARYRVAHDLLRARILEDAIDLLGIYKDEVVRARSDTPDLCKERVCRFLGLDSQAIDWDRMTPYTDFIAQAFVAKGGDPKVVTIKPNGKPKMFRKDRSIKNMKGIRLDISKEASSAYREAQWAIQRESPDFQRLAVWQSQLTKRIVNQLVAWAKKCTQCDTVVLAFEDLNIGMMHGSGKWANGGWNALFLHKQENRWFMQAFHKALTELSAHKGIPTIEVLPHRTSITCTQCGHCHPGNRDGERFKCLKCEFLANTDLEIATDNIERVALTGLPMPKGERSSAKRKPGGTRKTKKSKHSGNSPLAAE | >IMGVR_UViG_3300037471_001492 |
| Cas12j-18 | MLPPSNKIGKSMSLKEFINKRNFKSSIIKQAGKILKKEGEEAVKKYLDDNYVEGYKKRDFPITAKCNIVASNRKIEDFDISKFSSFIQNYVFNLNKDNFEEFSKIKYNRKSFDELYKKIANEIGLEKPNYENIQGEIAVIRNAINIYNGVLKKVENRNKKIQEKNQSKDPPKLLSAFDDNGFLAERPGINETIYGYQSVRLRHLDVEKDKDIIVQLPDIYQKYNKKSTDKISVKKRLNKYNVDEYGKLISKRRKERINKDDAILCVSNFGDDWIIFDARGLLRQTYRYKLKKKGLCIKDLLNLFTGDPIINPTKTDLKEALSLSFKDGIINNRTLKVKNYKKCPELISELIRDKGKVAMISIDLGQTNPISYRLSKFTANNVAYIENGVISEDDIVKMKKWREKSDKLENLIKEEAIASLSDDEQREVRLYENDIADNTKKKILEKFNIREEDLDFSKMSNNTYFIRDCLKNKNIDESEFTFEKNGKKLDPTDACFAREYKNKLSELTRKKINEKIWEIKKNSKEYHKISIYKKETIRYIVNKLIKQSKEKSECDDIIVNIEKLQIGGNFFGGRGKRDPGWNNFFLPKEENRWFINACHKAFSELAPHKGIIVIESDPAYTSQTCPKCENCDKENRNGEKFKCKKCNYEANADIDVATENLEKIAKNGRRLIKNFDQLGERLPGAEMPGGARKRKPSKSLPKNGRGAGVGSEPELINQSPSQVIA | >IMGVR_UViG_3300032893_002930 |
| Cas12j-8 | GIHGVPAAIKPTVSQFLTPGFKLIRNHSRTAGLKLKNEGEEACKKFVRENEIPKDECPNFQGGPAIANIIAKSREFTEWEIYQSSLAIQEVIFTLPKDKLPEPILKEEWRAQWLSEHGLDTVPYKEAAGLNLIIKNAVNTYKGVQVKVDNKNKNNLAKINRKNEIAKLNGEQEISFEEIKAFDDKGYLLQKPSPNKSIYCYQSVSPKPFITSKYHNVNLPEEYIGYYRKSNEPIVSPYQFDRLRIPIGEPGYVPKWQYTFLSKKENKRRKLSKRIKNVSPILGIICIKKDWCVFDMRGLLRTNHWKKYHKPTDSINDLFDYFTGDPVIDTKANVVRFRYKMENGIVNYKPVREKKGKELLENICDQNGSCKLATVDVGQNNPVAIGLFELKKVNGELTKTLISRHPTPIDFCNKITAYRERYDKLESSIKLDAIKQLTSEQKIEVDNYNNNFTPQNTKQIVCSKLNINPNDLPWDKMISGTHFISEKAQVSNKSEIYFTSTDKGKTKDVMKSDYKWFQDYKPKLSKEVRDALSDIEWRLRRESLEFNKLSKSREQDARQLANWISSMCDVIGIENLVKKNNFFGGSGKREPGWDNFYKPKKENRWWINAIHKALTELSQNKGKRVILLPAMRTSITCPKCKYCDSKNRNGEKFNCLKCGIELNADIDVATENLATVAITAQSMPKPTCERSGDAKKPVRARKAKAPEFHDKLAPSYTVVLREAV | Wang et al., (2023) |

**Supplementary Table S2.** Protein sequences of T5Exo-Cas12j editors

| **Protein** | **Sequence** |
| --- | --- |
| T5Exo-Cas12j-11 | MSKSWGKFIEEEEAEMASRRNLMIVDGTNLGFRFKHNNSKKPFASSYVSTIQSLAKSYSARTTIVLGDKGKSVFRLEHLPEYKGNRDEKYAQRTEEEKALDEQFFEYLKDAFELCKTTFPTFTIRGVEADDMAAYIVKLIGHLYDHVWLISTDGDWDTLLTDKVSRFSFTTRREYHLRDMYEHHNVDDVEQFISLKAIMGDLGDNIRGVEGIGAKRGYNIIREFGNVLDIIDQLPLPGKQKYIQNLNASEELLFRNLILVDLPTYCVDAIAAVGQDVLDKFTKDILEIAEQGGGGSGGGGSKLATMVMPGPTLSDVAPTNCECGAAAPSATLKLSDVIKTHFPAGRFRKDHQKTAGKKLKHEGEEACVEYLRNKVSDYPPNFKPPAKGTIVAQSRPFSEWPIVRASEAIQKYVYGLTVAELDVFSPGTSKPSHAEWFAKTGVENYGYRQVQGLNTIFQNTVNRFKGVLKKVENRNKKSLKRQEGANRRRVEEGLPEVPVTVESATDDEGRLLQPPGVNPSIYGYQGVAPRVCTDLQGFSGMSVDFAGYRRDPDAVLVESLPEGRLSIPKGERGYVPEWQRDPERNKFPLREGSRRQRKWYSNACHKPKPGRTSKYDPEALKKASAKDALLVSISIGEDWAIIDVRGLLRDARRRGFTPEEGLSLNSLLGLFTEYPVFDVQRGLITFTYKLGQVDVHSRKTVPTFRSRALLESLVAKEEIALVSVDLGQTNPASMKVSRVRAQEGALVAEPVHRMFLSDVLLGELSSYRKRMDAFEDAIRAQAFETMTPEQQAEITRVCDVSVEVARRRVCEKYSISPQDVPWGEMTGHSTFIVDAVLRKGGDESLVYFKNKEGETLKFRDLRISRMEGVRPRLTKDTRDALNKAVLDLKRAHPTFAKLAKQKLELARRCVNFIEREAKRYTQCERVVFVIEDLNVGFFHGKGKRDRGWDAFFTAKKENRWVIQALHKAFSDLGLHRGSYVIEVTPQRTSMTCPRCGHCDKGNRNGEKFVCLQCGATLHADLEVATDNIERVALTGKAMPKPPVRERSGDVQKAGTARKARKPLKPKQKTEPSVQEGSSDDGVDKSPGDASRNPVYNPSDTLSI |
| T5Exo-Cas12j-12 | MSKSWGKFIEEEEAEMASRRNLMIVDGTNLGFRFKHNNSKKPFASSYVSTIQSLAKSYSARTTIVLGDKGKSVFRLEHLPEYKGNRDEKYAQRTEEEKALDEQFFEYLKDAFELCKTTFPTFTIRGVEADDMAAYIVKLIGHLYDHVWLISTDGDWDTLLTDKVSRFSFTTRREYHLRDMYEHHNVDDVEQFISLKAIMGDLGDNIRGVEGIGAKRGYNIIREFGNVLDIIDQLPLPGKQKYIQNLNASEELLFRNLILVDLPTYCVDAIAAVGQDVLDKFTKDILEIAEQGGGGSGGGGSKLATMVMEKAGPTSPLSVLIHKNFEGCRFQIDHLKIAGRKLAREGEAAAIEYLLDKKCEGLPPNFQPPAKGNVIAQSRPFTEWAPYRASVAIQKYIYSLSVDERKVCDPGSSSDSHEKWFKQTGVQNYGYTHVQGLNLIFKHALARYDGVLKKVDNRNEKNRKKAERVNSFRREEGLPEEVFEEEKATDETGHLLQPPGVNHSIYCYQSVRPKPFNPRKPGGISLPEAYSGYSLKPQDELPIGSLDRLSIPPGQPGYVPEWQRSQLTTQKHRRKRSWYSAQKWKPRTGRTSTFDPDRLNCARAQGAILAVVRIHEDWVVFDVRGLLRNALWRELAGKGLTVRDLLDFFTGDPVVDTKRGVVTFTYKLGKVDVHSLRTVRGKRSKKVLEDLTLSSDVGLVTIDLGQTNVLAADYSKVTRSENGELLAVPLSKSFLPKHLLHEVTAYRTSYDQMEEGFRRKALLTLTEDQQVEVTLVRDFSVESSKTKLLQLGVDVTSLPWEKMSSNTTYISDQLLQQGADPASLFFDGERDGKPCRHKKKDRTWAYLVRPKVSPETRKALNEALWALKNTSPEFESLSKRKIQFSRRCMNYLLNEAKRISGCGQVVFVIEDLNVRVHHGRGKRAIGWDNFFKPKRENRWFMQALHKAASELAIHRGMHIIEACPARSSITCPKCGHCDPENRCSSDREKFLCVKCGAAFHADLEVATFNLRKVALTGTALPKSIDHSRDGLIPKGARNRKLKEPQANDEKACA |
| T5Exo-Cas12j-13 | MSKSWGKFIEEEEAEMASRRNLMIVDGTNLGFRFKHNNSKKPFASSYVSTIQSLAKSYSARTTIVLGDKGKSVFRLEHLPEYKGNRDEKYAQRTEEEKALDEQFFEYLKDAFELCKTTFPTFTIRGVEADDMAAYIVKLIGHLYDHVWLISTDGDWDTLLTDKVSRFSFTTRREYHLRDMYEHHNVDDVEQFISLKAIMGDLGDNIRGVEGIGAKRGYNIIREFGNVLDIIDQLPLPGKQKYIQNLNASEELLFRNLILVDLPTYCVDAIAAVGQDVLDKFTKDILEIAEQGGGGSGGGGSKLATMVMKKPNNIRRIREEHFEGLCFGKDVLTKAGKIYEKDGEEAAIDFLMGKDEEDPPNFKPPAKTTIVAQSRPFDQWPIYQVSQAVQERVFAYTEEEFNASKEALFSGDISSKSRDFWFKTNNISDQGIGAQGLNTILSHAFSRYSGVIKKVENRNKKRLKKLSKKNQLKIEEGLEILEFKPDSAFNENGLLAQPPGINPNIYGYQAVTPFVFDPDNPGDVILPKQYEGYSRKPDDIIEKGPSRLDIPKGQPGYVPEHQRKNLKKKGRVRLYRRTPPKTKALASILAVLQIGKDWVLFDMRGLLRSVYMREAATPGQISAKDLLDTFTGCPVLNTRTGEFTFCYKLRSEGALHARKIYTKGETRTLLTSLTSENNTIALVTVDLGQRNPAAIMISRLSRKEELSEKDIQPVSRRLLPDRYLNELKRYRDAYDAFRQEVRDEAFTSLCPEHQEQVQQYEALTPEKAKNLVLKHFFGTHDPDLPWDDMTSNTHYIANLYLERGGDPSKVFFTRPLKKDSKSKKPRKPTKRTDASISRLPEIRPKMPEDARKAFEKAKWEIYTGHEKFPKLAKRVNQLCREIANWIEKEAKRLTLCDTVVVGIEDLSLPPKRGKGKFQETWQGFFRQKFENRWVIDTLKKAIQNRAHDKGKYVLGLAPYWTSQRCPACGFIHKSNRNGDHFKCLKCEALFHADSEVATWNLALVAVLGKGITNPDSKKPSGQKKTGTTRKKQIKGKNKGKETVNVPPTTQEVEDIIAFFEKDDETVRNPVYKPTGT |
| T5ExoCas12j-14 | MSKSWGKFIEEEEAEMASRRNLMIVDGTNLGFRFKHNNSKKPFASSYVSTIQSLAKSYSARTTIVLGDKGKSVFRLEHLPEYKGNRDEKYAQRTEEEKALDEQFFEYLKDAFELCKTTFPTFTIRGVEADDMAAYIVKLIGHLYDHVWLISTDGDWDTLLTDKVSRFSFTTRREYHLRDMYEHHNVDDVEQFISLKAIMGDLGDNIRGVEGIGAKRGYNIIREFGNVLDIIDQLPLPGKQKYIQNLNASEELLFRNLILVDLPTYCVDAIAAVGQDVLDKFTKDILEIAEQGGGGSGGGGSKLATMVMPDKKETPLVALCKKSFPGLRFKKHDSRQAGRILKSKGEGAAVAFLEGKGGTTQPNFKPPVKCNIVAMSRPLEEWPIYKASVVIQKYVYAQSYEEFKATDPGKSEAGLRAWLKATRVDTDGYFNVQGLNLIFQNARATYEGVLKKVENRNSKKVAKIEQRNEHRAERGLPLLTLDEPETALDETGHLRHRPGINCSVFGYQHMKLKPYVPGSIPGVTGYSRDPSTPIAACGVDRLEIPEGQPGYVPPWDRENLSVKKHRRKRASWARSRGGAIDDNMLLAVVRVADDWALLDLRGLLRNTQYRKLLDRSVPVTIESLLNLVTNDPTLSVVKKPGKPVRYTATLIYKQGVVPVVKAKVVKGSYVSKMLDDTTETFSLVGVDLGVNNLIAANALRIRPGKCVERLQAFTLPEQTVEDFFRFRKAYDKHQENLRLAAVRSLTAEQQAEVLALDTFGPEQAKMQVCGHLGLSVDEVPWDKVNSRSSILSDLAKERGVDDTLYMFPFFKGKGKKRKTEIRKRWDVNWAQHFRPQLTSETRKALNEAKWEAERNSSKYHQLSIRKKELSRHCVNYVIRTAEKRAQCGKVIVAVEDLHHSFRRGGKGSRKSGWGGFFAAKQEGRWLMDALFGAFCDLAVHRGYRVIKVDPYNTSRTCPECGHCDKANRDRVNREAFICVCCGYRGNADIDVAAYNIAMVAITGVSLRKAARASVASTPLESLAAE |
| T5Exo-Cas12j-15 | MSKSWGKFIEEEEAEMASRRNLMIVDGTNLGFRFKHNNSKKPFASSYVSTIQSLAKSYSARTTIVLGDKGKSVFRLEHLPEYKGNRDEKYAQRTEEEKALDEQFFEYLKDAFELCKTTFPTFTIRGVEADDMAAYIVKLIGHLYDHVWLISTDGDWDTLLTDKVSRFSFTTRREYHLRDMYEHHNVDDVEQFISLKAIMGDLGDNIRGVEGIGAKRGYNIIREFGNVLDIIDQLPLPGKQKYIQNLNASEELLFRNLILVDLPTYCVDAIAAVGQDVLDKFTKDILEIAEQGGGGSGGGGSKLATMVMTPSPQIARLVETPLAAALKAHHPGKKFRSDYLKKAGKILKDQGVEAAMAHLDGKDQAEPPNFKPPAKCRIVARSREFSEWPIVKASVEIQKYIYGLTLEERKACDPGKSSASHKAWFAKTGVNTFGYSSVQGFNLIFGHTLGRYDGVLVKTENLNKKRAEKNERFRAKALAEGRAEPVCPPLVTATNDTGQDVTLEDGRVVRPGQLLQPPGINPNIYAYQQVSPKAYVPGIIELPEEFQGYSRDPNAVILPLVPRDRLSIPKGQPGYVPEPHREGLTGRKDRRMRRYYETERGTKLKRPPLTAKGRADKANEALLVVVRIDSDWVVMDVRGLLRNARWRRLVSKEGITLNGLLDLFTGDPVLNPKDCSVSRDTGDPVNDPRHGVVTFCYKLGVVDVCSKDRPIKGFRTKEVLERLTSSGTVGMVSIDLGQTNPVAAAVSRVTKGLQAETLETFTLPDDLLGKVRAYRAKTDRMEEGFRRNALRKLTAEQQAEITRYNDATEQQAKALVCSTYGIGPEEVPWERMTSNTTYISDHILDHGGDPDTVFFMATKRGQNKPTLHKRKDKAWGQKFRPAISVETRLARQAAEWELRRASLEFQKLSVWKTELCRQAVNYVMERTKKRTQCDVIIPVIEDLPVPLFHGSGKRDPGWANFFVHKRENRWFIDGLHKAFSELGKHRGIYVFEVCPQRTSITCPKCGHCDPDNRDGEKFVCLSCQATLHADLDVATTNLVRVALTGKVMPRSERSGDAQTPGPARKARTGKIKGSKPTSAPQGATQTDAKAHLSQTGV |
| T5Exo-Cas12j-16 | MSKSWGKFIEEEEAEMASRRNLMIVDGTNLGFRFKHNNSKKPFASSYVSTIQSLAKSYSARTTIVLGDKGKSVFRLEHLPEYKGNRDEKYAQRTEEEKALDEQFFEYLKDAFELCKTTFPTFTIRGVEADDMAAYIVKLIGHLYDHVWLISTDGDWDTLLTDKVSRFSFTTRREYHLRDMYEHHNVDDVEQFISLKAIMGDLGDNIRGVEGIGAKRGYNIIREFGNVLDIIDQLPLPGKQKYIQNLNASEELLFRNLILVDLPTYCVDAIAAVGQDVLDKFTKDILEIAEQGGGGSGGGGSKLATMVMGCYNAGAMKKTTNLKIENGVSPLAQMTRKHFPGKRFPASVLKPAGRKLKDHGEQAAIEFLQANIDVPYGNFKAPAKCNVVATSRPYSEWPLYKFSSELQKAVFALSKDRLMEIEPSKQSDAENEKFLSAIGVSADPSINVTFVSACISKAVHTYLGMEKKAENKYQKKLSRCRSESELSSVTPENVYNEDGTLSDGWRPGFNANLYGNSNSKLSLFGSVRSHNKVELPVWLSEYREWAKNRSKDSKINEYSASVDRLSIPEGQPGHVPLWQRDSSKRTKPGGGEIKEGVRRHRWYSNRNNANRRNKVDQATRLAASAMEMVLAIAFFGEDWVLFDIRGLLRNARYRKLVNKNTTYGDLMELFTADPVLDTKRGIITASYKDTTLKIVQQTIVGEKKSKSKILEEVQKNGPVAVVSIDLGVNEPVSYRVSRVDVAGGNAIVAELAAEGFMSNELKKEISSYREKSDELNGDLREKAVLSLSDEMQAEIRRVDATNASDSKNRICEMLSLDPESVDWSKMTTQTRFIFNKHVENGGDPNVLLFTPTEDKKNKGKKSKNKKGEYGDRVPHSDSGIARNIAREKLSMETCEALNKAKRELQQEDPRYGKLSKRKQEFARRVVNGVVVRAQEVTGCDNVVLVVEKLNVSNKMFSGSGKRAPGWDNFFVHKKENRWFIQALHKAFTDKAAHKGIPVIEIKPSYTSQTCPACEHCDKDNRDGVHFCCTRCGFTGHADLNVACFNIEKVALTGEAMSGPGSATAHKKTRKPKKAMVESDKAA |
| T5Exo-Cas12j-17 | MSKSWGKFIEEEEAEMASRRNLMIVDGTNLGFRFKHNNSKKPFASSYVSTIQSLAKSYSARTTIVLGDKGKSVFRLEHLPEYKGNRDEKYAQRTEEEKALDEQFFEYLKDAFELCKTTFPTFTIRGVEADDMAAYIVKLIGHLYDHVWLISTDGDWDTLLTDKVSRFSFTTRREYHLRDMYEHHNVDDVEQFISLKAIMGDLGDNIRGVEGIGAKRGYNIIREFGNVLDIIDQLPLPGKQKYIQNLNASEELLFRNLILVDLPTYCVDAIAAVGQDVLDKFTKDILEIAEQGGGGSGGGGSKLATMVMSKEKTPPSAYAILKAKHFPDLDFEKKHKMMAGRMFKNGASEQEVVQYLQGKGSESLMDVKPPAKSPILAQSRPFDEWEMVRTSRLIQETIFGIPKRGSIPKRDGLSETQFNELVASLEVGGKPMLNKQTRAIFYGLLGIKPPTFHAMAQNILIDLAINIRKGVLKKVDNLNEKNRKKVKRIRDAGEQDVMVPAEVTAHDDRGYLNHPPGVNPTIPGYQGVVIPFPEGFEGLPSGMTPVDWSHVLVDYLPHDRLSIPKGSPGYIPEWQRPLLNRHKGRRHRSWYANSLNKPRKSRTEEAKDRQNAGKRTALIEAERLKGVLPVLMRFKEDWLIIDARGLLRNARYRGVLPEGSTLGNLIDLFSDSPRVDTRRGICTFLYRKGRAYSTKPVKRKESKETLLKLTEKSTIALVSIDLGQTNPLTAKLSKVRQVDGCLVAEPVLRKLIDNASEDGKEIARYRVAHDLLRARILEDAIDLLGIYKDEVVRARSDTPDLCKERVCRFLGLDSQAIDWDRMTPYTDFIAQAFVAKGGDPKVVTIKPNGKPKMFRKDRSIKNMKGIRLDISKEASSAYREAQWAIQRESPDFQRLAVWQSQLTKRIVNQLVAWAKKCTQCDTVVLAFEDLNIGMMHGSGKWANGGWNALFLHKQENRWFMQAFHKALTELSAHKGIPTIEVLPHRTSITCTQCGHCHPGNRDGERFKCLKCEFLANTDLEIATDNIERVALTGLPMPKGERSSAKRKPGGTRKTKKSKHSGNSPLAAE |
| T5Exo-Cas12j-18 | MSKSWGKFIEEEEAEMASRRNLMIVDGTNLGFRFKHNNSKKPFASSYVSTIQSLAKSYSARTTIVLGDKGKSVFRLEHLPEYKGNRDEKYAQRTEEEKALDEQFFEYLKDAFELCKTTFPTFTIRGVEADDMAAYIVKLIGHLYDHVWLISTDGDWDTLLTDKVSRFSFTTRREYHLRDMYEHHNVDDVEQFISLKAIMGDLGDNIRGVEGIGAKRGYNIIREFGNVLDIIDQLPLPGKQKYIQNLNASEELLFRNLILVDLPTYCVDAIAAVGQDVLDKFTKDILEIAEQGGGGSGGGGSKLATMVMLPPSNKIGKSMSLKEFINKRNFKSSIIKQAGKILKKEGEEAVKKYLDDNYVEGYKKRDFPITAKCNIVASNRKIEDFDISKFSSFIQNYVFNLNKDNFEEFSKIKYNRKSFDELYKKIANEIGLEKPNYENIQGEIAVIRNAINIYNGVLKKVENRNKKIQEKNQSKDPPKLLSAFDDNGFLAERPGINETIYGYQSVRLRHLDVEKDKDIIVQLPDIYQKYNKKSTDKISVKKRLNKYNVDEYGKLISKRRKERINKDDAILCVSNFGDDWIIFDARGLLRQTYRYKLKKKGLCIKDLLNLFTGDPIINPTKTDLKEALSLSFKDGIINNRTLKVKNYKKCPELISELIRDKGKVAMISIDLGQTNPISYRLSKFTANNVAYIENGVISEDDIVKMKKWREKSDKLENLIKEEAIASLSDDEQREVRLYENDIADNTKKKILEKFNIREEDLDFSKMSNNTYFIRDCLKNKNIDESEFTFEKNGKKLDPTDACFAREYKNKLSELTRKKINEKIWEIKKNSKEYHKISIYKKETIRYIVNKLIKQSKEKSECDDIIVNIEKLQIGGNFFGGRGKRDPGWNNFFLPKEENRWFINACHKAFSELAPHKGIIVIESDPAYTSQTCPKCENCDKENRNGEKFKCKKCNYEANADIDVATENLEKIAKNGRRLIKNFDQLGERLPGAEMPGGARKRKPSKSLPKNGRGAGVGSEPELINQSPSQVIA |

**Supplementary Table S3.** Protein sequences of Be-(d)Cas12j editors

| **Protein** | **Sequence** |
| --- | --- |
| Be-(d)Cas12j-11 | MSEVEFSHEYWMRHALTLAKRARDEREVPVGAVLVLNNRVIGEGWNRAIGLHDPTAHAEIMALRQGGLVMQNYRLIDATLYVTFEPCVMCAGAMIHSRIGRVVFGVRNSKRGAAGSLMNVLNYPGMNHRVEITEGILADECAALLCDFYRMPRQVFNAQKKAQSSINSGGSSGGSSGSETPGTSESATPESSGGSSGGSMPGPTLSDVAPTNCECGAAAPSATLKLSDVIKTHFPAGRFRKDHQKTAGKKLKHEGEEACVEYLRNKVSDYPPNFKPPAKGTIVAQSRPFSEWPIVRASEAIQKYVYGLTVAELDVFSPGTSKPSHAEWFAKTGVENYGYRQVQGLNTIFQNTVNRFKGVLKKVENRNKKSLKRQEGANRRRVEEGLPEVPVTVESATDDEGRLLQPPGVNPSIYGYQGVAPRVCTDLQGFSGMSVDFAGYRRDPDAVLVESLPEGRLSIPKGERGYVPEWQRDPERNKFPLREGSRRQRKWYSNACHKPKPGRTSKYDPEALKKASAKDALLVSISIGEDWAIIDVRGLLRDARRRGFTPEEGLSLNSLLGLFTEYPVFDVQRGLITFTYKLGQVDVHSRKTVPTFRSRALLESLVAKEEIALVSVDLGQTNPASMKVSRVRAQEGALVAEPVHRMFLSDVLLGELSSYRKRMDAFEDAIRAQAFETMTPEQQAEITRVCDVSVEVARRRVCEKYSISPQDVPWGEMTGHSTFIVDAVLRKGGDESLVYFKNKEGETLKFRDLRISRMEGVRPRLTKDTRDALNKAVLDLKRAHPTFAKLAKQKLELARRCVNFIEREAKRYTQCERVVFVIADLNVGFFHGKGKRDRGWDAFFTAKKENRWVIQALHKAFSDLGLHRGSYVIEVTPQRTSMTCPRCGHCDKGNRNGEKFVCLQCGATLHADLEVATDNIERVALTGKAMPKPPVRERSGDVQKAGTARKARKPLKPKQKTEPSVQEGSSDDGVDKSPGDASRNPVYNPSDTLSI |
| Be-(d)Cas12j-12 | MSEVEFSHEYWMRHALTLAKRARDEREVPVGAVLVLNNRVIGEGWNRAIGLHDPTAHAEIMALRQGGLVMQNYRLIDATLYVTFEPCVMCAGAMIHSRIGRVVFGVRNSKRGAAGSLMNVLNYPGMNHRVEITEGILADECAALLCDFYRMPRQVFNAQKKAQSSINSGGSSGGSSGSETPGTSESATPESSGGSSGGSMEKAGPTSPLSVLIHKNFEGCRFQIDHLKIAGRKLAREGEAAAIEYLLDKKCEGLPPNFQPPAKGNVIAQSRPFTEWAPYRASVAIQKYIYSLSVDERKVCDPGSSSDSHEKWFKQTGVQNYGYTHVQGLNLIFKHALARYDGVLKKVDNRNEKNRKKAERVNSFRREEGLPEEVFEEEKATDETGHLLQPPGVNHSIYCYQSVRPKPFNPRKPGGISLPEAYSGYSLKPQDELPIGSLDRLSIPPGQPGYVPEWQRSQLTTQKHRRKRSWYSAQKWKPRTGRTSTFDPDRLNCARAQGAILAVVRIHEDWVVFDVRGLLRNALWRELAGKGLTVRDLLDFFTGDPVVDTKRGVVTFTYKLGKVDVHSLRTVRGKRSKKVLEDLTLSSDVGLVTIDLGQTNVLAADYSKVTRSENGELLAVPLSKSFLPKHLLHEVTAYRTSYDQMEEGFRRKALLTLTEDQQVEVTLVRDFSVESSKTKLLQLGVDVTSLPWEKMSSNTTYISDQLLQQGADPASLFFDGERDGKPCRHKKKDRTWAYLVRPKVSPETRKALNEALWALKNTSPEFESLSKRKIQFSRRCMNYLLNEAKRISGCGQVVFVIADLNVRVHHGRGKRAIGWDNFFKPKRENRWFMQALHKAASELAIHRGMHIIEACPARSSITCPKCGHCDPENRCSSDREKFLCVKCGAAFHADLEVATFNLRKVALTGTALPKSIDHSRDGLIPKGARNRKLKEPQANDEKACA |
| Be-(d)Cas12j-13 | MSEVEFSHEYWMRHALTLAKRARDEREVPVGAVLVLNNRVIGEGWNRAIGLHDPTAHAEIMALRQGGLVMQNYRLIDATLYVTFEPCVMCAGAMIHSRIGRVVFGVRNSKRGAAGSLMNVLNYPGMNHRVEITEGILADECAALLCDFYRMPRQVFNAQKKAQSSINSGGSSGGSSGSETPGTSESATPESSGGSSGGSMKKPNNIRRIREEHFEGLCFGKDVLTKAGKIYEKDGEEAAIDFLMGKDEEDPPNFKPPAKTTIVAQSRPFDQWPIYQVSQAVQERVFAYTEEEFNASKEALFSGDISSKSRDFWFKTNNISDQGIGAQGLNTILSHAFSRYSGVIKKVENRNKKRLKKLSKKNQLKIEEGLEILEFKPDSAFNENGLLAQPPGINPNIYGYQAVTPFVFDPDNPGDVILPKQYEGYSRKPDDIIEKGPSRLDIPKGQPGYVPEHQRKNLKKKGRVRLYRRTPPKTKALASILAVLQIGKDWVLFDMRGLLRSVYMREAATPGQISAKDLLDTFTGCPVLNTRTGEFTFCYKLRSEGALHARKIYTKGETRTLLTSLTSENNTIALVTVDLGQRNPAAIMISRLSRKEELSEKDIQPVSRRLLPDRYLNELKRYRDAYDAFRQEVRDEAFTSLCPEHQEQVQQYEALTPEKAKNLVLKHFFGTHDPDLPWDDMTSNTHYIANLYLERGGDPSKVFFTRPLKKDSKSKKPRKPTKRTDASISRLPEIRPKMPEDARKAFEKAKWEIYTGHEKFPKLAKRVNQLCREIANWIEKEAKRLTLCDTVVVGIADLSLPPKRGKGKFQETWQGFFRQKFENRWVIDTLKKAIQNRAHDKGKYVLGLAPYWTSQRCPACGFIHKSNRNGDHFKCLKCEALFHADSEVATWNLALVAVLGKGITNPDSKKPSGQKKTGTTRKKQIKGKNKGKETVNVPPTTQEVEDIIAFFEKDDETVRNPVYKPTGT |
| Be-(d)Cas12j-14 | MSEVEFSHEYWMRHALTLAKRARDEREVPVGAVLVLNNRVIGEGWNRAIGLHDPTAHAEIMALRQGGLVMQNYRLIDATLYVTFEPCVMCAGAMIHSRIGRVVFGVRNSKRGAAGSLMNVLNYPGMNHRVEITEGILADECAALLCDFYRMPRQVFNAQKKAQSSINSGGSSGGSSGSETPGTSESATPESSGGSSGGSMPDKKETPLVALCKKSFPGLRFKKHDSRQAGRILKSKGEGAAVAFLEGKGGTTQPNFKPPVKCNIVAMSRPLEEWPIYKASVVIQKYVYAQSYEEFKATDPGKSEAGLRAWLKATRVDTDGYFNVQGLNLIFQNARATYEGVLKKVENRNSKKVAKIEQRNEHRAERGLPLLTLDEPETALDETGHLRHRPGINCSVFGYQHMKLKPYVPGSIPGVTGYSRDPSTPIAACGVDRLEIPEGQPGYVPPWDRENLSVKKHRRKRASWARSRGGAIDDNMLLAVVRVADDWALLDLRGLLRNTQYRKLLDRSVPVTIESLLNLVTNDPTLSVVKKPGKPVRYTATLIYKQGVVPVVKAKVVKGSYVSKMLDDTTETFSLVGVDLGVNNLIAANALRIRPGKCVERLQAFTLPEQTVEDFFRFRKAYDKHQENLRLAAVRSLTAEQQAEVLALDTFGPEQAKMQVCGHLGLSVDEVPWDKVNSRSSILSDLAKERGVDDTLYMFPFFKGKGKKRKTEIRKRWDVNWAQHFRPQLTSETRKALNEAKWEAERNSSKYHQLSIRKKELSRHCVNYVIRTAEKRAQCGKVIVAVADLHHSFRRGGKGSRKSGWGGFFAAKQEGRWLMDALFGAFCDLAVHRGYRVIKVDPYNTSRTCPECGHCDKANRDRVNREAFICVCCGYRGNADIDVAAYNIAMVAITGVSLRKAARASVASTPLESLAAE |
| Be-(d)Cas12j-15 | MSEVEFSHEYWMRHALTLAKRARDEREVPVGAVLVLNNRVIGEGWNRAIGLHDPTAHAEIMALRQGGLVMQNYRLIDATLYVTFEPCVMCAGAMIHSRIGRVVFGVRNSKRGAAGSLMNVLNYPGMNHRVEITEGILADECAALLCDFYRMPRQVFNAQKKAQSSINSGGSSGGSSGSETPGTSESATPESSGGSSGGSMTPSPQIARLVETPLAAALKAHHPGKKFRSDYLKKAGKILKDQGVEAAMAHLDGKDQAEPPNFKPPAKCRIVARSREFSEWPIVKASVEIQKYIYGLTLEERKACDPGKSSASHKAWFAKTGVNTFGYSSVQGFNLIFGHTLGRYDGVLVKTENLNKKRAEKNERFRAKALAEGRAEPVCPPLVTATNDTGQDVTLEDGRVVRPGQLLQPPGINPNIYAYQQVSPKAYVPGIIELPEEFQGYSRDPNAVILPLVPRDRLSIPKGQPGYVPEPHREGLTGRKDRRMRRYYETERGTKLKRPPLTAKGRADKANEALLVVVRIDSDWVVMDVRGLLRNARWRRLVSKEGITLNGLLDLFTGDPVLNPKDCSVSRDTGDPVNDPRHGVVTFCYKLGVVDVCSKDRPIKGFRTKEVLERLTSSGTVGMVSIDLGQTNPVAAAVSRVTKGLQAETLETFTLPDDLLGKVRAYRAKTDRMEEGFRRNALRKLTAEQQAEITRYNDATEQQAKALVCSTYGIGPEEVPWERMTSNTTYISDHILDHGGDPDTVFFMATKRGQNKPTLHKRKDKAWGQKFRPAISVETRLARQAAEWELRRASLEFQKLSVWKTELCRQAVNYVMERTKKRTQCDVIIPVIADLPVPLFHGSGKRDPGWANFFVHKRENRWFIDGLHKAFSELGKHRGIYVFEVCPQRTSITCPKCGHCDPDNRDGEKFVCLSCQATLHADLDVATTNLVRVALTGKVMPRSERSGDAQTPGPARKARTGKIKGSKPTSAPQGATQTDAKAHLSQTGV |
| Be-(d)Cas12j-16 | MSEVEFSHEYWMRHALTLAKRARDEREVPVGAVLVLNNRVIGEGWNRAIGLHDPTAHAEIMALRQGGLVMQNYRLIDATLYVTFEPCVMCAGAMIHSRIGRVVFGVRNSKRGAAGSLMNVLNYPGMNHRVEITEGILADECAALLCDFYRMPRQVFNAQKKAQSSINSGGSSGGSSGSETPGTSESATPESSGGSSGGSMGCYNAGAMKKTTNLKIENGVSPLAQMTRKHFPGKRFPASVLKPAGRKLKDHGEQAAIEFLQANIDVPYGNFKAPAKCNVVATSRPYSEWPLYKFSSELQKAVFALSKDRLMEIEPSKQSDAENEKFLSAIGVSADPSINVTFVSACISKAVHTYLGMEKKAENKYQKKLSRCRSESELSSVTPENVYNEDGTLSDGWRPGFNANLYGNSNSKLSLFGSVRSHNKVELPVWLSEYREWAKNRSKDSKINEYSASVDRLSIPEGQPGHVPLWQRDSSKRTKPGGGEIKEGVRRHRWYSNRNNANRRNKVDQATRLAASAMEMVLAIAFFGEDWVLFDIRGLLRNARYRKLVNKNTTYGDLMELFTADPVLDTKRGIITASYKDTTLKIVQQTIVGEKKSKSKILEEVQKNGPVAVVSIDLGVNEPVSYRVSRVDVAGGNAIVAELAAEGFMSNELKKEISSYREKSDELNGDLREKAVLSLSDEMQAEIRRVDATNASDSKNRICEMLSLDPESVDWSKMTTQTRFIFNKHVENGGDPNVLLFTPTEDKKNKGKKSKNKKGEYGDRVPHSDSGIARNIAREKLSMETCEALNKAKRELQQEDPRYGKLSKRKQEFARRVVNGVVVRAQEVTGCDNVVLVVAKLNVSNKMFSGSGKRAPGWDNFFVHKKENRWFIQALHKAFTDKAAHKGIPVIEIKPSYTSQTCPACEHCDKDNRDGVHFCCTRCGFTGHADLNVACFNIEKVALTGEAMSGPGSATAHKKTRKPKKAMVESDKAA |
| Be-(d)Cas12j-17 | MSEVEFSHEYWMRHALTLAKRARDEREVPVGAVLVLNNRVIGEGWNRAIGLHDPTAHAEIMALRQGGLVMQNYRLIDATLYVTFEPCVMCAGAMIHSRIGRVVFGVRNSKRGAAGSLMNVLNYPGMNHRVEITEGILADECAALLCDFYRMPRQVFNAQKKAQSSINSGGSSGGSSGSETPGTSESATPESSGGSSGGSMSKEKTPPSAYAILKAKHFPDLDFEKKHKMMAGRMFKNGASEQEVVQYLQGKGSESLMDVKPPAKSPILAQSRPFDEWEMVRTSRLIQETIFGIPKRGSIPKRDGLSETQFNELVASLEVGGKPMLNKQTRAIFYGLLGIKPPTFHAMAQNILIDLAINIRKGVLKKVDNLNEKNRKKVKRIRDAGEQDVMVPAEVTAHDDRGYLNHPPGVNPTIPGYQGVVIPFPEGFEGLPSGMTPVDWSHVLVDYLPHDRLSIPKGSPGYIPEWQRPLLNRHKGRRHRSWYANSLNKPRKSRTEEAKDRQNAGKRTALIEAERLKGVLPVLMRFKEDWLIIDARGLLRNARYRGVLPEGSTLGNLIDLFSDSPRVDTRRGICTFLYRKGRAYSTKPVKRKESKETLLKLTEKSTIALVSIDLGQTNPLTAKLSKVRQVDGCLVAEPVLRKLIDNASEDGKEIARYRVAHDLLRARILEDAIDLLGIYKDEVVRARSDTPDLCKERVCRFLGLDSQAIDWDRMTPYTDFIAQAFVAKGGDPKVVTIKPNGKPKMFRKDRSIKNMKGIRLDISKEASSAYREAQWAIQRESPDFQRLAVWQSQLTKRIVNQLVAWAKKCTQCDTVVLAFADLNIGMMHGSGKWANGGWNALFLHKQENRWFMQAFHKALTELSAHKGIPTIEVLPHRTSITCTQCGHCHPGNRDGERFKCLKCEFLANTDLEIATDNIERVALTGLPMPKGERSSAKRKPGGTRKTKKSKHSGNSPLAAE |
| Be-(d)Cas12j-18 | MSEVEFSHEYWMRHALTLAKRARDEREVPVGAVLVLNNRVIGEGWNRAIGLHDPTAHAEIMALRQGGLVMQNYRLIDATLYVTFEPCVMCAGAMIHSRIGRVVFGVRNSKRGAAGSLMNVLNYPGMNHRVEITEGILADECAALLCDFYRMPRQVFNAQKKAQSSINSGGSSGGSSGSETPGTSESATPESSGGSSGGSMLPPSNKIGKSMSLKEFINKRNFKSSIIKQAGKILKKEGEEAVKKYLDDNYVEGYKKRDFPITAKCNIVASNRKIEDFDISKFSSFIQNYVFNLNKDNFEEFSKIKYNRKSFDELYKKIANEIGLEKPNYENIQGEIAVIRNAINIYNGVLKKVENRNKKIQEKNQSKDPPKLLSAFDDNGFLAERPGINETIYGYQSVRLRHLDVEKDKDIIVQLPDIYQKYNKKSTDKISVKKRLNKYNVDEYGKLISKRRKERINKDDAILCVSNFGDDWIIFDARGLLRQTYRYKLKKKGLCIKDLLNLFTGDPIINPTKTDLKEALSLSFKDGIINNRTLKVKNYKKCPELISELIRDKGKVAMISIDLGQTNPISYRLSKFTANNVAYIENGVISEDDIVKMKKWREKSDKLENLIKEEAIASLSDDEQREVRLYENDIADNTKKKILEKFNIREEDLDFSKMSNNTYFIRDCLKNKNIDESEFTFEKNGKKLDPTDACFAREYKNKLSELTRKKINEKIWEIKKNSKEYHKISIYKKETIRYIVNKLIKQSKEKSECDDIIVNIAKLQIGGNFFGGRGKRDPGWNNFFLPKEENRWFINACHKAFSELAPHKGIIVIESDPAYTSQTCPKCENCDKENRNGEKFKCKKCNYEANADIDVATENLEKIAKNGRRLIKNFDQLGERLPGAEMPGGARKRKPSKSLPKNGRGAGVGSEPELINQSPSQVIA |

**Supplementary Table S4.** crRNA sequences for respective proteins

| **crRNA (5′–3′)** | **Corresponding proteins** | **Notes** |
| --- | --- | --- |
| CUAGGAACGCACGCAGAUUGCUCGGUACGCCGAGAC | Cas12j-11, T5Exo-Cas12j-11, Be-(d)Cas12j-11 | 36-nt pre-processed crRNA |
| GUUGAACCUAGAUCAGAUGGCUCAGUACGCUGAGAC | Cas12j-12, T5Exo-Cas12j-12, Be-(d)Cas12j-12 |  |
| GCUGGAAGACUCAAUGAUGGCUCCUUACGAGGAGAC | Cas12j-13, T5Exo-Cas12j-13, Be-(d)Cas12j-13 |  |
| CUGGGGACCGAUCCUGAUUGCUCGCUGCGGCGAGAC | Cas12j-14, T5Exo-Cas12j-14, Be-(d)Cas12j-14 |  |
| GGUUGAACCCUCAACAGAUUGCUCGGUAAGCCGAGAC | Cas12j-15, T5Exo-Cas12j-15, Be-(d)Cas12j-15 |  |
| AUAGAAACCCCUACAAAUUGCGCUCUGAGGAGCGAC | Cas12j-16, T5Exo-Cas12j-16, Be-(d)Cas12j-16 |  |
| GUUCGGCGAUCCUUUGAUUGCUCAGUACGCUGAGAC | Cas12j-17, T5Exo-Cas12j-17, Be-(d)Cas12j-17 |  |
| GUCGCAAGACUCGAAUAAUUGCCCCUCUAUGGGGAC | Cas12j-18, T5Exo-Cas12j-18, Be-(d)Cas12j-18 |  |
| GAUCAGAUGGCUCAGUACGCUGAGAC | Cas12j-12, T5Exo-Cas12j-12 | 26-nt mature crRNA |
| UCAAUGAUGGCUCCUUACGAGGAGAC | Cas12j-13, T5Exo-Cas12j-13 |  |
| CCUUUGAUUGCUCAGUACGCUGAGAC | Cas12j-17, T5Exo-Cas12j-17 |  |
| UCGAAUAAUUGCCCCUCUAUGGGGAC | Cas12j-18, T5Exo-Cas12j-18 |  |
| CUUUCAAGACUAAUAGAUUGCUCCUUACGAGGAGAC | Cas12j-8 |  |
| UAAUUUCUACUAAGUGUAGAU | LbCas12a |  |

**Supplementary Table S5.** Guide RNAs used in this study

| **S. No.** | **Target name** | **Guide RNA sequence (5′–3′)** | **Tested in** |
| --- | --- | --- | --- |
| 1 | *CLTA* target-1 | UACAAACAACCCUUCGCUGA | HEK293T cells |
| 2 | *CLTA* target-2 | UGUUAAGGAAAGUAAUGGUC |  |
| 3 | *CLTA* target-3 | GGCCUGCUUGCUAGACUUGG |  |
| 4 | *CLTA* target-4 | GGACCUCUUGCUGUCUAGGG |  |
| 5 | *CLTA* target-5 | GAUGUUUCUGCUUCUCAAUG |  |
| 6 | *CLTA* target-6 | UAGAAAGCUUCAUCUGCCAC |  |
| 7 | *CLTA* target-7 | UACUGUAUAGUCAUGCUUCU |  |
| 8 | *CLTA* target-8 | UACAUACGUGAGUUGUUAUA |  |
| 9 | *CLTA* target-9 | UACCCCUGUGUUCUGCAAUG |  |
| 10 | *CLTA* target-12 | ACAUACAUUAUGUUGAUUCA |  |
| 11 | *EMX* target-1 | UACCCCUGGGUCCUGCGGAA |  |
| 12 | *EMX* target-2 | AUGACACGGGCAUCCAGCUC |  |
| 13 | *HBB* target-1 | UAUUGGUCUCCUUAAACCUG |  |
| 14 | *HBB* target-2 | AGGAGACCAAUAGAAACUGG |  |
| 15 | *HBB* target-3 | AGGUUGCUAGUGAACACAGU |  |
| 16 | *HBB* target-4 | AUGCCCAGCCCUGGCUCCUG |  |
| 17 | *AIFM* target-1 | UACAUUGGUAAGACAAUGAA |  |
| 18 | *AIFM* target-2 | UACCAAGAGAAGACACUCAC |  |
| 19 | *Caspase-3* target-1 | UACUAUGGUCUAGCAAUAAU |  |
| 20 | *Caspase-3* target-2 | UACGACUAAAAUGCAAUGCC |  |
| 21 | *CLTA* target-1a | AAACAACCCUUCGCUGACGU |  |
| 22 | *CLTA* target-1b | AAACAACCCUUCGCUGA |  |
| 23 | *CLTA* target-9a | CCCUGUGUUCUGCAAUGAUG |  |
| 24 | *CLTA* target-9b | CCCUGUGUUCUGCAAUG |  |
| 25 | pJET1.2 plasmid | CGCUCACUGCCAAUUGCUUUCCAG | *In vitro* |

**Supplementary Table S6.** Primers used in this study

| **Primer name** | **Primer sequence (5′–3′)** | **Targets amplified** | **Notes** |
| --- | --- | --- | --- |
| *CLTA* - F | GGCAGTGCTTGCTCTGTCGTG | *CLTA* target-1, 4, 5, 6, 1a, 1b | To generate PCR amplicon for T7EI and deep sequencing studies |
| *CLTA* - R | GCAGGGAGTCACAAATCTGGAAG | *CLTA* target-1, 4, 5, 6, 1a, 1b |  |
| *CLTA* target -2 - F | CGCAGGCACCTGTAATCTCA | *CLTA* target -2 | To generate PCR amplicon for T7EI assay |
| *CLTA* target -2 - R | CCAGCTCAACTACTCCACTGA | *CLTA* target -2 |  |
| *CLTA* target -3 - F | GCCAGGGGCTGTTATCTTGGG | *CLTA* target -3 |  |
| *CLTA* target -3 - R | AGGTTTGGTCCAAAAGAACTCA | *CLTA* target -3 |  |
| *CLTA* target -7 - F | AGTGTATCCCACCCAGACAGTGT | *CLTA* target -7 | To generate PCR amplicon for T7EI and deep sequencing studies |
| *CLTA* target -7 - R | CTTTGGCTCACCCCAGCAACAC | *CLTA* target -7 |  |
| *CLTA* target -8 - F | TTTGGTAATCACTGTGTGCTATTT | *CLTA* target -8 |  |
| *CLTA* target -8 - R | ATGGTTCGAGTGATGCGGAA | *CLTA* target -8 |  |
| *CLTA* target -9 - F | GGATTGGTCACTGGCTGGAAC | *CLTA* target -9 |  |
| *CLTA* target -9 - R | TCTCCACGTTACAACCCTCA | *CLTA* target -9 |  |
| *CLTA* target -9 - F2 | TAGCCAGGCAAAACCCTGAT | *CLTA* target -9a, 9b | To generate PCR amplicon for T7EI assay |
| *CLTA* target -9 - R2 | GCCTGAAACCCAGTGATGAGA | *CLTA* target -9a, 9b |  |
| *CLTA* target-12 - F | TAGCGCATCTGGTGTCTGGC | *CLTA* target-12 | To generate PCR amplicon for T7EI and deep sequencing studies |
| *CLTA* target-12 - R | CCAAGGCTCCTTCCTAACCC | *CLTA* target-12 |  |
| *EMX* target-1 - F | CCATGAACCACCCCGCGCTGACC | *EMX* target - 1 |  |
| *EMX* target-1 - R | CGCCTGGGCTTGCGTCCGAACTG | *EMX* target - 1 |  |
| *EMX* target-2 - F | TCCGTGTCTCCAATCTCCCT | *EMX* target - 2 | To generate PCR amplicon for T7EI assay |
| *EMX* target-2 - R | GCCTGATTCCCACCTCTCAA | *EMX* target - 2 |  |
| *HBB* - F | AGGGTAGACCACCAGCAGCCTAAG | *HBB* target - 1, 2, 3, 4 | To generate PCR amplicon for deep sequencing studies |
| *HBB* - R | CTAAGCCAGTGCCAGAAGAGCCAAGG | *HBB* target - 1, 2, 3, 4 |  |
| *AIFM* target - 1 - F | CGATCAGCTTAGCAGGTCACA | *AIFM* target - 1 | To generate PCR amplicon for T7EI and deep sequencing studies |
| *AIFM* target - 1 - R | TGCTCCAGGATTGCAGAATGT | *AIFM* target - 1 |  |
| *AIFM* target - 2 - F | AGCTGGATGTGAGAGACAACA | *AIFM* target - 2 |  |
| *AIFM* target - 2 - R | TTGCAACTGGCAATGAAGCC | *AIFM* target - 2 |  |
| *Caspase-3* target - 1 - F | TGCAGCAAACCTCAGGGAAA | *Caspase-3* target - 1 |  |
| *Caspase-3* target - 1 - R | TTCACCATGGCTCAGAAGCA | *Caspase-3* target - 1 |  |
| *Caspase-3* target - 2 - F | AGGCCTAGTAGGGTGTGTGA | *Caspase-3* target - 2 |  |
| *Caspase-3* target - 2 - R | TCAGGGCAGCCGAGAATAAC | *Caspase-3* target - 2 |  |
| *CLTA*_adapter-F | TCGTCGGCAGCGTCAGATGTGTATAAGAGACAGGGCAGTGCTTGCTCTGTCGTG | *CLTA* target-1, 4, 5, 6 | Adapter linked gene specific primers used for deep amplicon sequencing  Adapter linked gene specific primers used for deep amplicon sequencing  Adapter linked gene specific primers used for deep amplicon sequencing |
| *CLTA*_adapter-R | GTCTCGTGGGCTCGGAGATGTGTATAAGAGACAGGCAGGGAGTCACAAATCTGGAAG | *CLTA* target-1, 4, 5, 6 |  |
| *CLTA* target -7 _adapter - F | TCGTCGGCAGCGTCAGATGTGTATAAGAGACAGAGTGTATCCCACCCAGACAGTGT | *CLTA* target -7 |  |
| *CLTA* target -7 _adapter - R | GTCTCGTGGGCTCGGAGATGTGTATAAGAGACAGCTTTGGCTCACCCCAGCAACAC | *CLTA* target -7 |  |
| *CLTA* target -8 _adapter - F | TCGTCGGCAGCGTCAGATGTGTATAAGAGACAGTTTGGTAATCACTGTGTGCTATTT | *CLTA* target -8 |  |
| *CLTA* target -8 _adapter - R | GTCTCGTGGGCTCGGAGATGTGTATAAGAGACAGATGGTTCGAGTGATGCGGAA | *CLTA* target -8 |  |
| *CLTA* target -9 _adapter - F | TCGTCGGCAGCGTCAGATGTGTATAAGAGACAGGGATTGGTCACTGGCTGGAA | *CLTA* target -9 |  |
| *CLTA* target -9 _adapter - R | GTCTCGTGGGCTCGGAGATGTGTATAAGAGACAGTCTCCACGTTACAACCCTCA | *CLTA* target -9 |  |
| *CLTA* target-12_adapter - F | TCGTCGGCAGCGTCAGATGTGTATAAGAGACAGTAGCGCATCTGGTGTCTGGC | *CLTA* target-12 |  |
| *CLTA* target-12_adapter - R | GTCTCGTGGGCTCGGAGATGTGTATAAGAGACAGCCAAGGCTCCTTCCTAACCC | *CLTA* target-12 |  |
| *EMX*_adapter-F | TCGTCGGCAGCGTCAGATGTGTATAAGAGACAGCCATGAACCACCCCGCGCTGACC | *EMX* target - 1 |  |
| *EMX*_adapter-R | GTCTCGTGGGCTCGGAGATGTGTATAAGAGACAGCGCCTGGGCTTGCGTCCGAACTG | *EMX* target - 1 |  |
| *HBB*_adapter - F | TCGTCGGCAGCGTCAGATGTGTATAAGAGACAGAGGGTAGACCACCAGCAGCCTAAG | *HBB* target - 1, 2, 3, 4 |  |
| *HBB*_adapter - R | GTCTCGTGGGCTCGGAGATGTGTATAAGAGACAGCTAAGCCAGTGCCAGAAGAGCCAAGG | *HBB* target - 1, 2, 3, 4 |  |
| *AIFM* target - 1_adapter - F | TCGTCGGCAGCGTCAGATGTGTATAAGAGACAGCGATCAGCTTAGCAGGTCACA | *AIFM* target - 1 |  |
| *AIFM* target - 1_adapter - R | GTCTCGTGGGCTCGGAGATGTGTATAAGAGACAGTGCTCCAGGATTGCAGAATGT | *AIFM* target - 1 |  |
| *AIFM* target - 2_adapter - F | TCGTCGGCAGCGTCAGATGTGTATAAGAGACAGAGCTGGATGTGAGAGACAACA | *AIFM* target - 2 |  |
| *AIFM* target - 2_adapter - R | GTCTCGTGGGCTCGGAGATGTGTATAAGAGACAGTTGCAACTGGCAATGAAGCC | *AIFM* target - 2 |  |
| *Caspase-3* target - 1_adapter - F | TCGTCGGCAGCGTCAGATGTGTATAAGAGACAGTGCAGCAAACCTCAGGGAAA | *Caspase-3* target - 1 |  |
| *Caspase-3* target - 1_adapter - R | GTCTCGTGGGCTCGGAGATGTGTATAAGAGACAGTTCACCATGGCTCAGAAGCA | *Caspase-3* target - 1 |  |
| *Caspase-3* target - 2_adapter - F | TCGTCGGCAGCGTCAGATGTGTATAAGAGACAGAGGCCTAGTAGGGTGTGTGA | *Caspase-3* target - 2 |  |
| *Caspase-3* target - 2_adapter - R | GTCTCGTGGGCTCGGAGATGTGTATAAGAGACAGTCAGGGCAGCCGAGAATAAC | *Caspase-3* target - 2 |  |
| crRNA forward | CGGTACCAAGGTCGGGCAGG |  | For U6-crRNA amplification |
| crRNA reverse | GAGGTACCTCGAGCGGCCC |  |  |
| *CLTA*_T7 OFF T1_F for 1st PCR | GCAGAGCAATTACGGCAAGAAGA |  | To generate off-target PCR amplicons for deep sequencing studies |
| *CLTA*_T7 OFF T1_R for 1st PCR | CTGAAGTCCAGAAATGAGATGCAA |  |  |
| *CLTA*_T9 OFF T1_F | CATTGTCTTGCTGATGTCCAGACG |  |  |
| *CLTA*_T9 OFF T1_R | TTTTATCCTTTCCCCTCCCCAGCA |  |  |
| *CLTA*_T9 OFF T2_F | CATGTAAAACACATGCCCGGTGA |  |  |
| *CLTA*_T9 OFF T2_R | ACAGGAAGAGTCTCGTACGCTC |  |  |
| *CLTA*_T9 OFF T3_F | TGTTCTTGCAGAGACAGACTGA |  |  |
| *CLTA*_T9 OFF T3_R | GCCTGCTCCCACCTCAAGAATA |  |  |
| *AIFM*_T1 OFF T1_F | TTAAAGTATACGCACGCACACACA |  |  |
| *AIFM*_T1 OFF T1_R | AGGCTCAGGACATTTAGCTTGT |  |  |
| *AIFM*_T1 OFF T2_F | ACACCCTTTGACAGCAATAGCCT |  |  |
| *AIFM*_T1 OFF T2_R | TTTGAAGTTTGTGGATGACAACGTA |  |  |
| *AIFM*_T1 OFF T3_F | TTTGGCATTGTGATAAGTCTTGAA |  |  |
| *AIFM*_T1 OFF T3_R | TCATCCATCCATTGTCTGGTTCT |  |  |
| *AIFM*_T1 OFF T4_F | TTTGGCCATCAGGCTATTGGAAG |  | To generate off-target PCR amplicons for deep sequencing studies |
| *AIFM*_T1 OFF T4_R | CTGTTCCACTCAGCTTCCCTTCT |  |  |
| *Caspase-3*_T1 OFF T1_F | ACAGGTGTTACTCAAAAGGTATGC |  |  |
| *Caspase-3*_T1 OFF T1_R | TCCCCAATGCTGCCTAAGTATCT |  |  |
| *Caspase-3*_T1 OFF T2_F | GCTTTTTCAGTAGACCAGTAGGG |  |  |
| *Caspase-3*_T1 OFF T2_R | TTCAAGAGGTGGAAGGGAGGAA |  |  |
| *Caspase-3*_T1 OFF T3_F | GAAGCATTGAATAACTCTACTCAGG |  |  |
| *Caspase-3*_T1 OFF T3_R | TGGCTGAAATTATGGTAGTGGGA |  |  |
| *CLTA*_T7 OFF T1_F_Adapter | TCGTCGGCAGCGTCAGATGTGTATAAGAGACAGCAGTTGACCCCTAGATAGTTATGG |  | Adapter linked off-target specific primers used for deep amplicon sequencing |
| *CLTA*_T7 OFF T1_R_Adapter | GTCTCGTGGGCTCGGAGATGTGTATAAGAGACAGCATTCTGCCTGTGATAATTCCTG |  |  |
| *CLTA*_T9 OFF T1_F_Adapter | TCGTCGGCAGCGTCAGATGTGTATAAGAGACAGCATTGTCTTGCTGATGTCCAGACG |  |  |
| *CLTA*_T9 OFF T1_R_Adapter | GTCTCGTGGGCTCGGAGATGTGTATAAGAGACAGTTTTATCCTTTCCCCTCCCCAGCA |  |  |
| *CLTA*_T9OFF T2_F_Adapter | TCGTCGGCAGCGTCAGATGTGTATAAGAGACAGCATGTAAAACACATGCCCGGTGA |  |  |
| *CLTA*_T9 OFF T2_R_Adapter | GTCTCGTGGGCTCGGAGATGTGTATAAGAGACAGACAGGAAGAGTCTCGTACGCTC |  |  |
| *CLTA*_T9 OFF T3_F_Adapter | TCGTCGGCAGCGTCAGATGTGTATAAGAGACAGTGTTCTTGCAGAGACAGACTGA |  |  |
| *CLTA*_T9 OFF T3_R_Adapter | GTCTCGTGGGCTCGGAGATGTGTATAAGAGACAGGCCTGCTCCCACCTCAAGAATA |  | Adapter linked off-target specific primers used for deep amplicon sequencing |
| *AIFM*_T1 OFF T1_F_Adapter | TCGTCGGCAGCGTCAGATGTGTATAAGAGACAGTTAAAGTATACGCACGCACACACA |  |  |
| *AIFM*_T1 OFF T1_R_Adapter | GTCTCGTGGGCTCGGAGATGTGTATAAGAGACAGAGGCTCAGGACATTTAGCTTGT |  |  |
| *AIFM*_T1 OFF T2_F_Adapter | TCGTCGGCAGCGTCAGATGTGTATAAGAGACAGACACCCTTTGACAGCAATAGCCT |  |  |
| *AIFM*_T1 OFF T2_R_Adapter | GTCTCGTGGGCTCGGAGATGTGTATAAGAGACAGTTTGAAGTTTGTGGATGACAACGTA |  |  |
| *AIFM*_T1 OFF T3_F_Adapter | TCGTCGGCAGCGTCAGATGTGTATAAGAGACAGTTTGGCATTGTGATAAGTCTTGAA |  |  |
| *AIFM*_T1 OFF T3_R_Adapter | GTCTCGTGGGCTCGGAGATGTGTATAAGAGACAGTCATCCATCCATTGTCTGGTTCT |  |  |
| *AIFM*_T1 OFF T4_F_Adapter | TCGTCGGCAGCGTCAGATGTGTATAAGAGACAGTTTGGCCATCAGGCTATTGGAAG |  |  |
| *AIFM*_T1 OFF T4_R_Adapter | GTCTCGTGGGCTCGGAGATGTGTATAAGAGACAGCTGTTCCACTCAGCTTCCCTTCT |  |  |
| *Caspase-3*_T1 OFF T1_F_Adapter | TCGTCGGCAGCGTCAGATGTGTATAAGAGACAGACAGGTGTTACTCAAAAGGTATGC |  |  |
| *Caspase-3*_T1 OFF T1_R_Adapter | GTCTCGTGGGCTCGGAGATGTGTATAAGAGACAGTCCCCAATGCTGCCTAAGTATCT |  |  |
| *Caspase-3*_T1 OFF T2_F_Adapter | TCGTCGGCAGCGTCAGATGTGTATAAGAGACAGGCTTTTTCAGTAGACCAGTAGGG |  |  |
| *Caspase-3*_T1 OFF T2_R_Adapter | GTCTCGTGGGCTCGGAGATGTGTATAAGAGACAGTTCAAGAGGTGGAAGGGAGGAA |  |  |
| *Caspase-3*_T1 OFF T3_F_Adapter | TCGTCGGCAGCGTCAGATGTGTATAAGAGACAGGAAGCATTGAATAACTCTACTCAGG |  |  |
| *Caspase-3*_T1 OFF T3_R_Adapter | GTCTCGTGGGCTCGGAGATGTGTATAAGAGACAGTGGCTGAAATTATGGTAGTGGGA |  |  |

**Supplementary Table S7.** Predicted off-target sequences and the chromosome locations

| **Off-target name** | **Off-target site** | **Off-target sequence (5′–3′)** |
| --- | --- | --- |
| *CLTA* target-7_Off1 | chr6; Position:24367743 | TTTTACTGTATAtTCATGtTTCT |
| *CLTA* target-9_Off1 | chr10, position:70450631 | TTTTACCtCTGTGTcCTGCAATG |
| *CLTA* target-9_Off2 | chr9; Position: 38319378 | TTTTcCCCaTGTGTTCTGCAATc |
| *CLTA* target-9_Off3 | chr6; Position: 38976932 | TTTTACtCCTGgGTTCTGCAtTG |
| *AIFM* target-1_Off1 | chr18, position:49739948 | TTTTtCATTGGTAAGACAcTGAA |
| *AIFM* target-1_Off2 | chr7; Position: 81059274 | TTTTACAgTGGTAAGAaAATtAA |
| *AIFM* target-1_Off3 | chr2, position: 166449074 | TTTTACATTGGTAAGACAgTtAA |
| *AIFM* target-1_Off4 | chr13; Position: 60855852 | TTTTACATTGGTAAGAttcTGAA |
| *Caspase-3* target-1_Off1 | chr5, position: 109646880 | TTGTgCTAgGGTCTAGCAATAAT |
| *Caspase-3* target-1_Off2 | chr3; Position: 69251050 | TTATAaTATGGTtTAGtAATAAT |
| *Caspase-3* target-1_Off3 | chr5; Position: 12298707 | TTTTACTATGGTtTtGCAATcAT |

**References:**

Wang, Y., Qi, T., Liu, J., Yang, Y., Wang, Z., Wang, Y., Wang, T., Li, M., Li, M., Lu, D. *et al.* (2023) A highly specific CRISPR-Cas12j nuclease enables allele-specific genome editing. *Science Advances*, **9**, eabo6405.
